# Redefining accessible food resources for paleodietary reconstruction in Britain: baseline recommendations based on isotopic data synthesis

**DOI:** 10.64898/2026.06.25.734399

**Authors:** Zexuan Chen, Andrew Millard, Eva Fernandez Dominguez

**Affiliations:** Department of Archaeology, Durham University, Durham, UK

## Abstract

Paleodietary reconstruction requires isotopic data from both humans and their accessible food resources. However, ideally defined accessible food resources, namely those from the same sites and periods as the target human individuals, are not always available for all ancient individuals. The number of sites with human C/N isotopic data far exceeds that with food-resource isotopic data. Consequently, many individuals cannot be linked to corresponding accessible food resources, a limitation that becomes more pronounced in large-scale quantitative dietary reconstructions incorporating a wide range of food-resource categories. Therefore, this study aims to broaden the definition of accessible food resources. To achieve this aim, we compiled food-resource and soil isotopic data (δ^13^C and δ^15^N) from Britain, including 4,012 ancient faunal and plant remains, 394 modern plant samples, and 260 modern soil samples. Region–period combined groups were established for the five major food-resource categories and served as the basic analytical units for detailed isotopic comparisons. Based on these comparisons, we propose broader criteria for defining accessible food resources. No significant intra-group variation was observed in the isotopic values of terrestrial herbivores and omnivores, suggesting that animals within each region–period combined group can serve as accessible food resources for humans from the same group. C_3_ plants showed substantial spatial variation but limited temporal variation. Accordingly, accessible food resources for C_3_ plants should be defined by region, namely England, Wales, and Scotland, regardless of chronological period, with humans from each region assigned plant data from their respective region. Marine and freshwater fish showed no clear temporal or spatial variation, and therefore unified datasets can be applied across all human individuals. Our findings enable each ancient human individual to be assigned appropriate accessible food resources, and therefore appropriate food-resource isotope baselines. We further demonstrate that such assignments can effectively reduce sampling bias arising from the use of traditionally defined accessible food resources, which are often limited by small sample sizes.

## Introduction

Dietary reconstruction using stable isotope analysis, whether through qualitative scatterplot comparisons[1] or quantitative Bayesian modeling[2–4], requires isotopic values of food resources accessible to the target population. Ideally, accessible food resources should originate from the same site as the target individuals and possess consistent chronological information (e.g., the study of diet in Paleolithic Gough’s Cave and Sun Hole)[5]. In cases where such direct food-resource data are unavailable, food resources from geographically and temporally proximate sites are sometimes used as substitutes (e.g., the study of diet in Anglo-Saxon York)[6].

These definitions of accessible food resources are generally feasible when dietary reconstruction involves only a small number of individuals from a few sites or focuses on a limited set of food-resource categories, in which case each individual can often be matched with traditionally defined accessible food resources. However, when dietary reconstruction involves a large number of individuals or incorporates five major food-resource categories (terrestrial herbivores, terrestrial omnivores, C_3_ plants, marine resources, and freshwater resources), not every individual can be matched with traditionally defined accessible food resources. As shown in Fig. 1, the number of sites with human samples greatly exceeds that of sites with samples from each food-resource category. Beyond the number of human samples and the breadth of food-resource categories considered in dietary reconstruction, the distinction between quantitative and qualitative approaches also influences the applicability of traditionally defined accessible food resources. In qualitative dietary reconstruction, approximate dietary interpretations can still be made even when accessible food resources for a particular food-resource category are unavailable. For example, even if isotope data for accessible freshwater fish resources are lacking for a given region, anomalously high human δ^15^N values relative to terrestrial animal baselines, in the absence of corresponding δ^13^C enrichment and after excluding potential manuring effects, may still suggest possible freshwater fish consumption, particularly when sites are located close to rivers[7]. By contrast, in quantitative dietary reconstruction, accessible food resources must be assigned to each individual[3]. Thus, when attempting to construct quantitative dietary models that incorporate five major food-resource categories for large numbers of human individuals (e.g., all ancient British individuals), the problem of lacking accessible food resources becomes particularly pronounced.

**Fig. 1.**
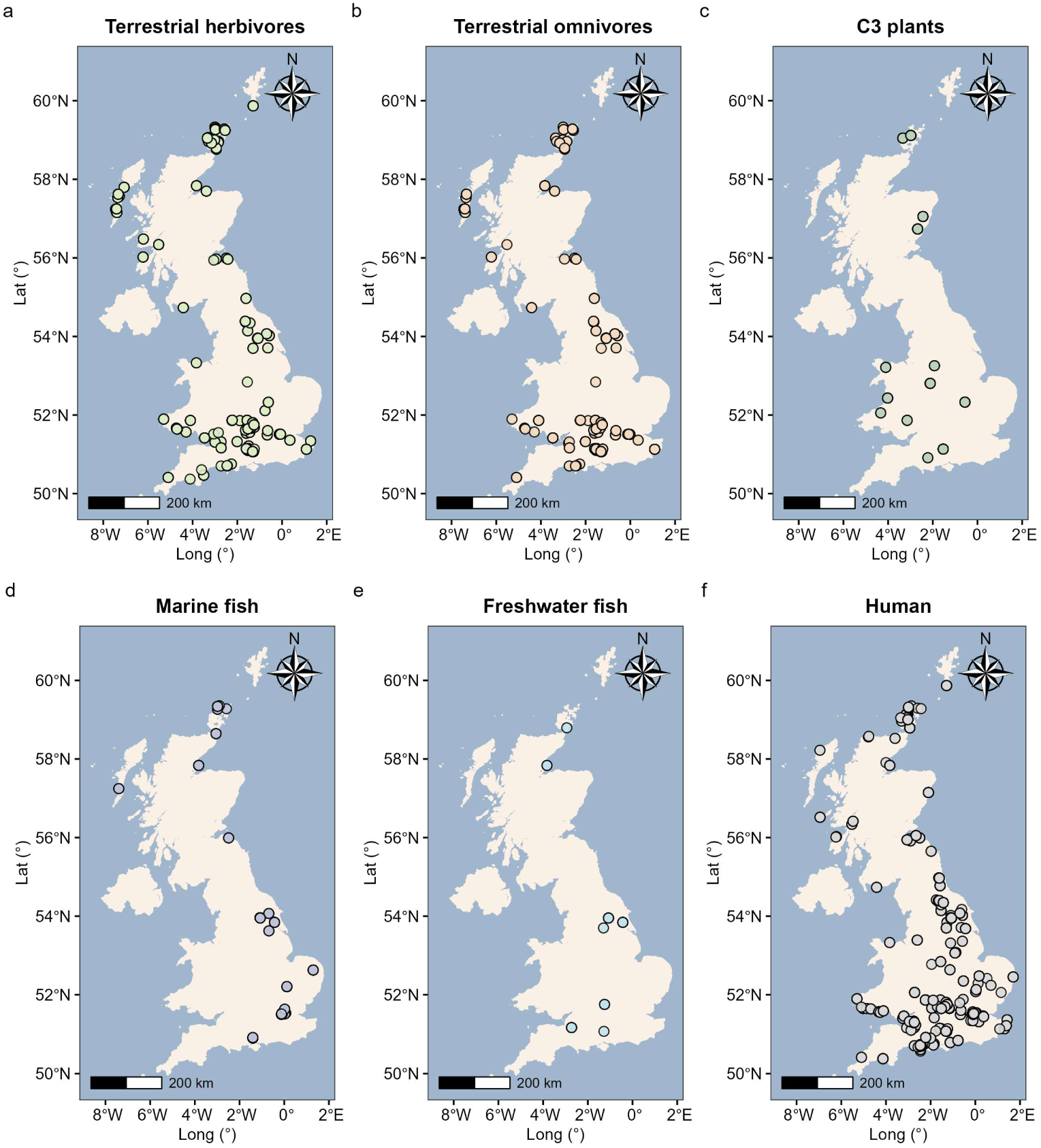
Spatial distributions of food-resource and human isotopic samples in Britain. **a,** Terrestrial herbivores. **b,** Terrestrial omnivores. **c,** C_3_ plants. **d,** Marine fish. **e,** Freshwater fish. **f,** Human.

In such cases, it becomes particularly important to determine how to reasonably extend the spatial and temporal limits of the accessible food resources. Therefore, this study aims to broaden the definition of accessible food resources by investigating whether food resources from wider temporal and spatial contexts, such as those from Bronze Age England, can be used as accessible food resources for the dietary reconstruction of human populations from the same broad period and region. To achieve this, we compiled stable isotope data (δ^13^C, δ^15^N) from 4,012 ancient faunal and plant remains[5–81], 394 modern plant remains[71,82–86], and 260 modern soil samples[82,87,88], thereby covering nearly all currently published food-resource isotope data from Britain, spanning the Paleolithic to the modern period. We then classified the collected samples into different region-period combined groups according to their locations and dating information (see Materials and methods for details). Using the combined groups as the basic analytical units, we conducted comprehensive isotopic comparisons by integrating qualitative assessments, statistical tests[89–95], and geographical analyses (see Results). Based on the comparative results, we propose a broader criterion for defining accessible food resources across five food categories, thereby enabling the construction of robust isotopic baselines in a broader manner and ensuring that each individual can be assigned appropriate accessible food resources in large-scale dietary reconstruction. In addition, we demonstrate that baselines constructed in this manner can produce more robust estimates by reducing sampling bias caused by using food samples from only one or a few sites as accessible food resources, where sample sizes are often small and therefore more prone to bias.

## Materials and methods

### Data collection

We collected stable isotope data from 4,012 ancient British faunal and plant remains[5–81], 394 modern British plant remains (grain from Scotland and plant biomass from Scotland and southern England) [71,82–86], and 260 modern British soil samples (from Scotland and southern England) [82,87,88]. Relevant studies from which these data were collected were identified through searches of Web of Science and Google Scholar using the keywords “stable isotopes”, “dietary reconstruction”, “food resources”, “δ^13^C”, “δ^15^N”, and “Britain”. The literature search included publications available up to November 2023. Detailed information for each sample is provided in Supplementary Data 1, including the sample ID, location (region, site name, latitude, and longitude), dating (period, dating results, and dating method), material analyzed (e.g., bone or dentine), isotopic values (δ^13^C, δ^15^N), data quality indicators (C/N, %C, %N, and collagen preservation), and the corresponding references. The categorization of food resource samples (e.g., cattle, terrestrial herbivores) is also documented in Supplementary Data 1.

### Regional and chronological grouping of ancient samples

We divided Britain into five regions: southern England (including samples from Doggerland), central England, northern England, Scotland, and Wales (see Supplementary Data 1). England was divided into northern, central, and southern regions based on the Government Office Regions of England. Northern England comprises the North East, North West, and Yorkshire and the Humber; central England comprises the East and West Midlands; southern England comprises the East of England, London, South East, and South West. Scotland and Wales were treated as separate regions following their existing administrative boundaries. The chronology was divided into nine periods: Paleolithic period (before 9000 BC), Mesolithic period (9000–4300 BC), Neolithic period (4300–2200 BC), Bronze Age (2200–750 BC), Iron Age (750 BC–AD 43), Roman period/Roman Iron Age (AD 43–410), Early Medieval period (AD 410–1066), Later Medieval period (AD 1066–1540), and Post-Medieval period (AD 1540–1900). Based on each sample’s latitude, longitude, and dating, we assigned them to the appropriate region–period combined groups (e.g., Iron Age southern England). This grouping scheme ensured relatively large food-resource sample sizes within each group while preserving sufficient temporal and geographical resolution. Some food resource samples span two periods (e.g., AD 0–99, which falls within both the Iron Age and Roman periods) and were therefore considered members of both groups (highlighted in green in Supplementary Data 1).

### Exclusion of rare food categories in ancient samples

Even after dividing samples into region–period combined groups, many species remained unavailable for most groups, particularly terrestrial carnivores, marine carnivores, marine omnivores, freshwater carnivores, freshwater omnivores, and freshwater herbivores. These six categories together comprise only 73 specimens, a minimal number compared with the 4,012 ancient food resource samples in total. Moreover, such species were likely not typical components of ancient human diets (e.g., foxes were rarely consumed). Therefore, we excluded these categories from further analyses. After this exclusion, five major food categories remained: terrestrial plants, terrestrial omnivores, terrestrial herbivores, marine fish, and freshwater fish. These categories represent the most common human food resources and were therefore the focus of subsequent comparison analyses.

### Comparison strategy by food group: ancient samples

For terrestrial herbivores (2,351 samples) and omnivores (857 samples), we conducted detailed comparisons of isotopic values among sites within each group (intra-group comparison). In addition, isotopic values were compared across groups (inter-group comparison) within the same period to explore broader regional patterns of variation (e.g., Iron Age: southern England vs. northern England vs. Scotland). If the isotopic variation within each group is not significant, this indicates that isotopic values are broadly similar among sites within the same group. Accordingly, the food resources within a group can be defined as the accessible food resources for humans belonging to the same group (i.e., used for reconstructing their diets).

Archaeological sites specifically define the locations of the samples. However, some samples from the same site may belong to different periods even if they fall within the same combined group (e.g., Mesolithic: 8500 BC vs. 6500 BC). In such cases, the site was treated as multiple independent sites, designated by the site name followed by a number (e.g., Foxhole Cave 1; Foxhole Cave 2). Given the large number of sites, abbreviations are used in the following sections (e.g., Sudden Farm: SF1; Foxhole Cave 1: FC1; Foxhole Cave 2: FC2). The full site names and corresponding abbreviations are provided in the associated figures.

Terrestrial plants (400 samples) and marine fish (280 samples) were insufficiently represented for such detailed intra-group analyses. Therefore, for C_3_ plants and marine fish, isotopic variation was assessed through inter-group comparisons, in which values were compared across regions while controlling for time (e.g., Iron Age: Scotland vs. southern England vs. Wales) and across time periods while controlling for region (e.g., southern England: Bronze Age vs. Iron Age vs. Later Medieval). This is to evaluate whether isotopic values varied across space and time. When isotopic values vary across regions but remain stable over time, accessible food resources can be defined regionally (e.g., England, Scotland, and Wales), ignoring temporal variation, and vice versa.

Given the very limited number of freshwater fish (51 samples), data from all region–period combined groups were pooled for comparison. If isotopic values show little variation across all combined groups, or if the isotopic values of groups with smaller sample sizes fall within the range defined by those with larger sample sizes, this suggests that all human populations could share a common freshwater fish resource pool.

### Comparison strategy by food group: modern samples

The modern isotopic dataset includes only a limited number of C_3_ plants and soil samples. These data were used as auxiliary evidence to support the ancient dataset, providing additional insights into whether spatial variation, under consistent temporal conditions, significantly affects plant isotopic values. The analytical strategy was straightforward. For plants, we directly compared grain from Scotland with plant biomass from Scotland and southern England. For soil, we directly compared soil samples from Scotland and southern England.

### Statistical methods for isotopic data comparison

Isotopic data were compared using scatterplot-based qualitative assessments and, where sample sizes permitted, non-parametric statistical tests. Geographical analyses were conducted alongside isotopic comparisons when necessary. Because our data distributions were often non-normal, variances unequal, and sample sizes across sites small or unbalanced, we applied non-parametric tests to δ^13^C and δ^15^N. For two-group comparisons, we used the Wilcoxon rank–sum test (Mann–Whitney U)[94,95]. For comparisons involving more than two groups, we used the Kruskal–Wallis test[92]. When the omnibus test was significant, pairwise post hoc comparisons were conducted with the Wilcoxon rank–sum test, and p-values were adjusted for multiple testing using the Holm method[89]. Statistical tests are not conducted for all comparisons. When the sample sizes of the sites or combined groups being compared are highly unbalanced (e.g., Fig. 4), or when all sites/groups being compared have very small sample sizes (e.g., Fig. 3), statistical testing is not conducted. Scatterplot-based qualitative assessment is the main method used in these two cases. For the first case, qualitative assessment focuses on whether the isotopic values from the sparsely sampled sites/groups fall within the isotopic range defined by the well-sampled sites/groups. For the second case, attention is given to the isotopic distributions of the sites/groups being compared. However, because the sample sizes of all sites/groups being compared are very small, these comparisons are mainly used to illustrate the observed isotopic distributions and are not used to draw firm conclusions. For some region–period groups, sites with very small sample sizes (sparsely sampled sites) were excluded from the statistical analyses; only sites with sufficient sample sizes (well-sampled sites) were retained (e.g., Fig. 2).

**Fig. 2.**
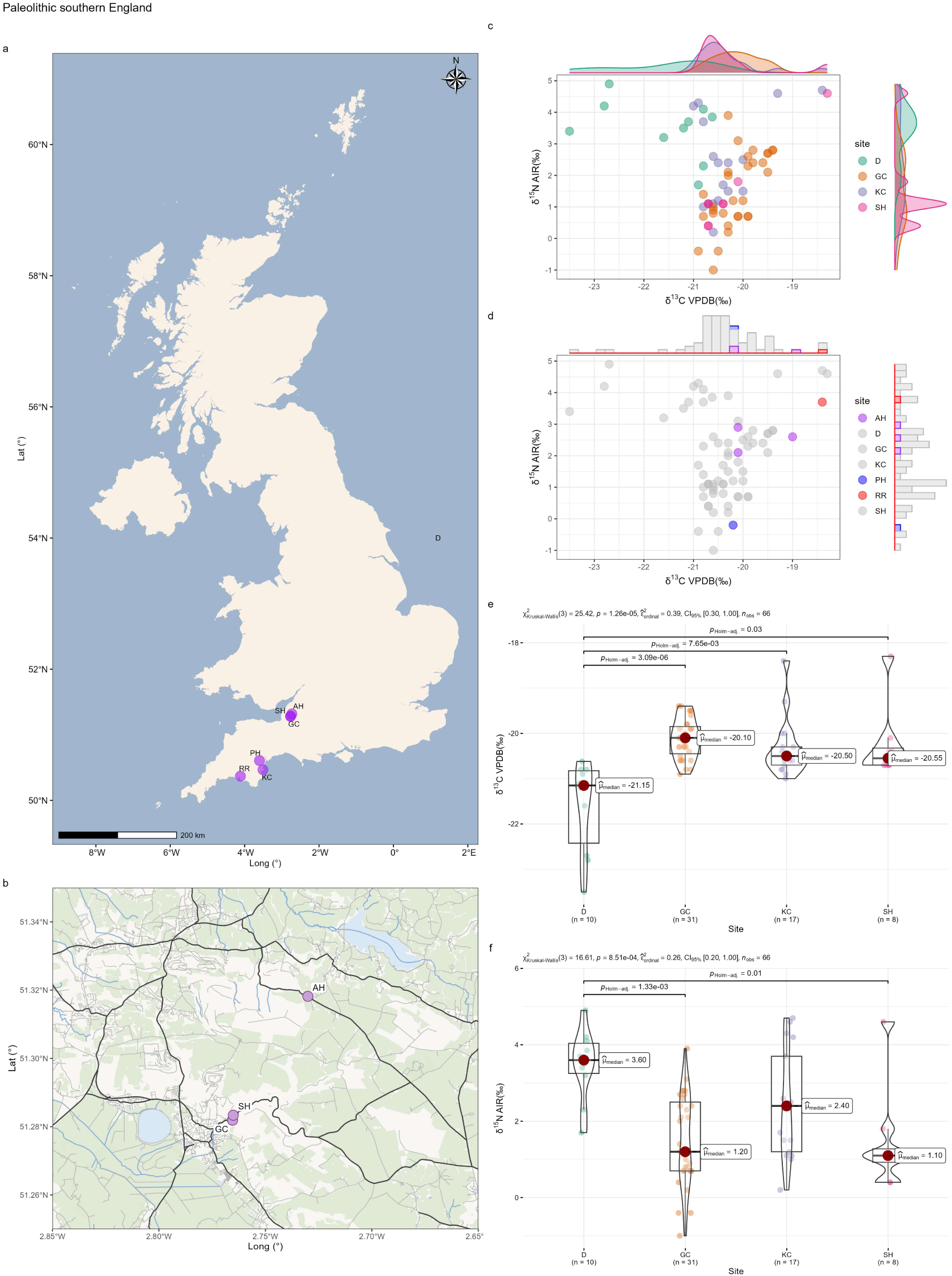
Intra-group comparison of terrestrial herbivores from Paleolithic southern England. **a,** Distribution of sites. **b,** Distribution of three proximate sites. **c,** Scatter plot (well-sampled sites). **d,** Scatter plot (sparsely sampled sites). **e,** Statistical tests for δ^13^C (well-sampled sites). **f,** Statistical tests for δ^15^N (well-sampled sites). Site names: D: Doggerland; GC: Gough’s Cave; KC: Kent’s Cavern; SH: Sun Hole; AH: Aveline’s Hole; PH: Pixies’ Hole; RR: Reindeer Rift.

**Fig. 3.**
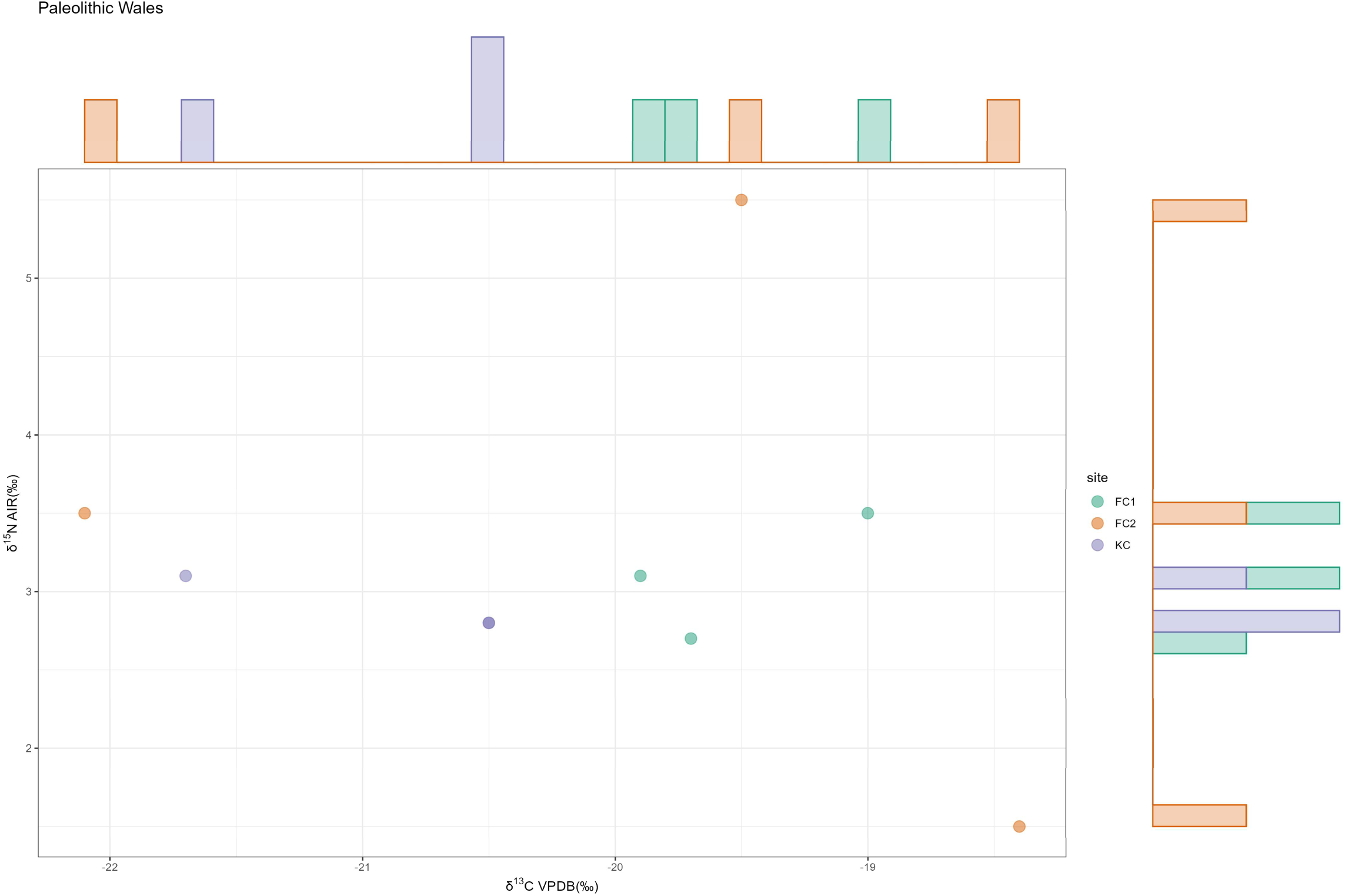
Intra-group comparison of terrestrial herbivores from Paleolithic Wales (Scatter plot). Site names: FC1: Foxhole Cave 1; FC2: Foxhole Cave 2; KC: Kendrick’s Cave.

**Fig. 4.**
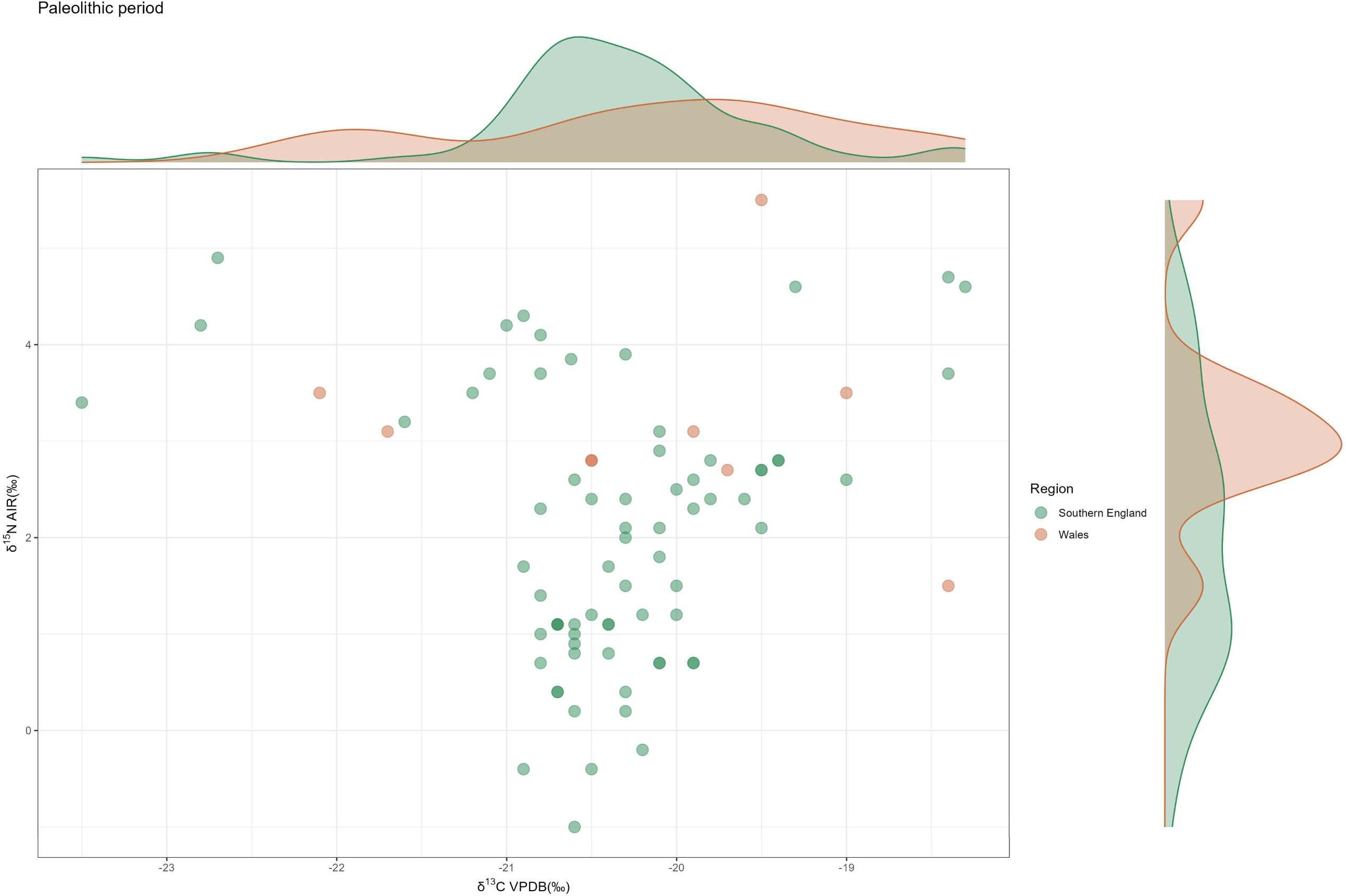
Inter-group comparison of terrestrial herbivores from the Paleolithic period (Scatter plot).

Data distributions were visualized with boxplots, violin plots, and associated kernel density estimates (KDE) using the default bandwidth selection method (Silverman’s rule-of-thumb)[93]. For some combined groups with a large number of sites, heatmaps were additionally generated with hierarchical clustering[90]. The dendrogram was generated by hierarchical clustering of a pairwise dissimilarity matrix constructed from post hoc p-values (Holm-adjusted), with Ward’s D linkage[91].

## Results

### Analysis of isotopic variation for terrestrial herbivores

#### Paleolithic and Mesolithic periods

For the Paleolithic period, analyses can be conducted on the Paleolithic southern England combined group. The overall pattern of this group is presented in Fig. 2. The scatter plots reveal no distinct clustering, with data from all archaeological sites largely overlapping, except for a subset of samples from Doggerland (Fig. 2c). This pattern is further supported by statistical tests: although both analyses indicate significant differences, the post hoc tests suggest that these differences are primarily driven by Doggerland (Fig. 2e, f). By contrast, the other three groups exhibit similar δ^13^C and δ^15^N patterns. Isotopic values from sites with insufficient sample sizes for statistical tests are displayed in Fig. 2d. These samples fall within the main cluster. The geographic locations of the Paleolithic southern England sites are shown in Fig. 2a and b. With the exception of Doggerland, all sites are situated in Southwest England, suggesting that the distinctive isotopic signal of Doggerland may reflect its different geographical setting.

For the Paleolithic Wales combined group, the Kruskal–Wallis test could not be performed due to the small sample sizes of individual sites. The scatter plot (Fig. 3) shows that samples from FC1 and KC cluster together, whereas those from FC2 are more widely dispersed. This indicates that the isotopic values of FC1 and KC fall within the range of variations observed at FC2. Since isotopic values from the same site (e.g., FC2) are expected to be highly similar, this comparison indicates that there are no significant isotopic differences among the Paleolithic Welsh sites. However, given the limited sample sizes, this conclusion should be treated with caution. The inter-group comparison of isotopic values between Paleolithic southern England and Wales (Fig. 4) shows that all Welsh samples fall within the main cluster of southern England.

For the Mesolithic period, site comparisons can be made between two groups: Mesolithic southern England and Mesolithic Scotland. The Mesolithic southern England group includes three archaeological sites—AH, H, and Doggerland (subdivided into periods D1, D2, and D3). However, the sample sizes for AH and H are very small (n = 3 for each), making it difficult to assess potential differences. The samples from different periods of Doggerland overlap substantially (Fig. 5a). Statistical tests were performed for the two groups with larger sample sizes (D1 and D2), but no significant differences were detected (Fig. 5c, d). Isotopic values from the small-sample sites (AH, H, and D3) were also added to the scatter plot, and these values largely fall within the cluster defined by D1 and D2 (Fig. 5b).

**Fig. 5.**
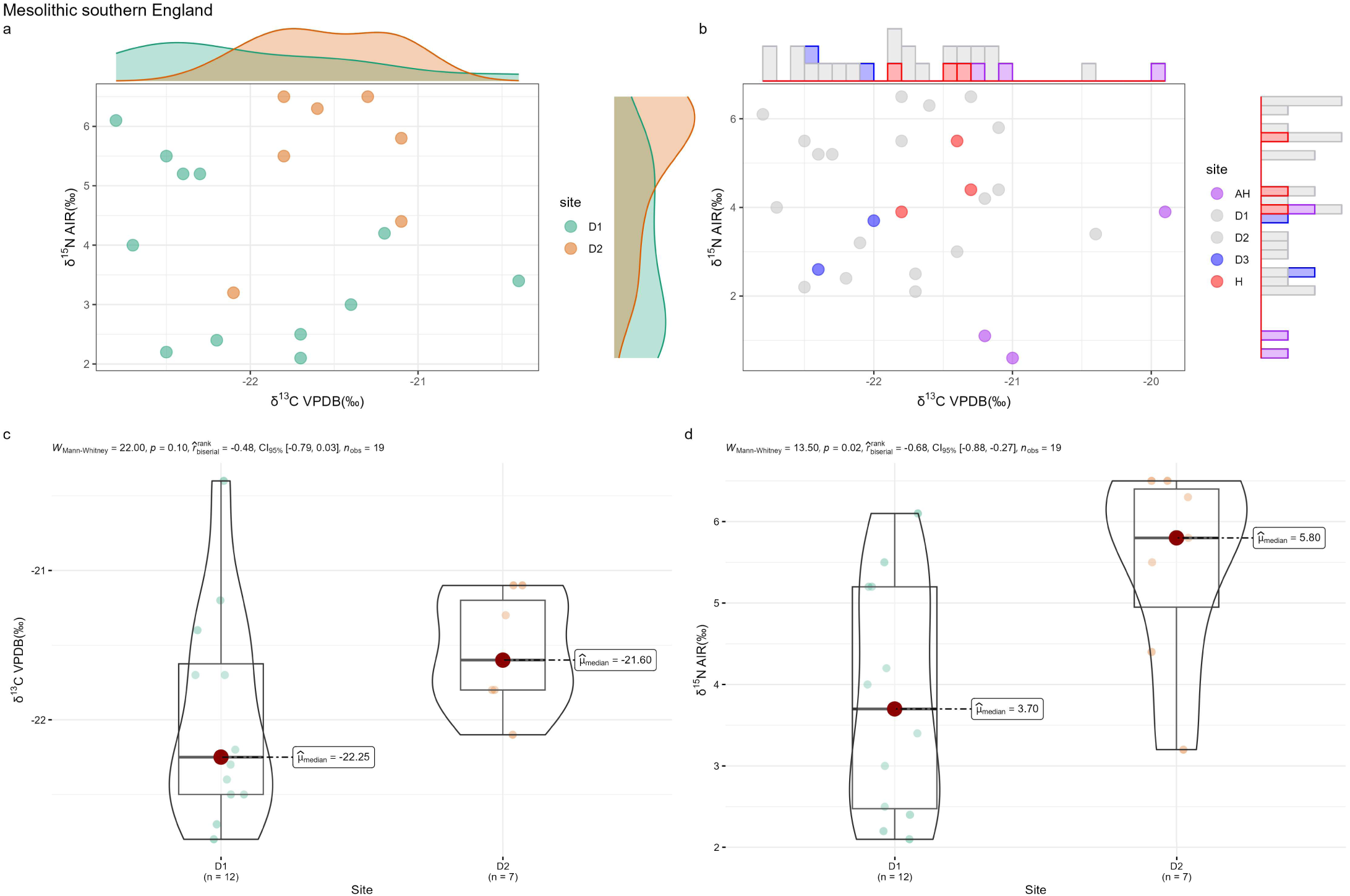
Intra-group comparison of terrestrial herbivores from Mesolithic southern England. **a,** Scatter plot (well-sampled sites). **b,** Scatter plot (sparsely sampled sites). **c,** Statistical tests for δ^13^C (well-sampled sites). **d,** Statistical tests for δ^15^N (well-sampled sites). Site names: D1: Doggerland 1; D2: Doggerland 2; AH: Aveline’s Hole; D3: Doggerland 3; H: Hazleton.

For the Mesolithic Scotland group, the scatter plot shows that all sites cluster together, with the exception of two outliers from Risga (Fig. 6a). The statistical tests indicated significant post hoc differences in δ^13^C between CM and R, as well as between CM and UC (Fig. 6c, d). However, although the results reached statistical significance, the p-values were relatively high (all equal to 0.04). In addition, the differences in both mean and median values between CM and the other two sites are less than 1‰, suggesting that these statistically significant results may not be of practical relevance. Samples from sparsely sampled sites (SC and WV) also fall within the main cluster (Fig. 6b). A comparison of isotopic values from Mesolithic southern England, Scotland, and Wales demonstrates that all three regions exhibit similar isotopic signatures (Fig. 7). The samples from southern England and Scotland overlap, while the single sample from Wales falls within their cluster.

**Fig. 6.**
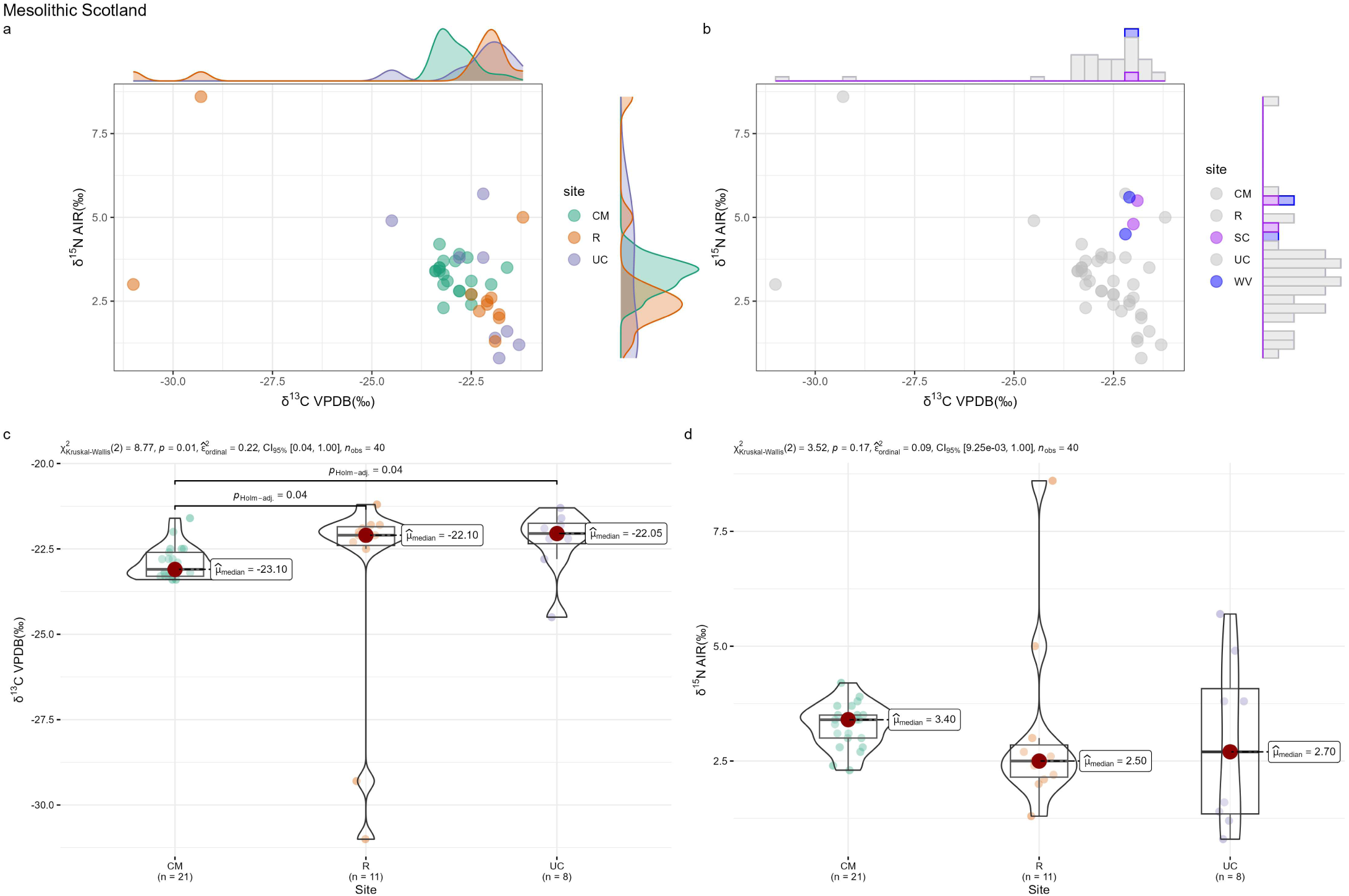
Intra-group comparison of terrestrial herbivores from Mesolithic Scotland. **a,** Scatter plot (well-sampled sites). **b,** Scatter plot (sparsely sampled sites). **c,** Statistical tests for δ^13^C (well-sampled sites). **d,** Statistical tests for δ^15^N (well-sampled sites). Site names: CM: Carding Mill Bay; R: Risga; UC: Ulva Cave; SC: Sumburgh cist; WV: West Voe.

**Fig. 7.**
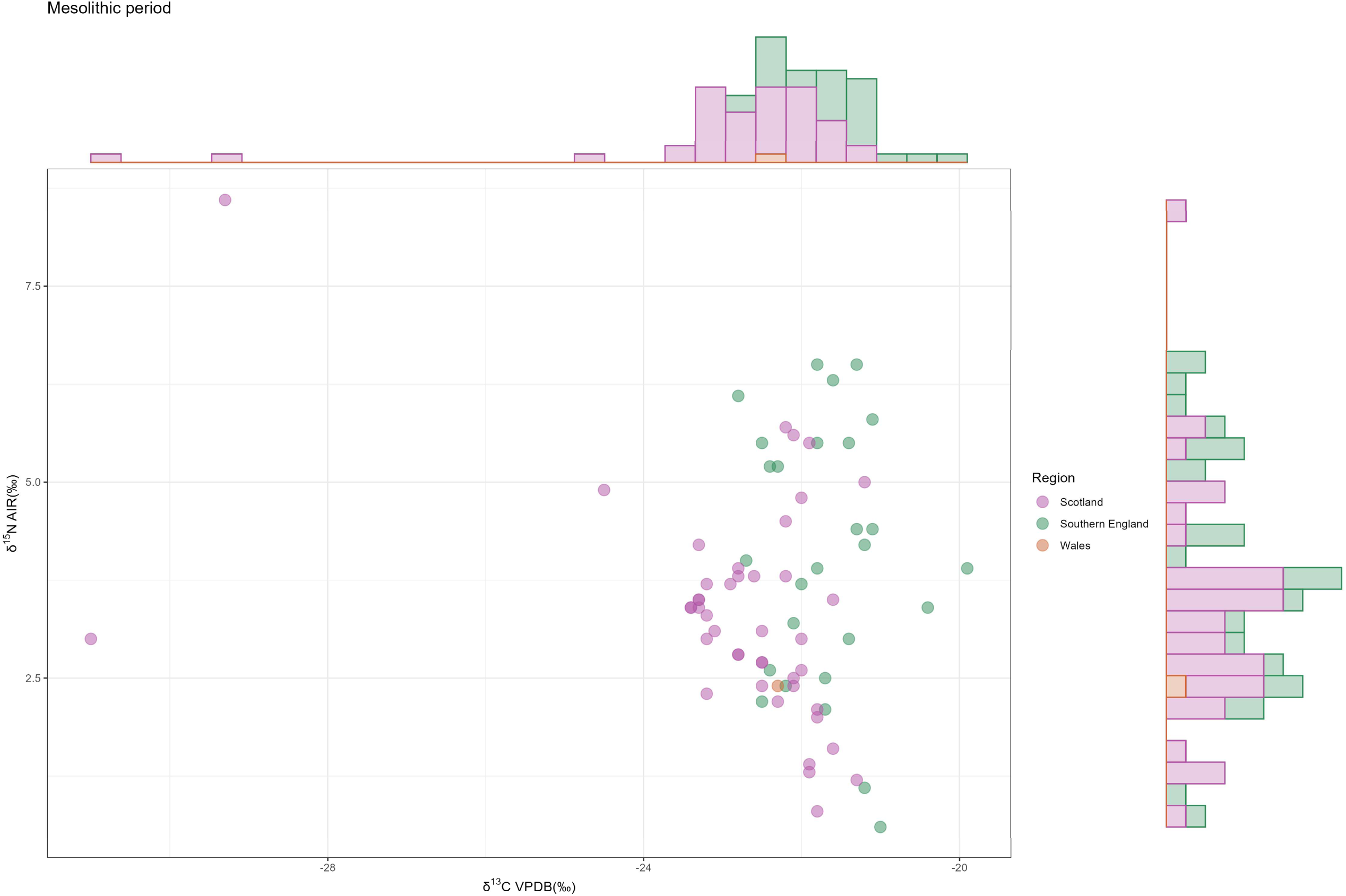
Inter-group comparison of terrestrial herbivores from the Mesolithic period (Scatter plot).

For both the Paleolithic and Mesolithic terrestrial herbivore datasets, no significant isotopic differences are detected among sites within the same combined group, apart from a subset of samples from Doggerland. Broader inter-group comparisons within each period likewise show no clear isotopic distinctions (Fig. 7). Isotopic values from groups with comparable sample sizes generally overlap (e.g., Mesolithic southern England vs. Scotland), whereas those from smaller datasets consistently fall within the clusters defined by larger ones (e.g., Mesolithic Wales).

### Neolithic period

Similar to the Mesolithic, the Neolithic dataset can also be divided into two region-period combined groups for comparison: Neolithic southern England and Neolithic Scotland. In Neolithic southern England, isotopic values from HN differ significantly from those of the other two sites, particularly with respect to δ^15^N (Fig. 8b-d). The geographic locations of all sites in southern England are shown on the maps (Fig. 8a). The three sites are located in close proximity. Notably, HN, which exhibits significantly different isotopic values, lies even closer to AUW than AUW does to EC.

**Fig. 8.**
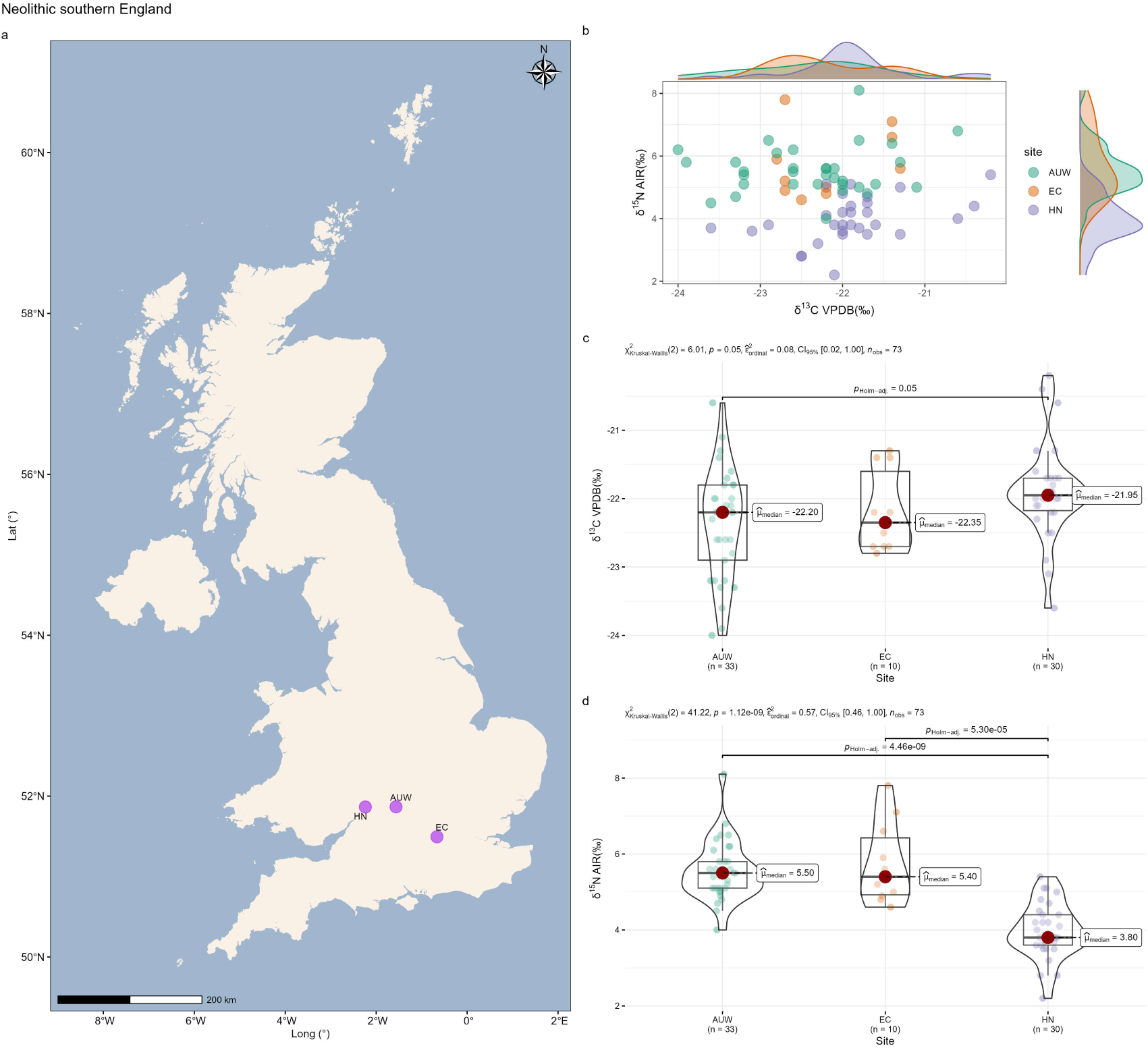
Intra-group comparison of terrestrial herbivores from Neolithic southern England. **a,** Distribution of sites. **b,** Scatter plot. **c,** Statistical tests for δ^13^C. **d,** Statistical tests for δ^15^N. Site names: AUW: Ascott-Under-Wychwood; EC: Eton College Rowing Course; HN: Hazleton North.

For Neolithic Scotland, we plotted mean isotopic values with standard deviations for each site (Fig. 9c). The results show that samples from the same site but different phases (LN1 and LN2) display remarkably similar isotopic signatures, and thus were merged into a single site (i.e., LN). Subsequent scatter plots and statistical tests reveal that site Q (δ^13^C) and site TN (δ^15^N) differ significantly from the others (Fig. 9d, f, g). Apart from these two sites, the remaining seven exhibit broadly similar δ^13^C and δ^15^N values. Two sites with small sample sizes (CB and PC) show anomalous isotopic values, which fall outside the main cluster defined by the sites with larger sample sizes (Fig. 9e). Geographic evidence indicates that sites with anomalous isotopic values are located in close proximity to most of the other sites (Fig. 9a, b). A closer examination shows that although site P is immediately adjacent to TN (virtually the same location), its δ^15^N values differ significantly (Fig. 9g). Similarly, site Q is situated near NB, yet its δ^13^C values differ markedly from those of NB (Fig. 9f).

**Fig. 9.**
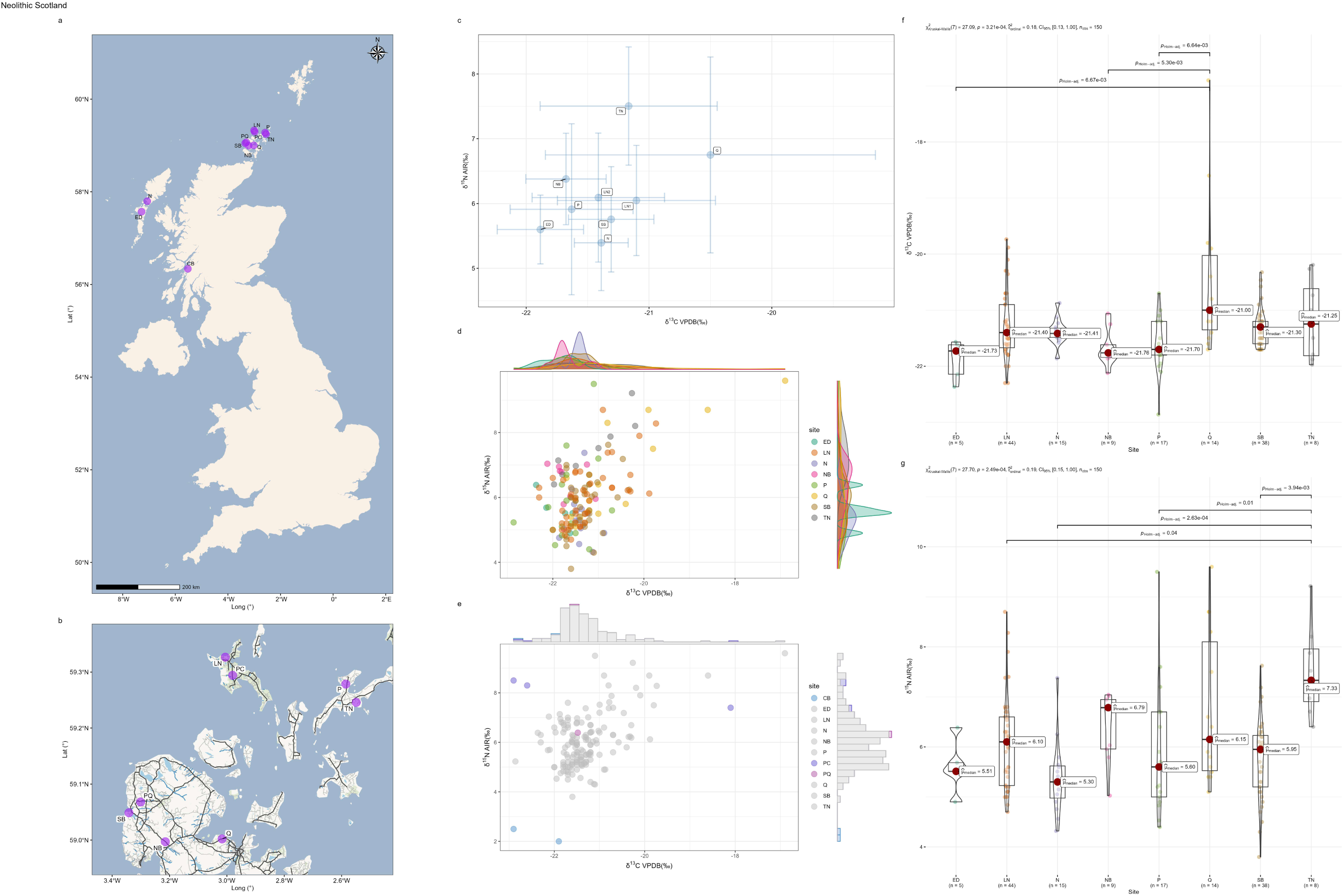
Intra-group comparison of terrestrial herbivores from Neolithic Scotland. **a,** Distribution of sites. **b,** Distribution of eight proximate sites. **c,** Average isotopic values with standard deviations (well-sampled sites). **d,** Scatter plot (well-sampled sites). **e,** Scatter plot (sparsely sampled sites). **f,** Statistical tests for δ^13^C (well-sampled sites). **g,** Statistical tests for δ^15^N (well-sampled sites). Site names: ED: Eilean Domhnuill; LN: Links of Noltland (LN1: Links of Noltland 1; LN2: Links of Noltland 2); N: Northton; NB: Ness of Brodgar; P: Pool; Q: Quanterness; SB: Skara Brae; TN: Tofts Ness. CB: Carding Mill Bay; PC: Point of Cott; PQ: Pierowall Quarry.

This phenomenon raises doubts about the validity of the statistical tests. Identifying isotopic values from two sites in extremely close proximity (e.g., P and TN) as significantly different appears unreasonable. We therefore interpret the significant results from the statistical tests as reflecting normal intra-site variability rather than meaningful inter-site differences. This may be due to the small sample size at individual sites and the uneven distribution of samples across sites. As a result, the data from a single site may be insufficient to capture the overall isotopic characteristics of that site, and sampling bias may have led to statistically significant results between sites that are located in very close proximity. Accordingly, the statistical results lack practical significance, and no substantial isotopic differences can be identified among sites within these two combined groups.

The isotopic values from Neolithic Wales are shown in Fig. 10. However, the sample sizes for each site are too small to support firm conclusions. A comparison of samples from southern England, Scotland, and Wales (Fig. 11) shows that the Welsh specimens (n = 5) fall within the clusters defined by southern England and Scotland. Most samples from southern England and Scotland overlap, although a small subset of Scottish samples (approximately 20, primarily from Q, LN, and TN) differ significantly from those of southern England.

**Fig. 10.**
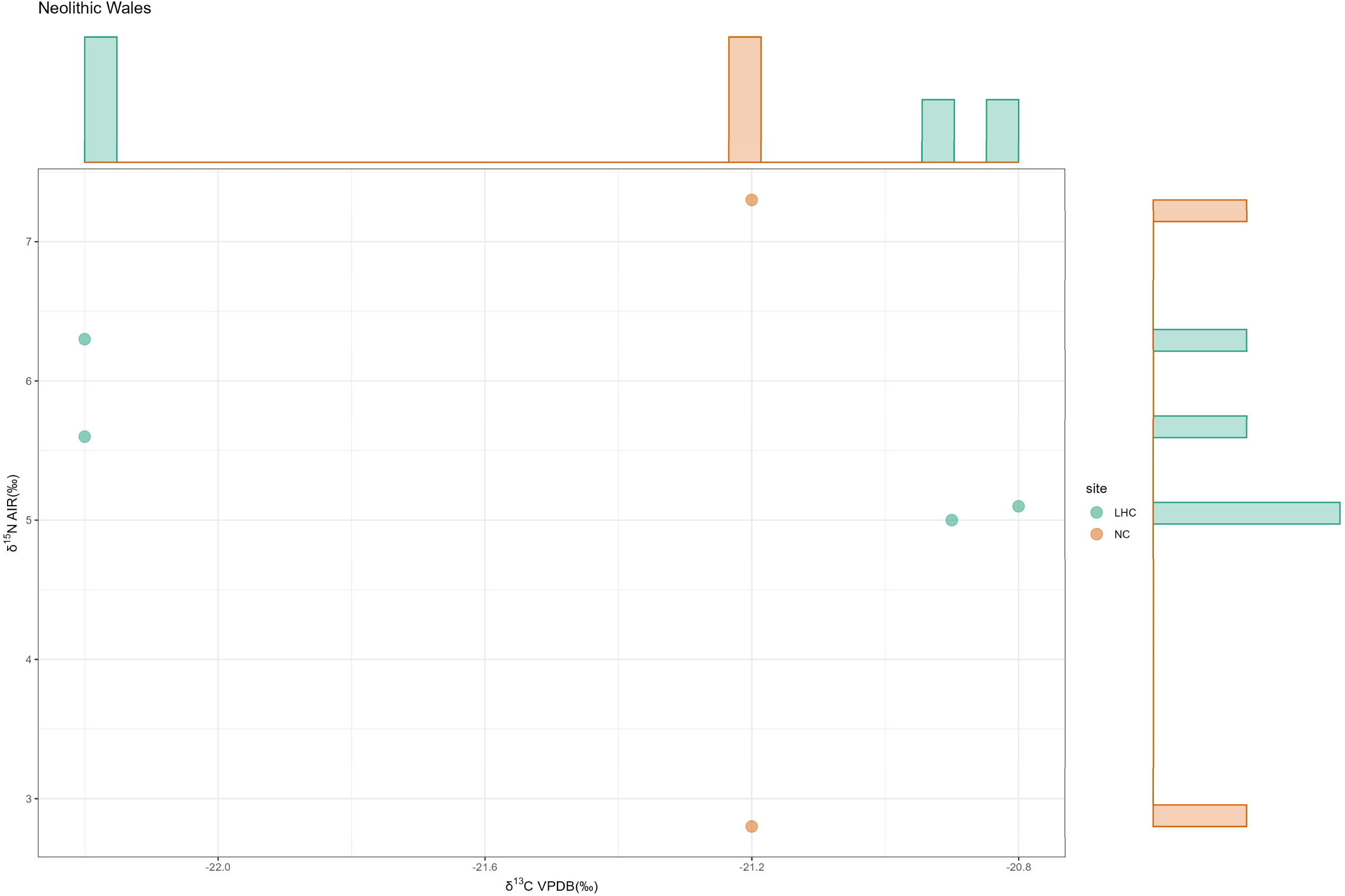
Intra-group comparison of terrestrial herbivores from Neolithic Wales (Scatter plot). Site names: LHC: Little Hoyle Cave; NC: Nanna’s Cave.

**Fig. 11.**
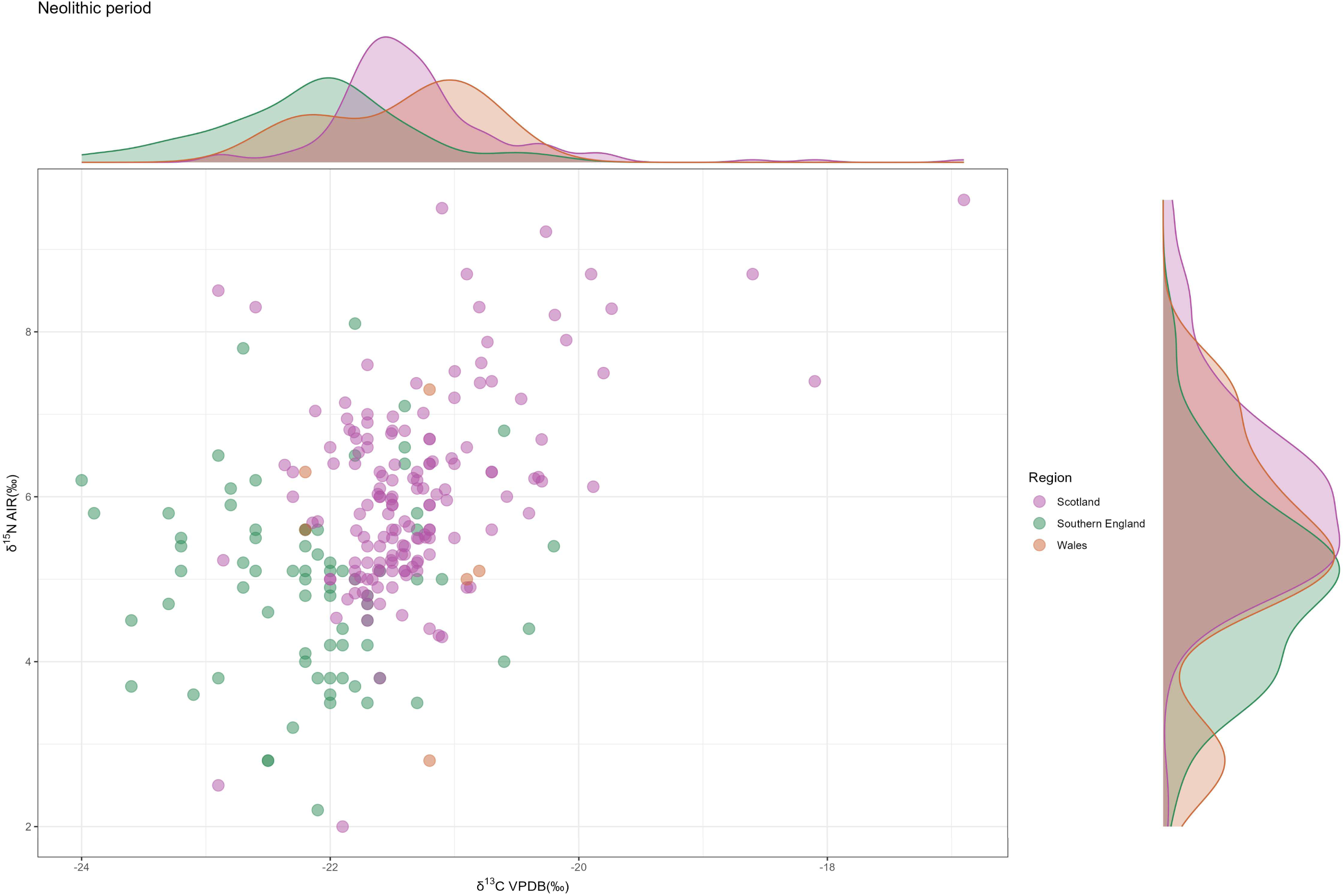
Inter-group comparison of terrestrial herbivores from the Neolithic period (Scatter plot).

Comparisons of the Neolithic datasets indicate that within each combined group, no substantial isotopic differences exist among sites. Although some statistical tests yielded significant results, consideration of the geographic proximity of these sites suggests that these findings likely lack practical significance and instead reflect normal intra-site variability. Inter-group comparisons show relatively pronounced differences between samples from southern England and Scotland, although the isotopic values of most samples still overlap.

### Bronze Age

The Bronze Age dataset can also be divided into two groups for comparison: Bronze Age southern England and Bronze Age Scotland.

For Bronze Age southern England, we first plotted the mean isotopic values with standard deviations for each site (Fig. 12b). Similar to the pattern observed in Neolithic Scotland, samples from the same site but different phases display highly consistent isotopic values (e.g., BD1 vs. BD2; P1 vs. P2), and were therefore treated as single sites. The scatter plot reveals clear clustering of the samples (Fig. 12c), forming three main clusters: Cluster 1 includes sites G and I; Cluster 2 contains site P; and Cluster 3 comprises site BD. The exceptional site EC (n = 5) shows a mixed pattern, with some samples falling into Cluster 2 and others into Cluster 3. This clustering is further supported by the statistical tests (Fig. 12d, e). For both δ^13^C and δ^15^N, no significant differences are evident among sites within the same cluster. In contrast, comparisons among sites across clusters reveal statistically significant differences, such as the δ^15^N contrasts between BD (Cluster 3) and G (Cluster 1), and between BD (Cluster 3) and P (Cluster 2) (Fig. 12e). For the exceptional site EC, its δ^13^C values resemble those of site P (Fig. 12d). However, its δ^15^N values span the widest range among all sites, even though the site is represented by only five samples. (Fig. 12e). The clusters correspond broadly to geography (Fig. 12a): sites in Cluster 1 (I and G) are located close to one another and relatively distant from those in Clusters 2 and 3 (BD, P, EC). Accordingly, Bronze Age southern England clearly demonstrates isotopic variation among sites.

**Fig. 12.**
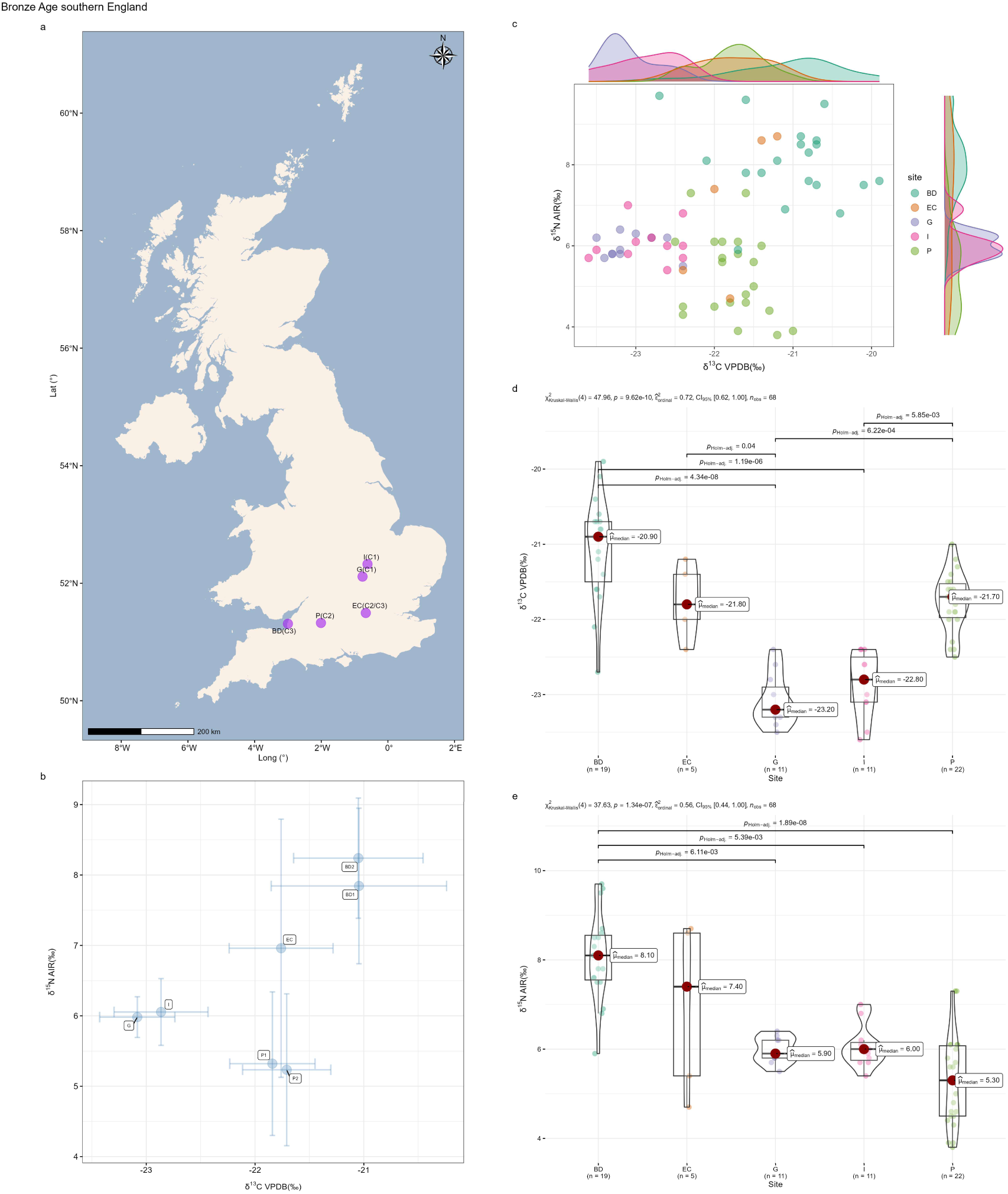
Intra-group comparison of terrestrial herbivores from Bronze Age southern England. **a,** Distribution of sites. C1–C3 denote Cluster 1 to Cluster 3. **b,** Average isotopic values with standard deviations. **c,** Scatter plot. **d,** Statistical tests for δ^13^C. **e,** Statistical tests for δ^15^N. Site names: BD: Brean Down (BD1: Brean Down 1; BD2: Brean Down 2); EC: Eton College Rowing Course; G: Gayhurst; I: Irthlingborough; P: Potterne (P1: Potterne 1; P2: Potterne 2).

For the Bronze Age Scotland combined group, the scatter plot shows that site CH spans a wide isotopic range (Fig. 13c). Most isotopic values from sites S and TN fall within this range, although they form distinct subclusters within the range of CH. Both sites have much smaller sample sizes than CH, which means the small sample sizes of S and TN render them unrepresentative. Therefore, the distinct isotopic distributions they exhibit may be attributable to the effect of sample size. Five samples from site SC form a separate cluster outside CH (higher values in δ^15^N). The statistical test of δ^13^C yields significant results (Fig. 13d), with post hoc tests indicating significant differences between SC and CH (p < 0.001), as well as between SC and TN (p < 0.05). However, as discussed above, these results may be largely driven by differences in sample size, making the statistical tests potentially misleading. For δ^15^N, post hoc tests reveal highly significant differences between S and TN (p < 0.001), whereas CH vs. TN and S vs. SC yield lower but still significant p-values (0.01 < p < 0.05) (Fig. 13e). As with δ^13^C, this chaotic pattern is likely an artifact of sample size rather than a reflection of genuine isotopic differences. The comparisons of S vs. TN, CH vs. TN, and S vs. SC may lack interpretive value because of the small sample sizes of S and TN. Isotopic values from the site Q (small sample size) are shown in Fig. 13b. Apart from one outlier, all samples fall within the main cluster. The isotopic values of sites in Scotland show no geographic pattern (Fig. 13a). As discussed above, the apparent significant results within Scotland are likely attributable to sample size effects. Thus, no genuine isotopic differences exist, let alone any geographic structure.

**Fig. 13.**
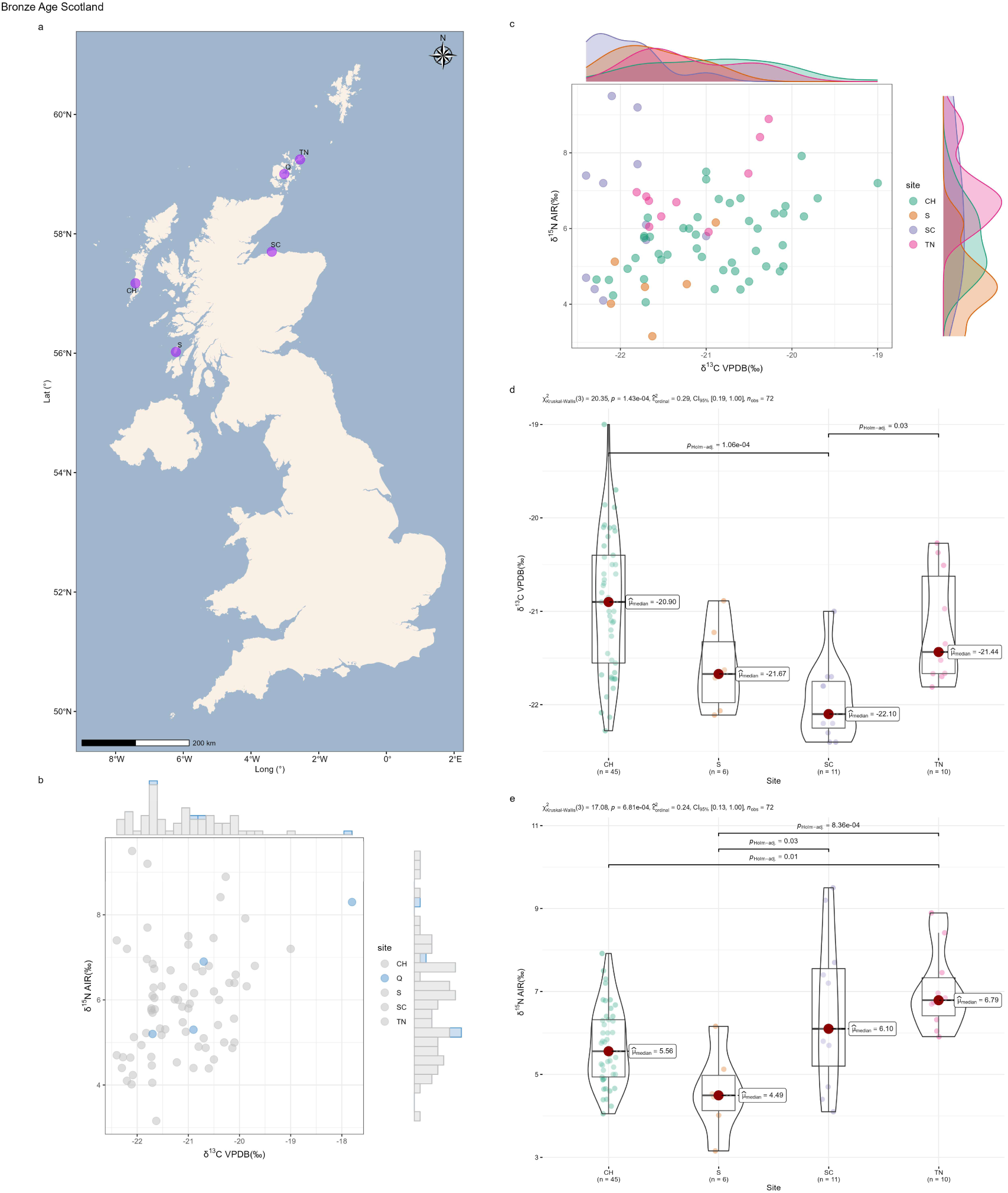
Intra-group comparison of terrestrial herbivores from Bronze Age Scotland. **a,** Distribution of sites. **b,** Scatter plot (sparsely sampled sites). **c,** Scatter plot (well-sampled sites). **d,** Statistical tests for δ^13^C (well-sampled sites). **e,** Statistical tests for δ^15^N (well-sampled sites). Site names: CH: Cladh Hallan; S: Sligenach; SC: Sculptor’s Cave; TN: Tofts Ness; Q: Quanterness.

The isotopic values of samples from Wales are shown in Fig. 14. Only one site contains a relatively large sample size, while the other two sites are represented by single samples, making meaningful comparisons within Wales impossible. Inter-group comparisons reveal that isotopic values from Wales and southern England show a high degree of overlap, whereas those from Scotland display somewhat different isotopic patterns (Fig. 15).

**Fig. 14.**
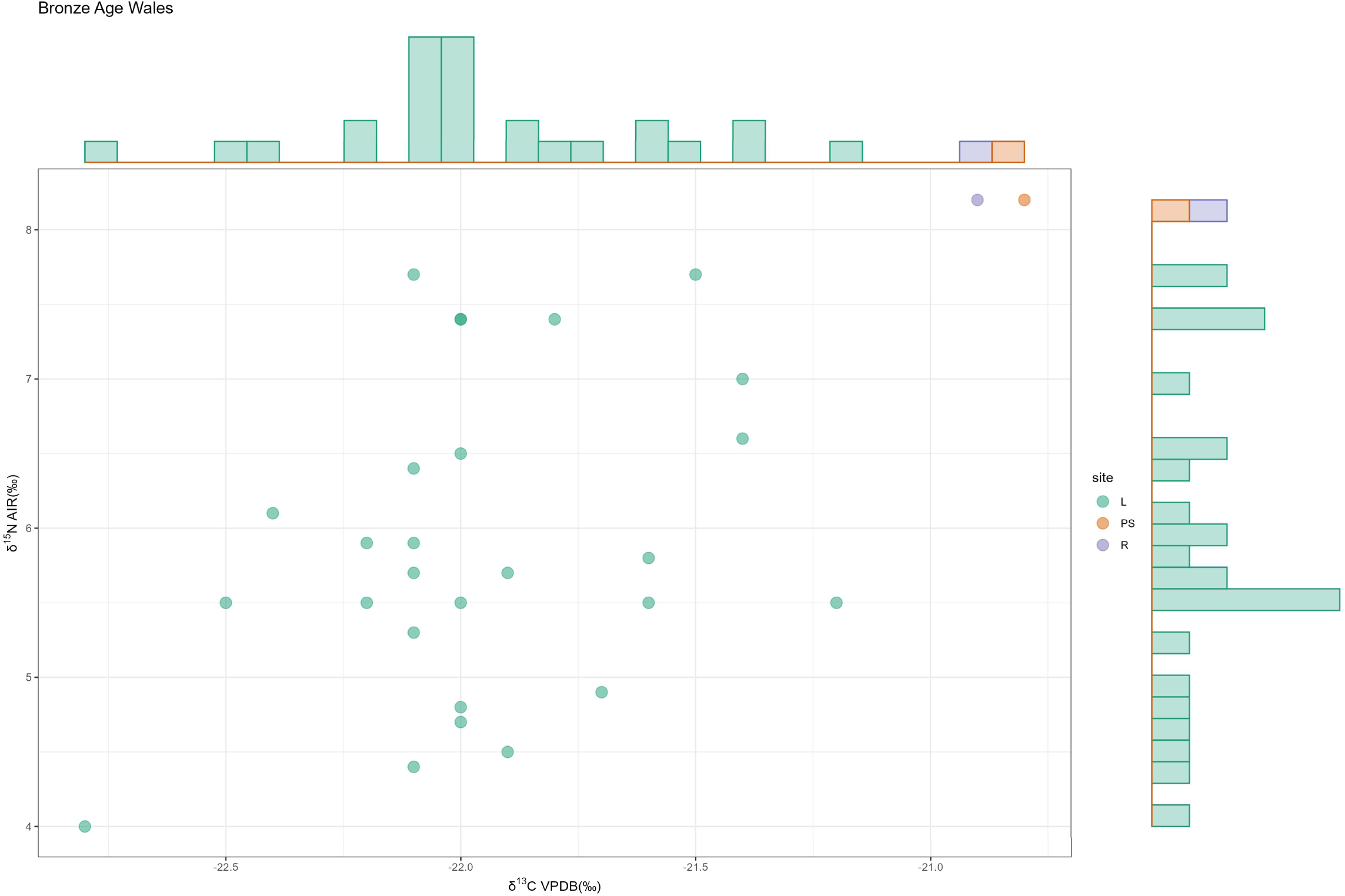
Intra-group comparison of terrestrial herbivores from Bronze Age Wales (Scatter plot). Site names: L: Llanmaes; PS: Peterstone; R: Redwick.

**Fig. 15.**
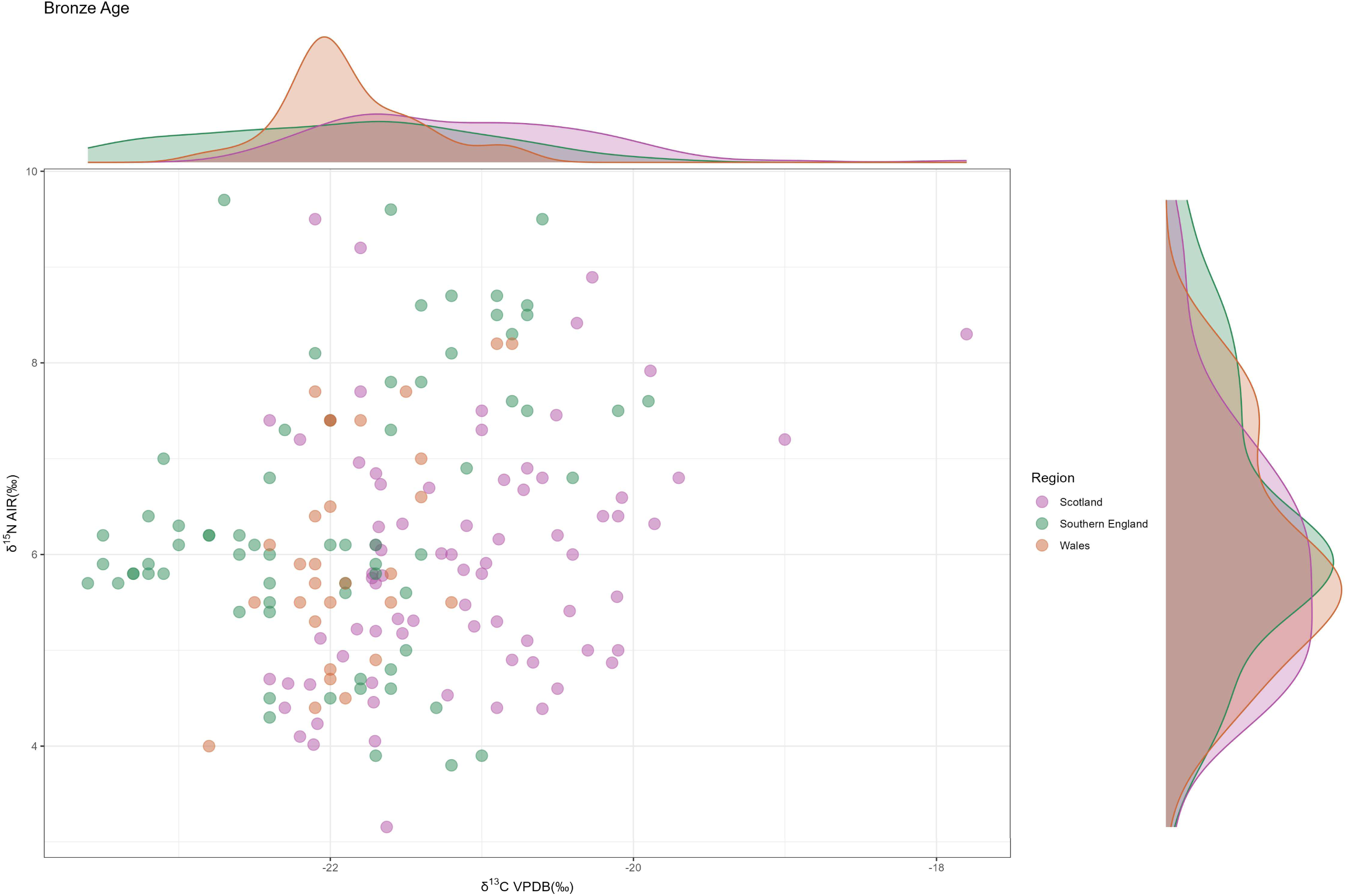
Inter-group comparison of terrestrial herbivores from the Bronze Age (Scatter plot).

The Bronze Age comparisons indicate that isotopic differences may exist among sites in southern England. In contrast, no meaningful differences are observed among sites in Scotland. Although the statistical tests yield significant results, the isotopic range of one site with a large sample size nearly encompasses those of all the others, highlighting that the statistical significance is likely driven by sample size limitations. Inter-group comparisons show that Wales falls entirely within the isotopic range of southern England, whereas Scotland displays a distinct isotopic pattern.

### Iron Age

We obtained a particularly rich dataset for terrestrial herbivores from the Iron Age southern England, comprising 34 archaeological sites. Average isotopic values and standard deviations were calculated for each site and plotted in Fig. 16a and b (sites with fewer than five samples were excluded). A notable pattern is that samples from the same site but different phases cluster together (e.g., DH1–DH4, DRH1–DRH3, SF1–SF5, NC1–NC2, Y1–Y3), indicating that minor temporal variation has little effect on isotopic values. This pattern is consistent with our earlier analyses (Fig. 3, Fig. 5, Fig. 9, Fig. 12). We therefore combined data from different phases of the same site and recalculated average values and standard deviations (Fig. 17c and Fig. 18c). In addition, isotopic values of individual samples and their associated sites are presented in Fig. 16 c-f. Data from the well-sampled sites were further analyzed by statistical tests. Results for both δ^13^C and δ^15^N are significant. The resulting p-values of post hoc tests were used to generate heat maps with dendrograms for δ^13^C and δ^15^N, respectively (Fig. 17d and Fig. 18d).

**Fig. 16.**
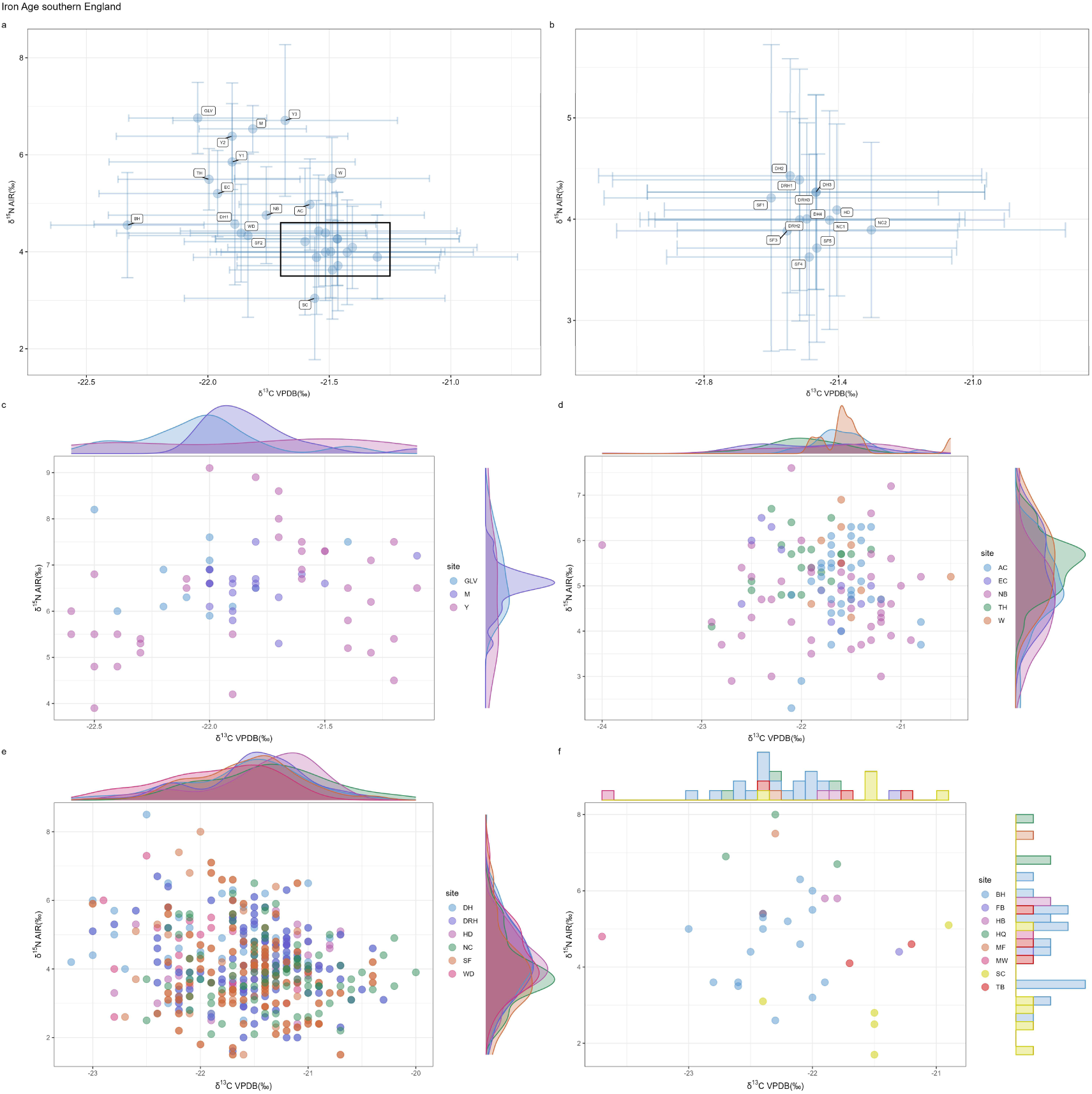
Intra-group comparison of terrestrial herbivores from Iron Age southern England (Scatter plots). **a,** Average isotopic values with standard deviations (well-sampled sites). **b,** Partial enlargement of **a**. **c,** Scatter plot 1 (well-sampled sites). **d,** Scatter plot 2 (well-sampled sites). **e,** Scatter plot 3 (well-sampled sites). **f,** Scatter plot 4 (BH, SC, and six sparsely sampled sites). Site names: AC: Alfred’s Castle; BH: Bury Hill; DH: Danebury hillfort (DH1: Danebury hillfort 1; DH2: Danebury hillfort 2; DH3: Danebury hillfort 3; DH4: Danebury hillfort 4); DRH: Danebury Ring Hillfort (DRH1: Danebury Ring Hillfort 1; DRH2: Danebury Ring Hillfort 2; DRH3: Danebury Ring Hillfort 3); EC: Eton College Rowing Course; FB: Fordington Bottom; GLV: Glastonbury Lake Village; HB: Harlyn Bay; HD: Houghton Down; HQ: Horcott Quarry; M: Marcham; MF: Manor Farm; MW: Micheldever Wood; NB: New Buildings; NC: Nettlebank Copse (NC1: Nettlebank Copse 1; NC2: Nettlebank Copse 2); SC: Segsbury Camp; SF: Sudden Farm (SF1: Sudden Farm 1; SF2: Sudden Farm 2; SF3: Sudden Farm 3; SF4: Sudden Farm 4; SF5: Sudden Farm 5); TB: Tolpuddle Ball; TH: Trethellan Head; W: Watchfield; WD: Winnall Down; Y: Yarnton (Y1: Yarnton 1; Y2: Yarnton 2; Y3: Yarnton 3). Scatter plots 1–3 separate sites with similar isotopic distributions into different panels to avoid overcrowding and improve readability.

**Fig. 17.**
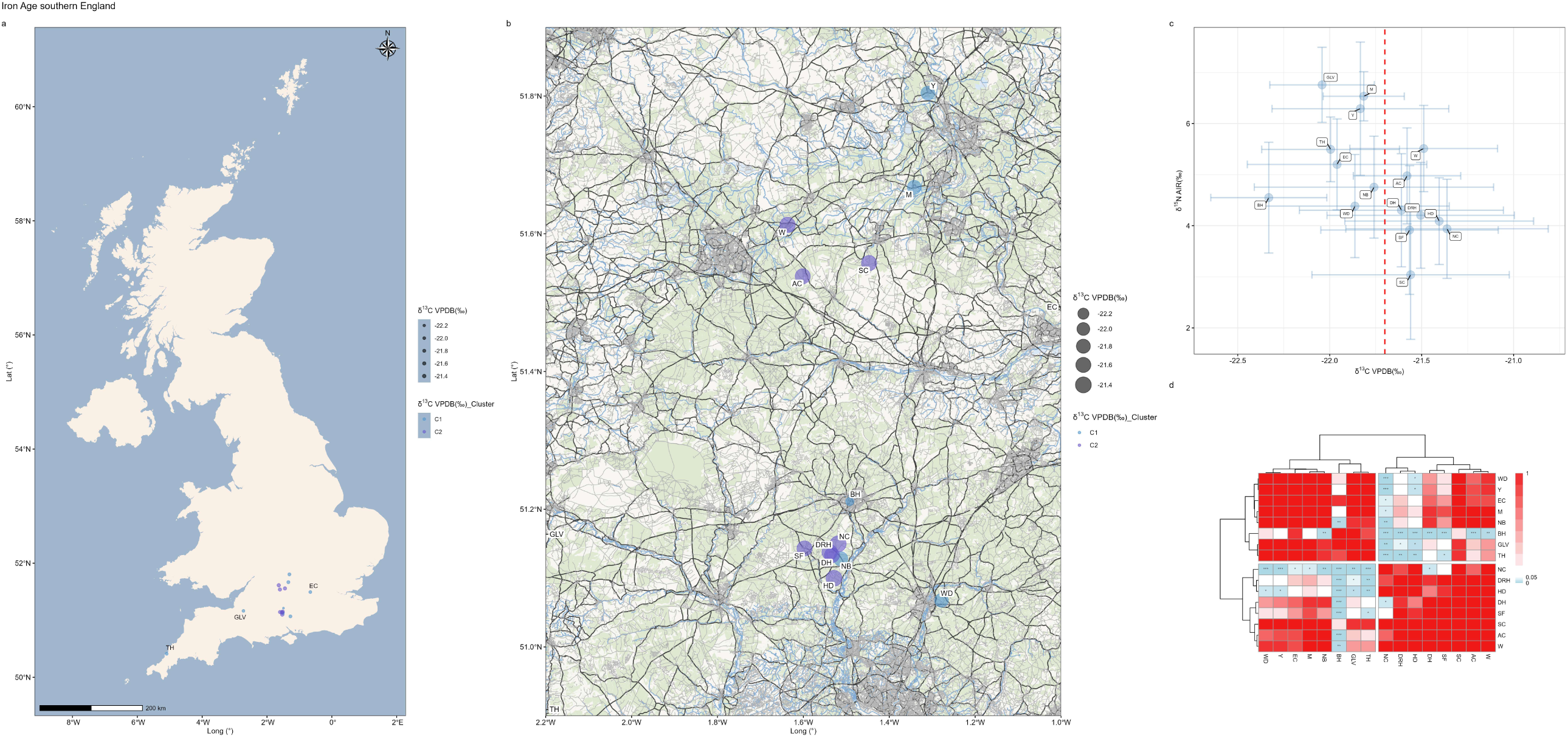
Intra-group comparison of terrestrial herbivores from Iron Age southern England (δ^13^C). **a,** Distribution of sites grouped by their δ^13^C clusters (well-sampled sites). C1 and C2 denote Cluster 1 and Cluster 2. **b,** Distribution of 13 proximate sites grouped by their δ^13^C clusters (well-sampled sites). **c,** Average isotopic values with standard deviations (well-sampled sites). **d,** The heat map with dendrogram for δ^13^C (well-sampled sites). Site names: AC: Alfred’s Castle; BH: Bury Hill; DH: Danebury hillfort; DRH: Danebury Ring Hillfort; EC: Eton College Rowing Course; GLV: Glastonbury Lake Village; HD: Houghton Down; M: Marcham; NB: New Buildings; NC: Nettlebank Copse; SC: Segsbury Camp; SF: Sudden Farm; TH: Trethellan Head; W: Watchfield; WD: Winnall Down; Y: Yarnton.

**Fig. 18.**
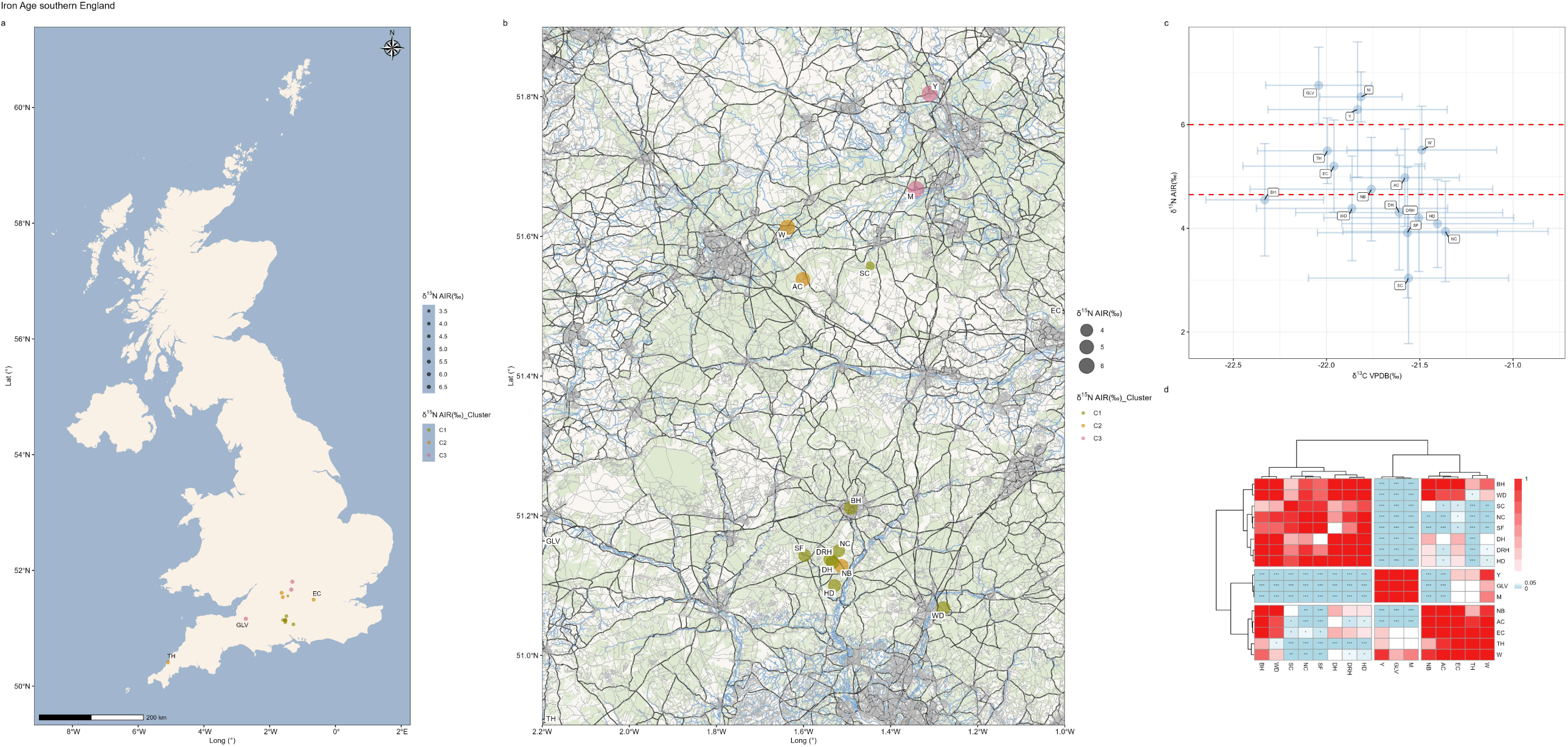
Intra-group comparison of terrestrial herbivores from Iron Age southern England (δ^15^N). **a,** Distribution of sites grouped by their δ^15^N clusters (well-sampled sites). C1–C3 denote Cluster 1 to Cluster 3. **b,** Distribution of 13 proximate sites grouped by their δ^15^N clusters (well-sampled sites). **c,** Average isotopic values with standard deviations (well-sampled sites). **d,** The heat map with dendrogram for δ^15^N (well-sampled sites). Site names: AC: Alfred’s Castle; BH: Bury Hill; DH: Danebury hillfort; DRH: Danebury Ring Hillfort; EC: Eton College Rowing Course; GLV: Glastonbury Lake Village; HD: Houghton Down; M: Marcham; NB: New Buildings; NC: Nettlebank Copse; SC: Segsbury Camp; SF: Sudden Farm; TH: Trethellan Head; W: Watchfield; WD: Winnall Down; Y: Yarnton.

The scatter plot of site averages and the heat maps reveal a highly consistent pattern. Based on δ^13^C values, sites can be preliminarily grouped into two clusters in the scatter plot: Cluster 1 (BH, EC, TH, GLV, NB, WD, M, Y) and Cluster 2 (DH, AC, SF, DRH, SC, W, HD, NC) (Fig. 17c). The dendrogram derived from the δ^13^C heat map supports this division and further clarifies the relationships among individual sites (Fig. 17d). The heat map shows that nearly all pairwise comparisons within the same cluster are statistically insignificant. Thus, the significant statistical results are attributable to pairwise comparisons between sites belonging to different clusters (i.e., sites from Cluster 1 vs. sites from Cluster 2). For δ^15^N, qualitative analysis of scatter plot suggests three clusters: Cluster 1 (SC, SF, NC, WD, DH, HD, DRH, BH), Cluster 2 (TH, W, EC, AC, NB), and Cluster 3 (Y, GLV, M) (Fig. 18c). The dendrogram supports this clustering as well (Fig. 18d). As with δ^13^C, the significant pairwise comparisons of δ^15^N also arise only from inter-cluster comparisons.

To assess whether the observed isotopic variation reflects geographic differences, we plotted the locations of these sites on maps, with point size indicating mean isotopic values and point color representing cluster membership (Fig. 17a, b and Fig. 18a, b). Geographic variation does not explain the observed δ^13^C patterns (Fig. 17a and b). As illustrated in Fig. 17b, several closely neighboring archaeological sites fall into different isotopic clusters (e.g., M and Y vs. W, AC, and SC). The most striking case is the comparison between BH and NC, which yields the lowest p-value and the largest difference in mean δ^13^C values (Fig. 17c), even though NC is the site closest to BH (Fig. 17b). This indicates that geographic proximity is not associated with isotopic similarity, contrary to the common expectation that sites located closer together should exhibit more similar isotopic values. However, this pattern can be understood by examining the δ^13^C values of each site. Among all Iron Age sites in southern England, the lowest mean δ^13^C value (BH: –22.33‰) is only 0.97‰ lower than the highest (NC: –21.36‰). Such a difference—less than one per mil—is minimal. Consequently, the results of statistical tests and clustering analyses have limited interpretive value. Given the very small differences in δ^13^C values among sites, we conclude that there are no meaningful isotopic differences across Iron Age southern England.

Unlike δ^13^C, the δ^15^N values vary more substantially among sites (Fig. 18c and d). However, the geographic structuring of δ^15^N values is also not evident. For instance, despite their close proximity, the sites Y, M, SC, AC, and W are assigned to three distinct clusters (Fig. 18b), indicating that isotopic variation can be substantial even within a relatively small region. Likewise, although NB, DRH, DH, and NC represent nearly the same site, NB (Cluster 2) falls into a different cluster from the others (Cluster 1). The underlying cause of this pattern is the extreme imbalance in sample sizes among sites. The number of sites in Iron Age southern England is very large, and the sample sizes of these sites vary considerably. For instance, SC includes only five samples, while SF is represented by 217. This uneven sampling introduces bias, which may strongly influence the statistical outcomes. Notably, sites with large sample sizes exhibit particularly wide ranges in δ^15^N values. For example, SF (217 samples) exhibits a nearly uniform distribution ranging from <2‰ to >8‰ (Fig. 16e), a span that almost encompasses the entire δ^15^N range observed at other sites. This strongly suggests that inter-site isotopic distributions may not truly differ, and that the apparent differences are largely the result of sampling bias at underrepresented sites. We therefore conclude that δ^15^N values among sites in Iron Age southern England are not significant. This raises a critical question: would isotopic variation between sites remain negligible under conditions of large and balanced sampling? At least the data analysis of Iron Age southern England, where numerous sites are represented and some are sampled extensively, supports this hypothesis.

Similar to Iron Age southern England, the dataset from Iron Age Scotland is also substantial, comprising 18 sites. Isotopic values of individual samples plotted by site are shown in Fig. 19. However, because of the large number of sites, the data are highly intertwined and difficult to interpret from these figures alone. We therefore calculated mean isotopic values and standard deviations for each site and plotted them in Fig. 20b and Fig. 21b, excluding sites with fewer than five samples (B and SC). Isotopic values from sites B and SC are presented separately in Fig. 19c, where they fall within the main cluster. Data from the well-sampled sites were further analyzed by statistical tests for both δ^13^C and δ^15^N, yielding significant results. The p-values from the post hoc tests were subsequently used to generate dendrogram heat maps for δ^13^C and δ^15^N (Fig. 20c and Fig. 21c).

**Fig. 19.**
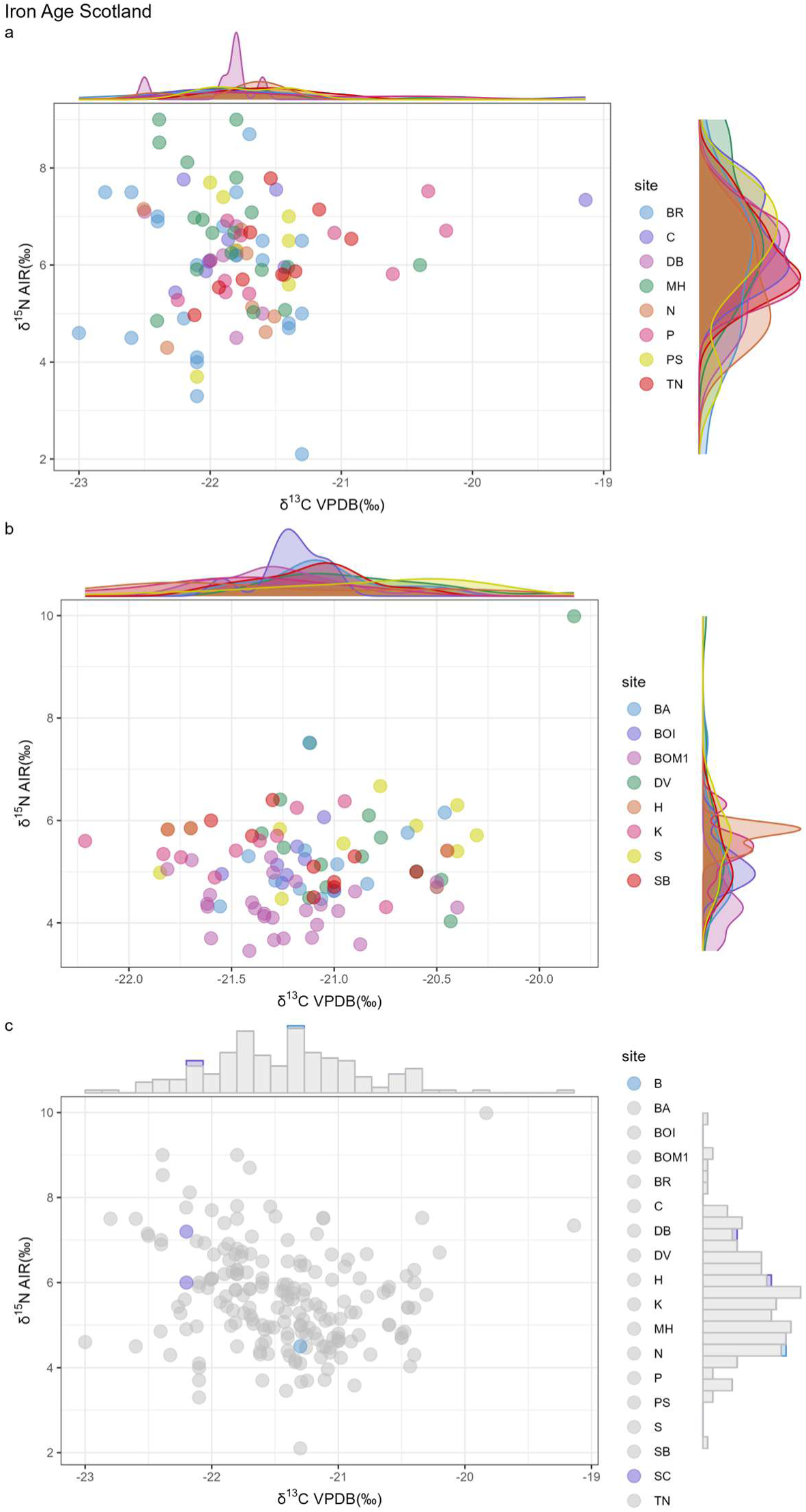
Intra-group comparison of terrestrial herbivores from Iron Age Scotland (Scatter plots). **a,** Scatter plot 1 (well-sampled sites). **b,** Scatter plot 2 (well-sampled sites). **c,** Scatter plot 3 (sparsely sampled sites). Site names: BR: Broxmouth; C: The Cairns; DB: Dryburn Bridge; MH: Mine Howe; N: Northton; P: Pool; PS: Port Seton; TN: Tofts Ness; BA: Baleshare; BOI: Bornish; BOM1: BOM1 (Western Isles); DV: Dun Vulan; H: Howe; K: Knowe o’ Skea; S: Sligenach; SB: Skara Brae; B: Bornais; SC: Sculptor’s Cave. Scatter plots 1 and 2 separate sites with similar isotopic distributions into different panels to avoid overcrowding and improve readability.

**Fig. 20.**
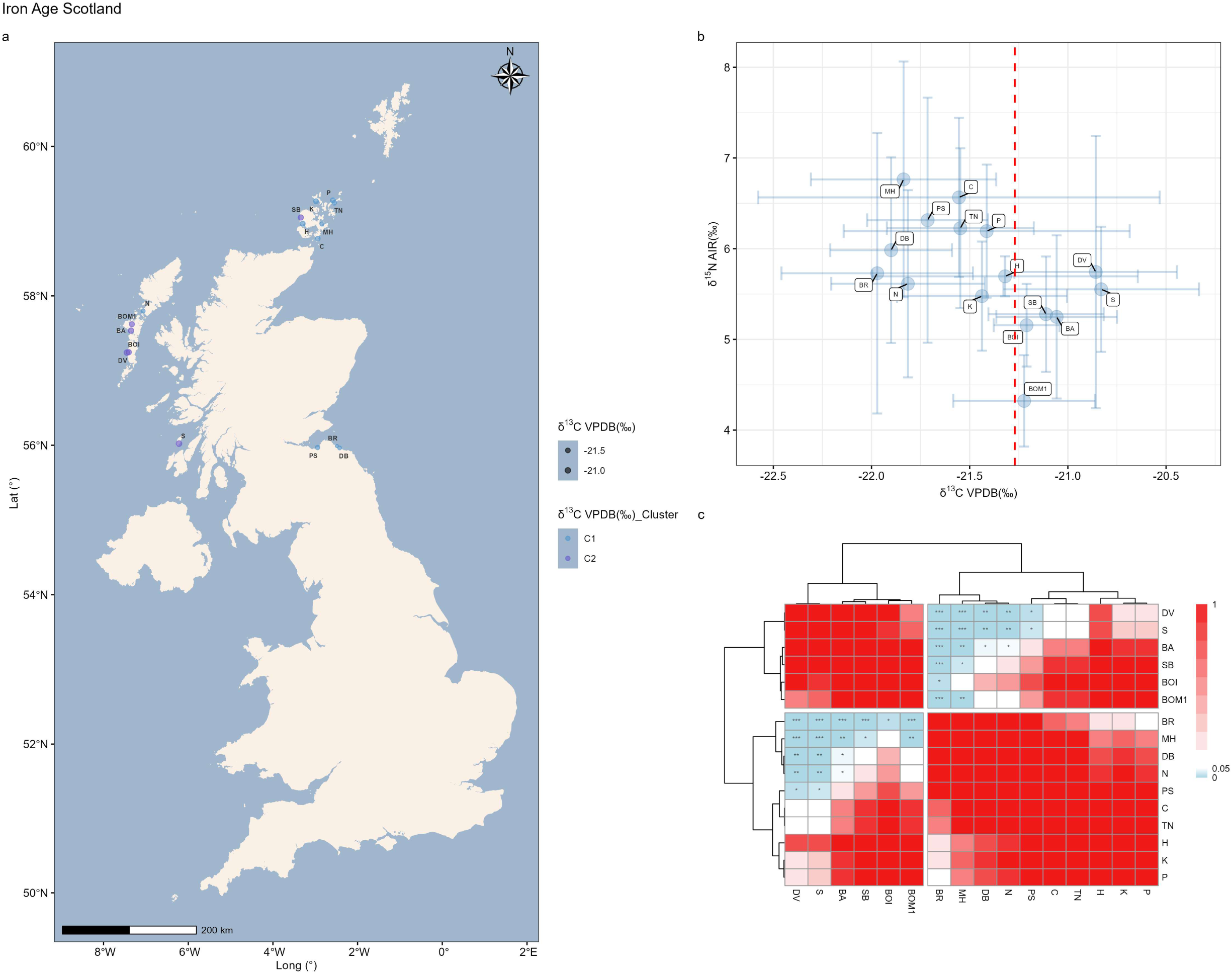
Intra-group comparison of terrestrial herbivores from Iron Age Scotland (δ^13^C). **a,** Distribution of sites grouped by their δ^13^C clusters (well-sampled sites). C1 and C2 denote Cluster 1 and Cluster 2. **b,** Average isotopic values with standard deviations (well-sampled sites). **c,** The heat map with dendrogram for δ^13^C (well-sampled sites). Site names: BR: Broxmouth; C: The Cairns; DB: Dryburn Bridge; MH: Mine Howe; N: Northton; P: Pool; PS: Port Seton; TN: Tofts Ness; BA: Baleshare; BOI: Bornish; BOM1: BOM1 (Western Isles); DV: Dun Vulan; H: Howe; K: Knowe o’ Skea; S: Sligenach; SB: Skara Brae.

**Fig. 21.**
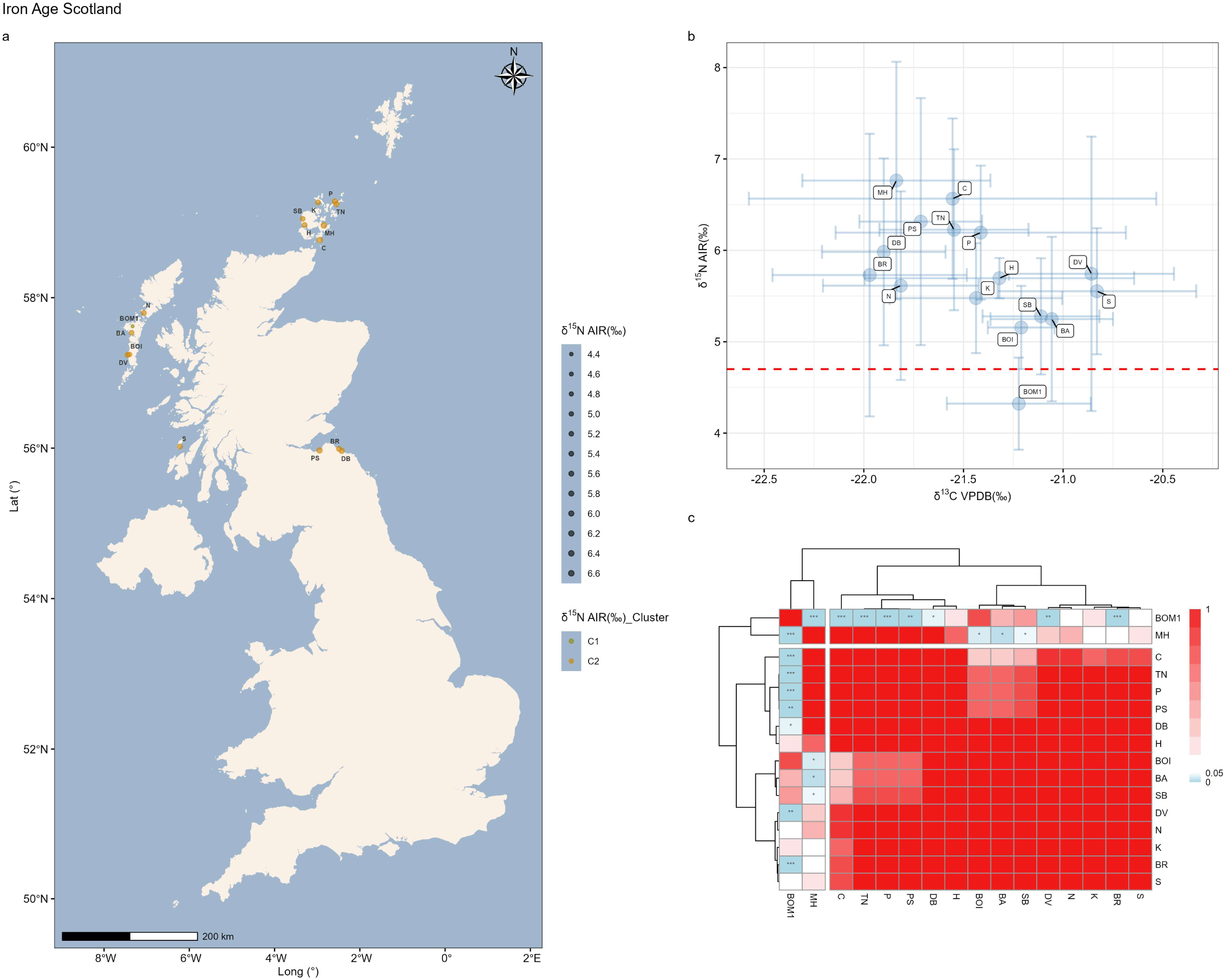
Intra-group comparison of terrestrial herbivores from Iron Age Scotland (δ^15^N). **a,** Distribution of sites grouped by their δ^15^N clusters (well-sampled sites). C1 and C2 denote Cluster 1 and Cluster 2. **b,** Average isotopic values with standard deviations (well-sampled sites). **c,** The heat map with dendrogram for δ^15^N (well-sampled sites). Site names: BR: Broxmouth; C: The Cairns; DB: Dryburn Bridge; MH: Mine Howe; N: Northton; P: Pool; PS: Port Seton; TN: Tofts Ness; BA: Baleshare; BOI: Bornish; BOM1: BOM1 (Western Isles); DV: Dun Vulan; H: Howe; K: Knowe o’ Skea; S: Sligenach; SB: Skara Brae.

For δ^13^C, qualitative analysis of the scatter plot divides the sites into two groups: Cluster 1 (BR, DB, MH, N, PS, C, TN, K, P, H) and Cluster 2 (BOM1, BOI, SB, BA, DV, S) (Fig. 20b). The dendrogram from the heat map is fully consistent with this clustering (Fig. 20c). The heat map further indicates that significant pairwise differences occur only in comparisons between sites belonging to different clusters (Cluster 1 vs. Cluster 2), whereas comparisons among sites within the same cluster show no significant differences. The geographic locations of sites, plotted with average δ^13^C values (point size) and cluster membership (point color), are shown in Fig. 20a. The map seems to suggest a spatial structuring of clusters: most sites on South Uist in the west belong to Cluster 2, while a number of eastern sites, as well as those on the Orkney Islands to the north, are largely classified as Cluster 1. Once again, we examined the isotopic values of individual sites. The mean δ^13^C values of Iron Age Scotland sites range from –20.83‰ (S) to –21.97‰ (BR), a difference of only 1.14‰, which is negligible for δ^13^C. Moreover, the standard deviation at site C is 1.02‰, almost equal to the entire range between the highest and lowest sites. This indicates that substantial overlap exists among site-level isotopic distributions, even where statistical tests detect differences in central tendency between sites. We therefore conclude that there are no meaningful differences in δ^13^C values among sites in Iron Age Scotland.

For δ^15^N, qualitative analysis of the scatter plot divides the sites into two groups: Cluster 1 (BOM1) and Cluster 2 (BOI, SB, BA, K, S, BR, N, H, DV, DB, TN, P, PS, C, MH) (Fig. 21b). The dendrogram, however, shows a slightly different pattern, clustering MH with BOM1 (Fig. 21c). This is inconsistent with the scatter plot, which clearly shows BOM1 as the site with the lowest δ^15^N value and MH as the site with the highest. Thus, the dendrogram result is likely spurious. Pairwise comparisons indicate that the significant statistical result is driven primarily by BOM1 (Fig. 21c). When BOM1 is excluded, all pairwise comparisons become insignificant. Yet, given that BOM1 is situated on South Uist (Fig. 21a), its isotopic characteristics would not be expected to differ markedly from those of adjacent sites on the same island. Accordingly, we interpret the anomalous pattern observed at BOM1 as a consequence of disparities in sample size. We therefore conclude that δ^15^N values show no meaningful differences among Iron Age Scotland sites.

We then compared isotopic values across the Iron Age combined groups (southern England, Scotland, and northern England) (Fig. 22). With the exception of a small subset of samples from Scotland, all other data are intermixed. Notably, all samples from northern England fall within the clusters defined by southern England. The isotopic distribution of northern England, represented entirely by a single site with a large sample size (Wetwang Slack), closely resembles that of southern England, which is based on 28 sites. Together with the case of Iron Age southern England, where SF (n = 217) encompassed nearly the entire isotopic range of all other sites, this raises an important question: if the sample size from a single site is sufficiently large, could its isotopic signature approximate that of an entire region? This finding suggests that constructing baselines from a single site, or from only a few sites, may introduce considerable sampling bias. Naturally, given the scarcity of archaeological samples, this analysis cannot be taken as a definitive conclusion but should instead be regarded as a working hypothesis.

**Fig. 22.**
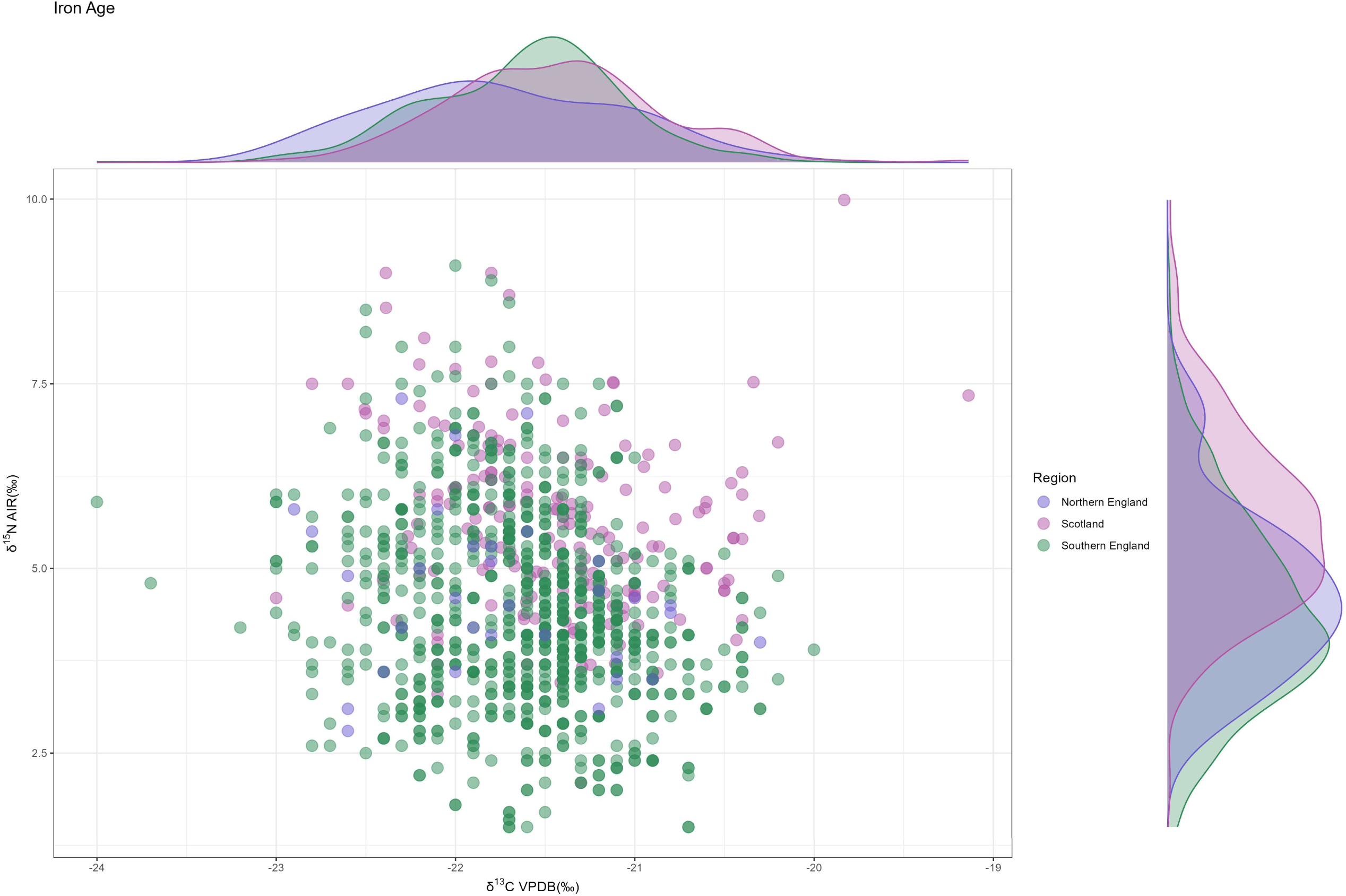
Inter-group comparison of terrestrial herbivores from the Iron Age (Scatter plot).

### Roman Period/Roman Iron Age

For the Roman period, intra-group comparisons can be made for northern and southern England. Results for northern England are shown in Fig. 23. Only WS and TR exhibit relatively large differences in δ^15^N values (p = 0.02). However, as the sample sizes at these sites are all below 20, sampling bias is likely, and such a result is therefore unsurprising.

**Fig. 23.**
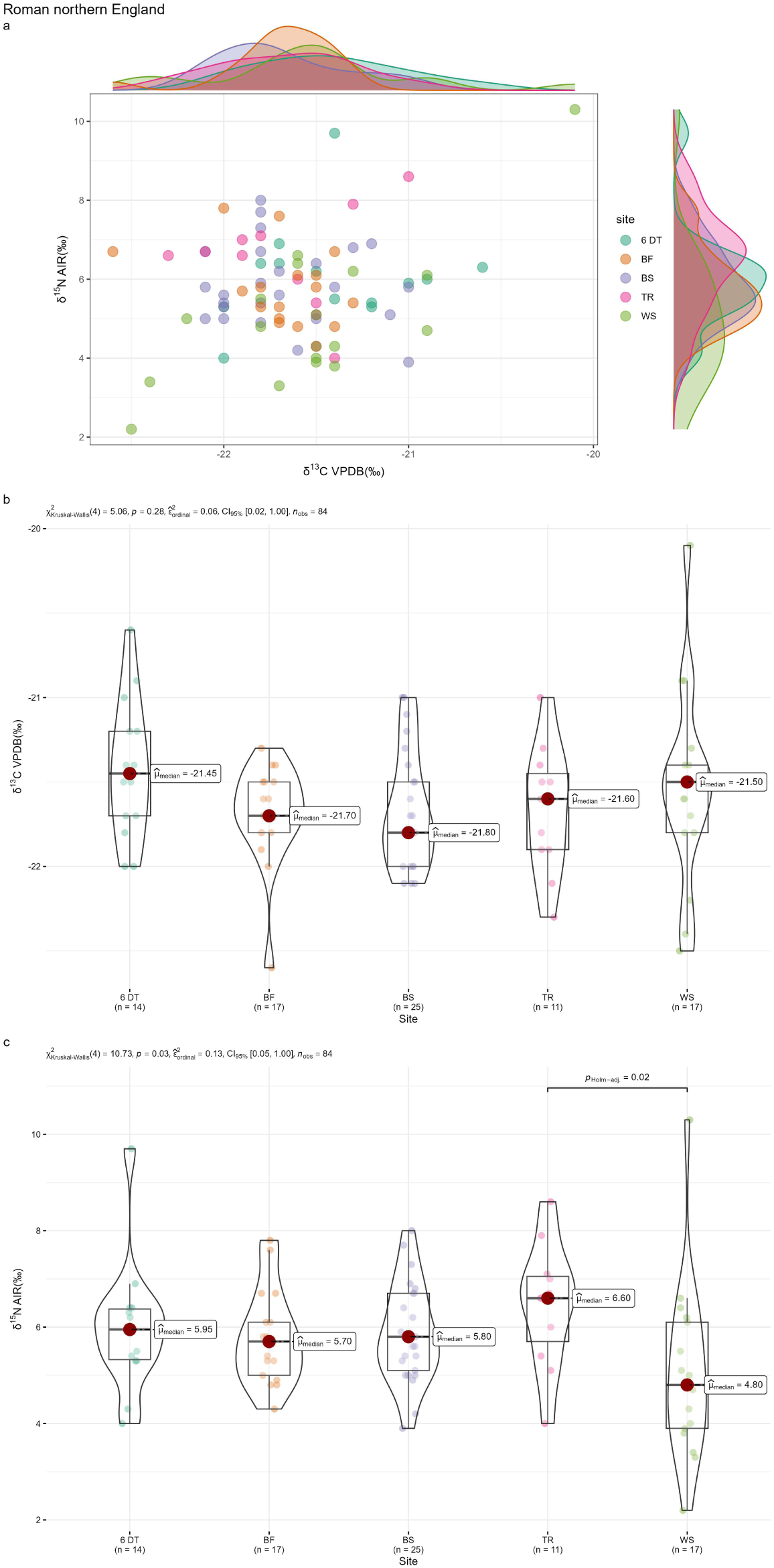
Intra-group comparison of terrestrial herbivores from Roman northern England. **a,** Scatter plot. **b,** Statistical tests for δ^13^C. **c,** Statistical tests for δ^15^N. Site names: 6 DT: 6 Driffield Terrace; BF: Bainesse Farm; BS: Blossom Street; TR: Tanner Row; WS: Wetwang Slack.

For southern England, mean isotopic values with standard deviations for the well-sampled sites are shown in Fig. 25d and Fig. 26d. Six sites (EC, P, R, LR, HQ, Y) exhibit relatively similar values, whereas three sites (L, SG, M) display wider variation. Isotopic values of individual samples from well-sampled sites are presented in Fig. 24a and b, while those from sites with small sample sizes are shown in Fig. 24c. The latter all fall within the range defined by the well-sampled sites.

**Fig. 24.**
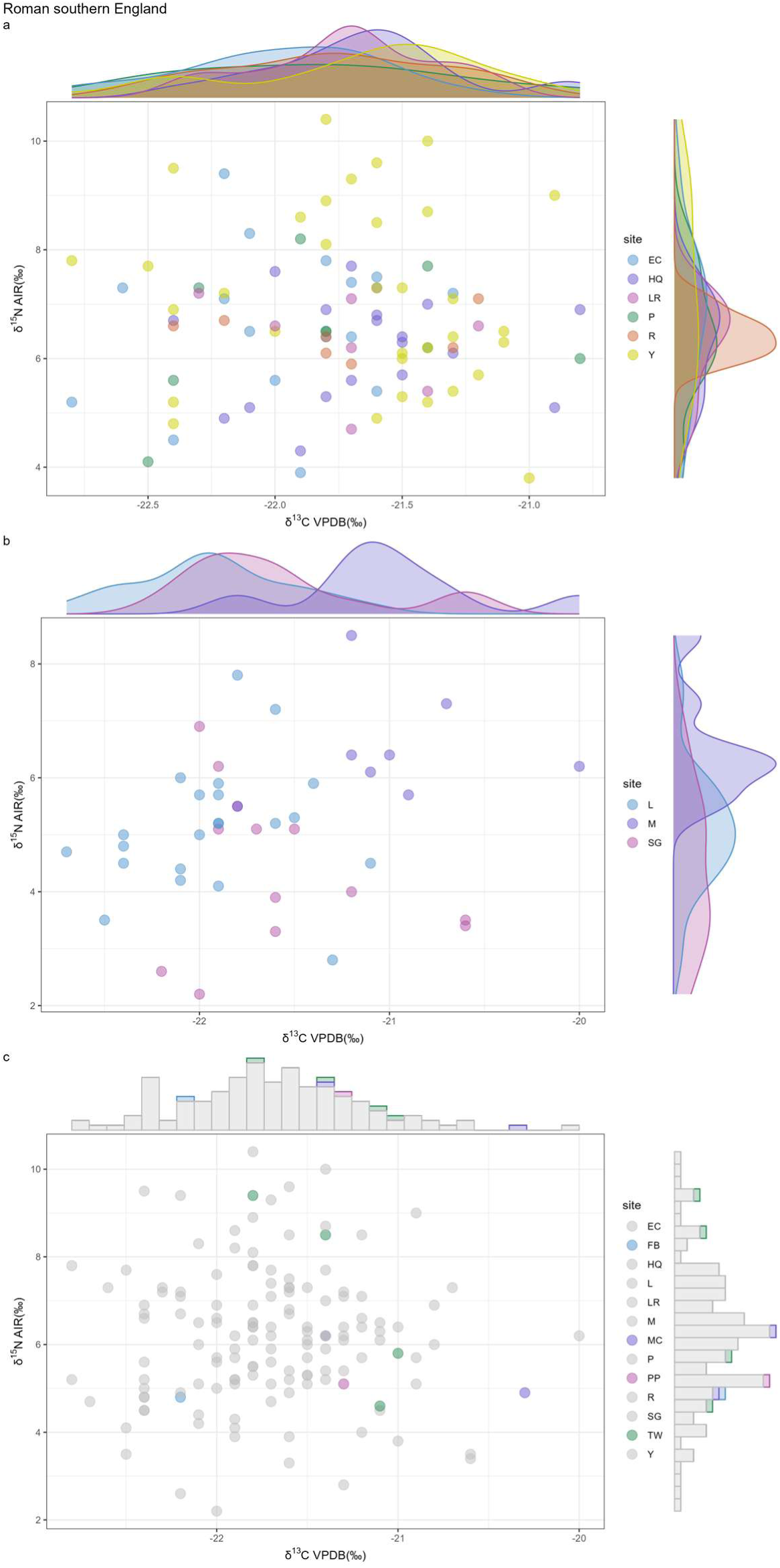
Intra-group comparison of terrestrial herbivores from Roman southern England (Scatter plots). **a,** Scatter plot 1 (well-sampled sites). **b,** Scatter plot 2 (well-sampled sites). **c,** Scatter plot 3 (sparsely sampled sites). Site names: EC: Eton College Rowing Course; HQ: Horcott Quarry; LR: 120-122 London Road; P: 1 Poultry; R: Radley; Y: Yarnton; L: Lankhills; M: Monkton; SG: Staple Gardens; FB: Fordington Bottom; MC: Maiden Castle Road; PP: Poundbury Pipeline; TW: Tubney Wood Quarry.

**Fig. 25.**
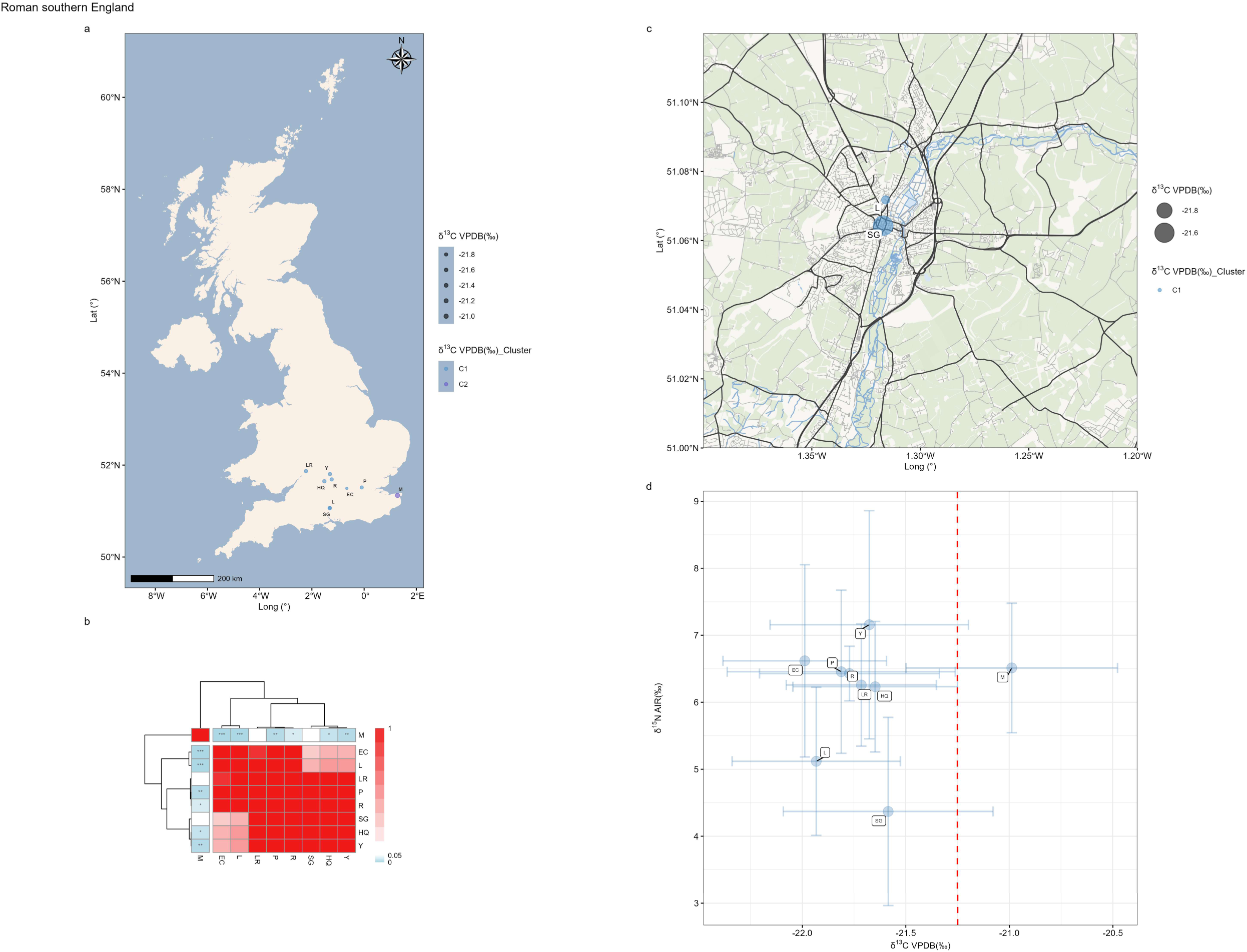
Intra-group comparison of terrestrial herbivores from Roman southern England (δ^13^C). **a,** Distribution of sites grouped by their δ^13^C clusters (well-sampled sites). C1 and C2 denote Cluster 1 and Cluster 2. **b,** The heat map with dendrogram for δ^13^C (well-sampled sites). **c,** Distribution of two proximate sites. **d,** Average isotopic values with standard deviations (well-sampled sites). Site names: EC: Eton College Rowing Course; HQ: Horcott Quarry; L: Lankhills; LR: 120-122 London Road; M: Monkton; P: 1 Poultry; R: Radley; SG: Staple Gardens; Y: Yarnton.

**Fig. 26.**
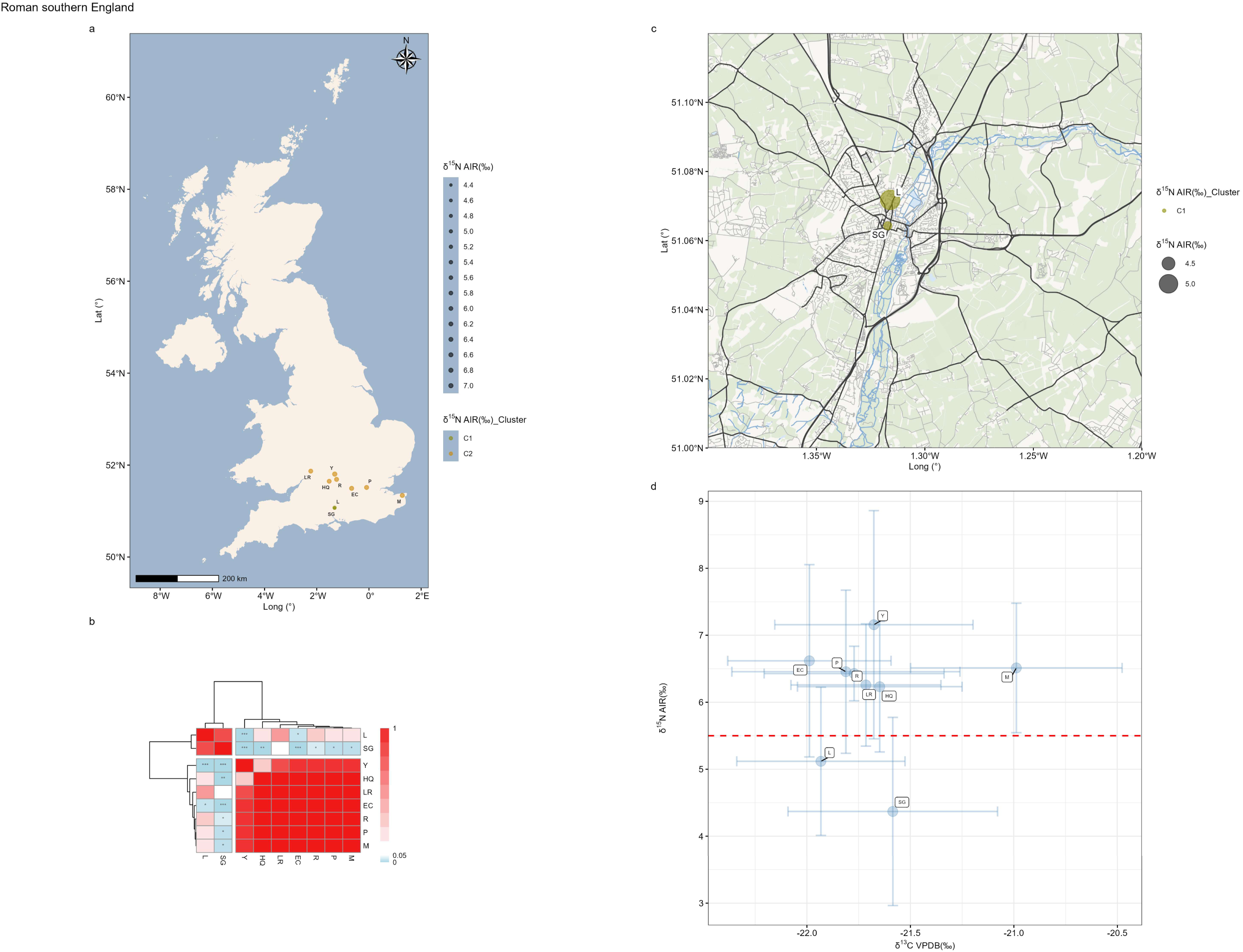
Intra-group comparison of terrestrial herbivores from Roman southern England (δ^15^N). **a,** Distribution of sites grouped by their δ^15^N clusters (well-sampled sites). C1 and C2 denote Cluster 1 and Cluster 2. **b,** The heat map with dendrogram for δ^15^N (well-sampled sites). **c,** Distribution of two proximate sites. **d,** Average isotopic values with standard deviations (well-sampled sites). Site names: EC: Eton College Rowing Course; HQ: Horcott Quarry; L: Lankhills; LR: 120-122 London Road; M: Monkton; P: 1 Poultry; R: Radley; SG: Staple Gardens; Y: Yarnton.

The statistical tests with post hoc tests was conducted for δ^13^C and δ^15^N using the isotopic data of well-sampled sites, and the resulting p-values were visualized as heat maps with dendrograms (Fig. 25b and Fig. 26b). In both δ^13^C and δ^15^N, the sites can be divided into two clusters (Cluster 1 and Cluster 2), with significant results arising exclusively from inter-cluster comparisons. Among all sites, M shows a significant difference in δ^13^C, while L and M differ significantly in δ^15^N. Mapping site locations (Fig. 25a, c and Fig. 26a, c) reveals that L, SG, and M are geographically more distant from other sites, suggesting that the isotopic variation may reflect geographic patterning.

However, the sample sizes of these three sites are relatively small. The site with the largest sample size in Roman southern England is Y (n=33). As shown in the scatter plots (Fig. 24a, b), the δ^13^C values at Y are broadly distributed between –21‰ and – 22.5‰, while the δ^15^N values span evenly from about 4‰ to 10‰. This distribution essentially encompasses most of the isotopic variation observed at L, M, and SG. Once again, the isotopic variation observed at sites with smaller sample sizes falls within the clusters defined by those with larger sample sizes, suggesting that the apparent differences are more likely due to insufficient and uneven sampling rather than genuine isotopic divergence. We therefore conclude that sites within the same Roman combined group do not exhibit significant isotopic differentiation for terrestrial herbivores.

When isotopic values were compared across the combined groups (Fig. 27), those from northern and southern England were intermixed. The few samples from Wales (n = 2) and Scotland (n = 1) also fall within the clusters defined by England. This pattern is consistent with the results of inter-group comparisons in earlier periods. It further supports that the significant results observed within the combined groups are likely due to insufficient sample sizes. Indeed, even samples from two geographically distant regions (northern and southern England) show remarkably similar isotopic distributions when the sample sizes are sufficiently large (Fig. 27).

**Fig. 27.**
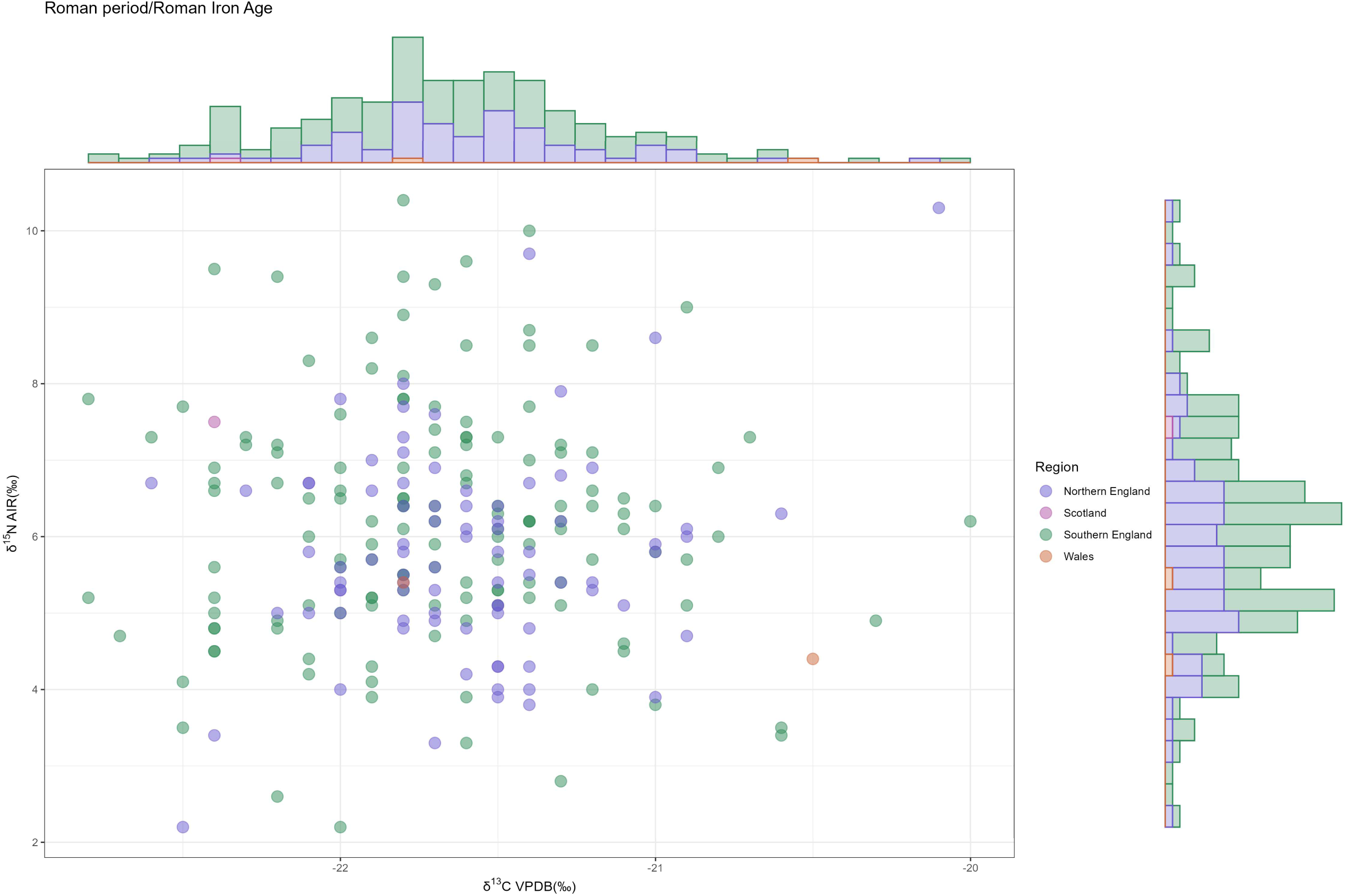
Inter-group comparison of terrestrial herbivores from the Roman period/Roman Iron Age (Scatter plot).

### Early Medieval Period

For the Early Medieval period, intra-group comparisons can be made for three regions: northern England, Scotland, and southern England. The scatter plot for northern England shows that the isotopic distributions of its three sites each display distinct characteristics (Fig. 28b). The statistical test indicates that the only low p-values are from post hoc tests of δ^15^N between BL and F (Fig. 28d). Nevertheless, the sample sizes are very small (BL: 6, F: 7, M: 5). Such limited sample sizes prevent us from drawing firm conclusions.

**Fig. 28.**
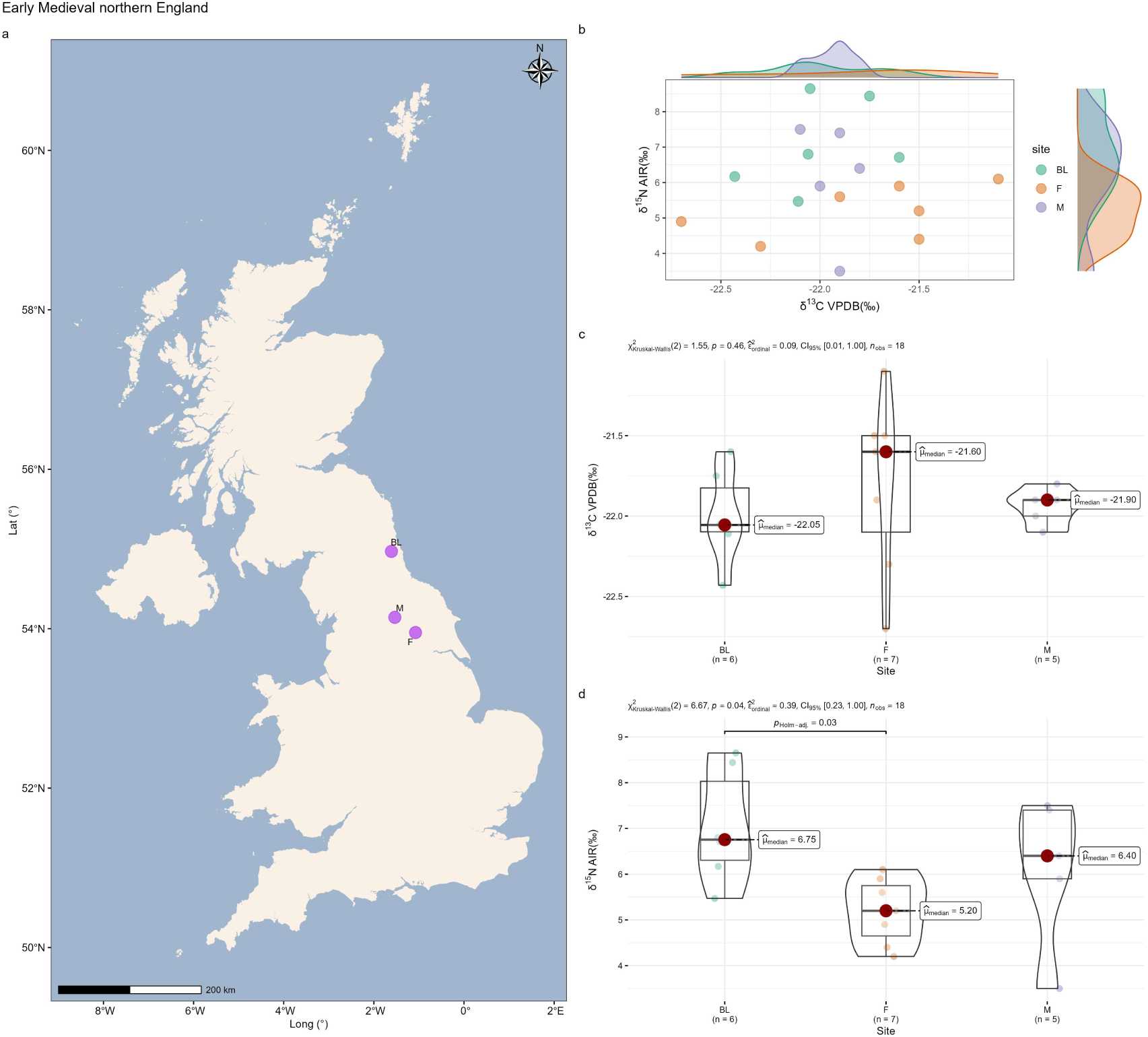
Intra-group comparison of terrestrial herbivores from Early Medieval northern England. **a,** Distribution of sites. **b,** Scatter plot. **c,** Statistical tests for δ^13^C. **d,** Statistical tests for δ^15^N. Site names: BL: Blackgate; F: Fishergate; M: Masham.

For Scotland, the statistical tests are significant for both δ^13^C and δ^15^N (Fig. 29d and e). However, the scatter plot shows that most samples from EB (n = 19) and BO (n = 14) fall within the cluster defined by P (n = 38), with only five exceptions (Fig. 29c). In terms of δ^15^N values, all BO and EB samples fall within the range defined by P. Although δ^13^C values show a slightly different pattern, the differences in mean and median values among the three sites are within 0.5‰ and can be considered negligible. This suggests that, if sample sizes were sufficiently large and balanced across sites, isotopic characteristics might prove similar. Isotopic values from sites with small sample sizes are shown in Fig. 29b. Most samples from J fall within the main cluster.

**Fig. 29.**
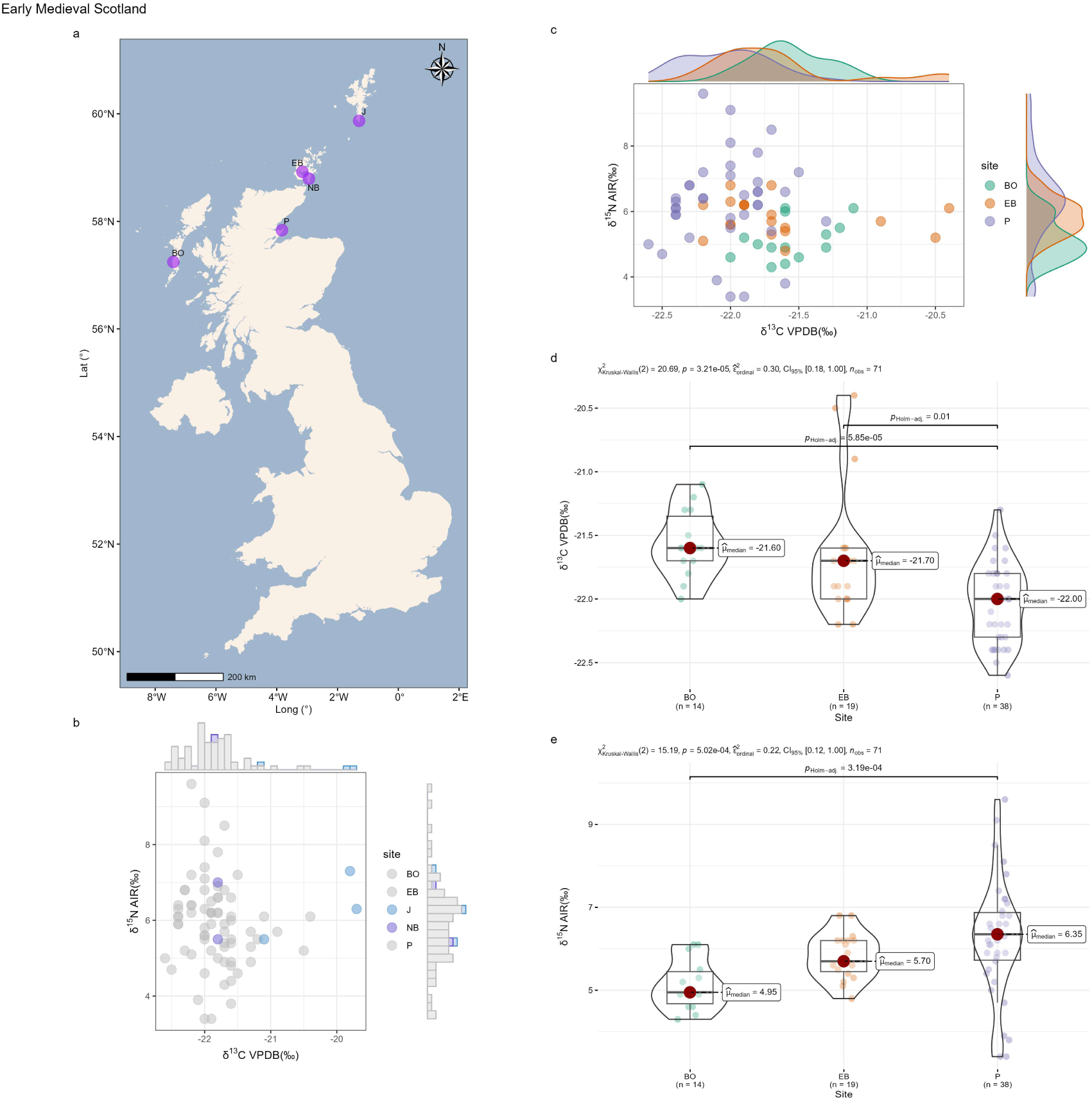
Intra-group comparison of terrestrial herbivores from Early Medieval Scotland. **a,** Distribution of sites. **b,** Scatter plot (sparsely sampled sites). **c,** Scatter plot (well-sampled sites). **d,** Statistical tests for δ^13^C (well-sampled sites). **e,** Statistical tests for δ^15^N (well-sampled sites). Site names: BO: Bornais; EB: Earl’s Bu; P: Portmahomack; J: Jarlshof; NB: Site of Newark Bay.

Sites in southern England display a comparable isotopic pattern. The isotopic values of all sites are intermixed (Fig. 30b). Only one post hoc test yields a low p-value (HQ vs. L: δ^15^N) (Fig. 30d), but because the distributions and mean values of these two sites are highly similar, the low p-value cannot be considered meaningful. With relatively large sample sizes (HQ, n = 50; L, n = 58), both sites show δ^13^C values ranging from –23‰ to –20.5‰ and δ^15^N values from 2‰ to 8‰. These broad and even distributions nearly cover the full isotopic variation observed at other sites, matching the patterns found in southern England during the Iron Age (site: SE) and the Roman period (site: Y). Our results indicate that, in the Early Medieval period, herbivore isotopic values were largely consistent across sites within each combined group.

**Fig. 30.**
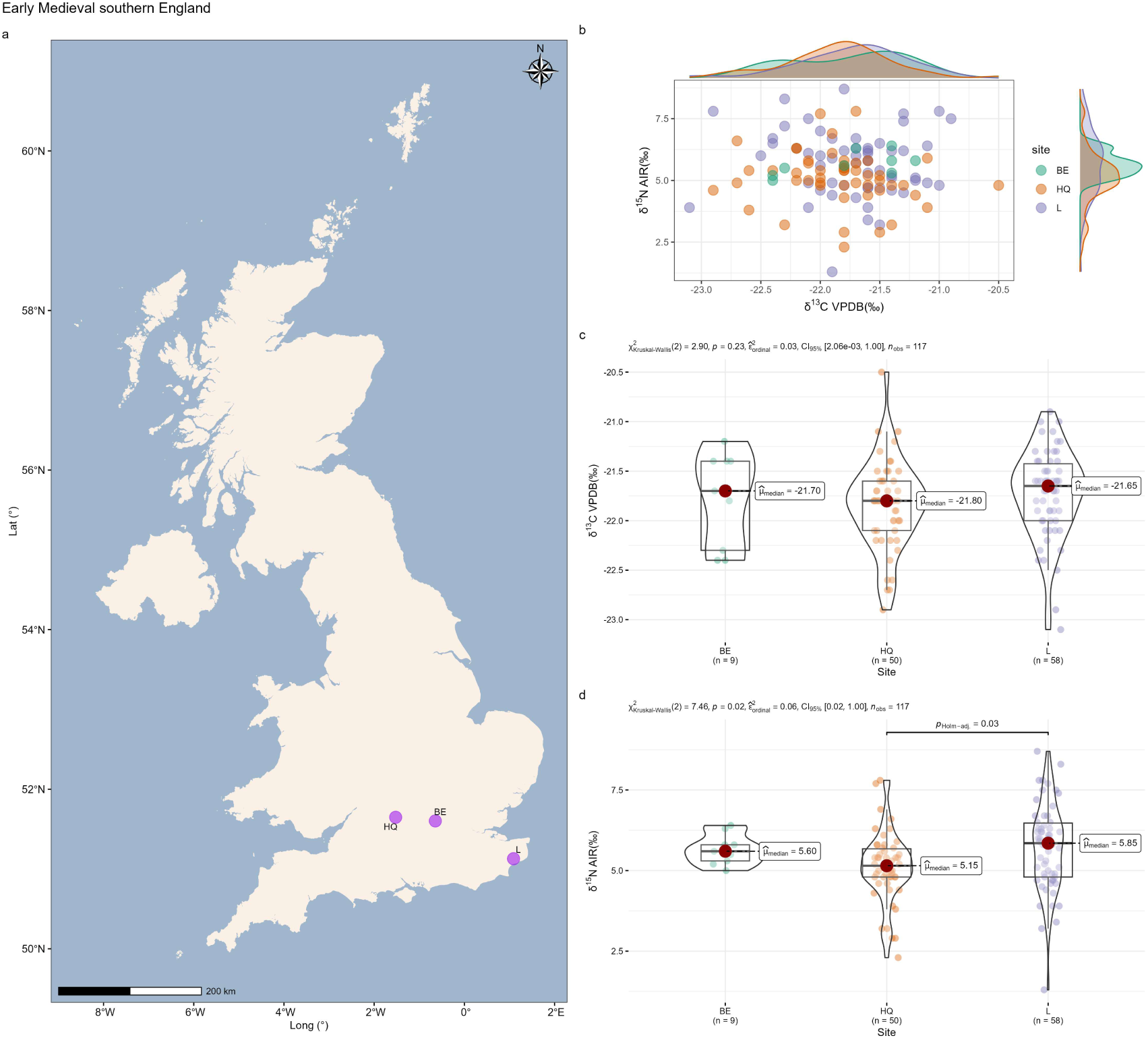
Intra-group comparison of terrestrial herbivores from Early Medieval southern England. **a,** Distribution of sites. **b,** Scatter plot. **c,** Statistical tests for δ^13^C. **d,** Statistical tests for δ^15^N. Site names: BE: Beaconsfield; HQ: Horcott Quarry; L: Lyminge.

Inter-group comparisons indicate that, except for a few samples from Scotland, nearly all regions with smaller sample sizes fall within the isotopic range defined by southern England, which has the largest sample size (Fig. 31). This once again suggests, albeit indirectly, that when sample sizes are sufficiently large, isotopic differences between sites tend to be minimal.

**Fig. 31.**
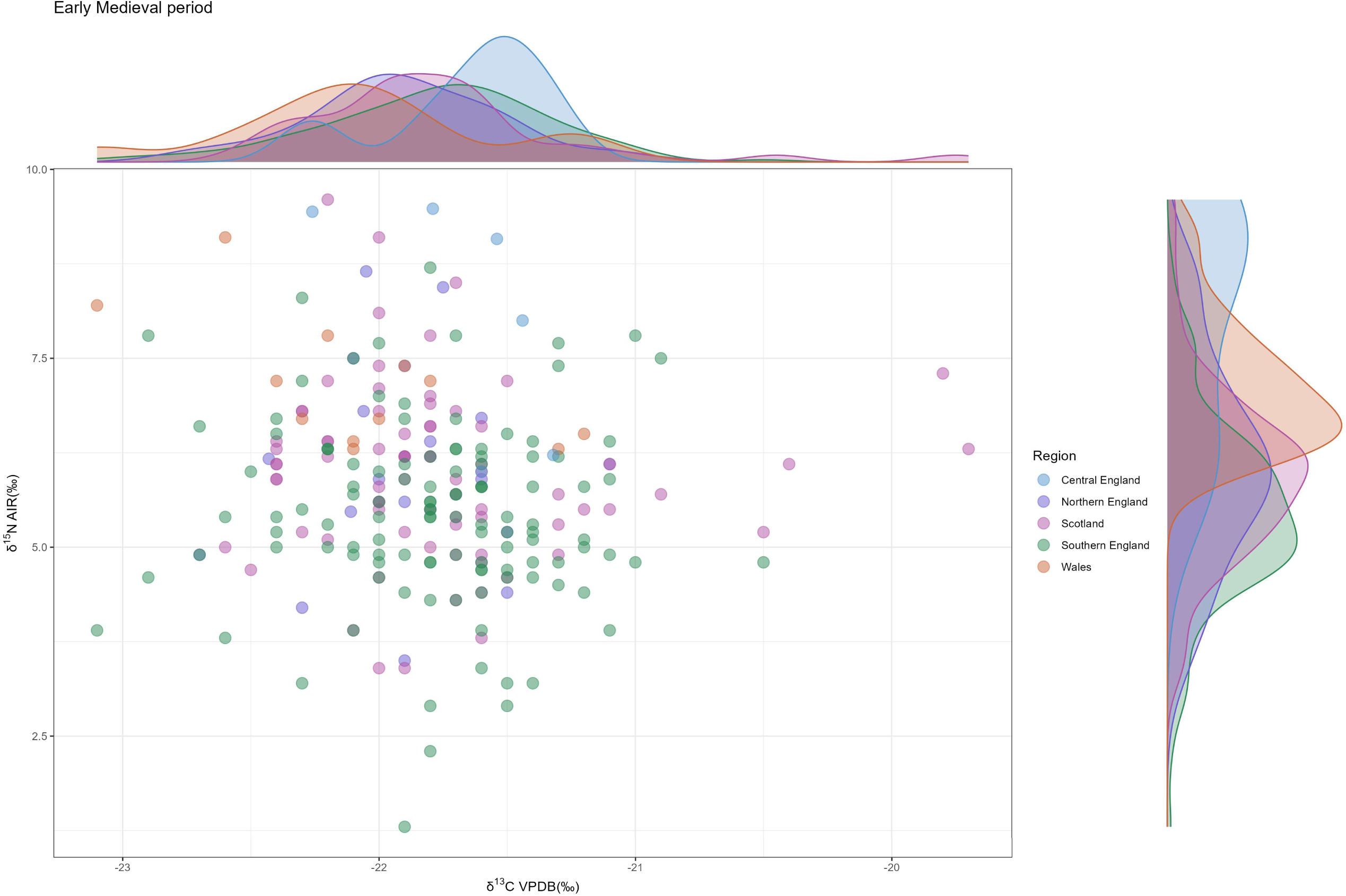
Inter-group comparison of terrestrial herbivores from the Early Medieval period (Scatter plot).

### Later Medieval and Post Medieval Periods

We conducted three intra-group comparisons for the Later Medieval period: southern England, northern England, and Scotland. Later Medieval southern England is affected by small and uneven sample sizes, similar to several combined groups in earlier periods. The scatter plot shows that most of the isotopic values of AS (n = 10) and OC (n = 10) fall within the cluster defined by SA (n = 20) (Fig. 32b). Post hoc tests are largely insignificant, with the exception of AS versus SA for δ^13^C (Fig. 32c). As noted above, this difference is likely attributable to small and uneven sample sizes. This point is further illustrated by the geographic locations of the sites. All three sites are located in close proximity (within the Oxford city center) and can effectively be regarded as a single site (Fig. 32a), which is unlikely to exhibit genuine isotopic differences.

**Fig. 32.**
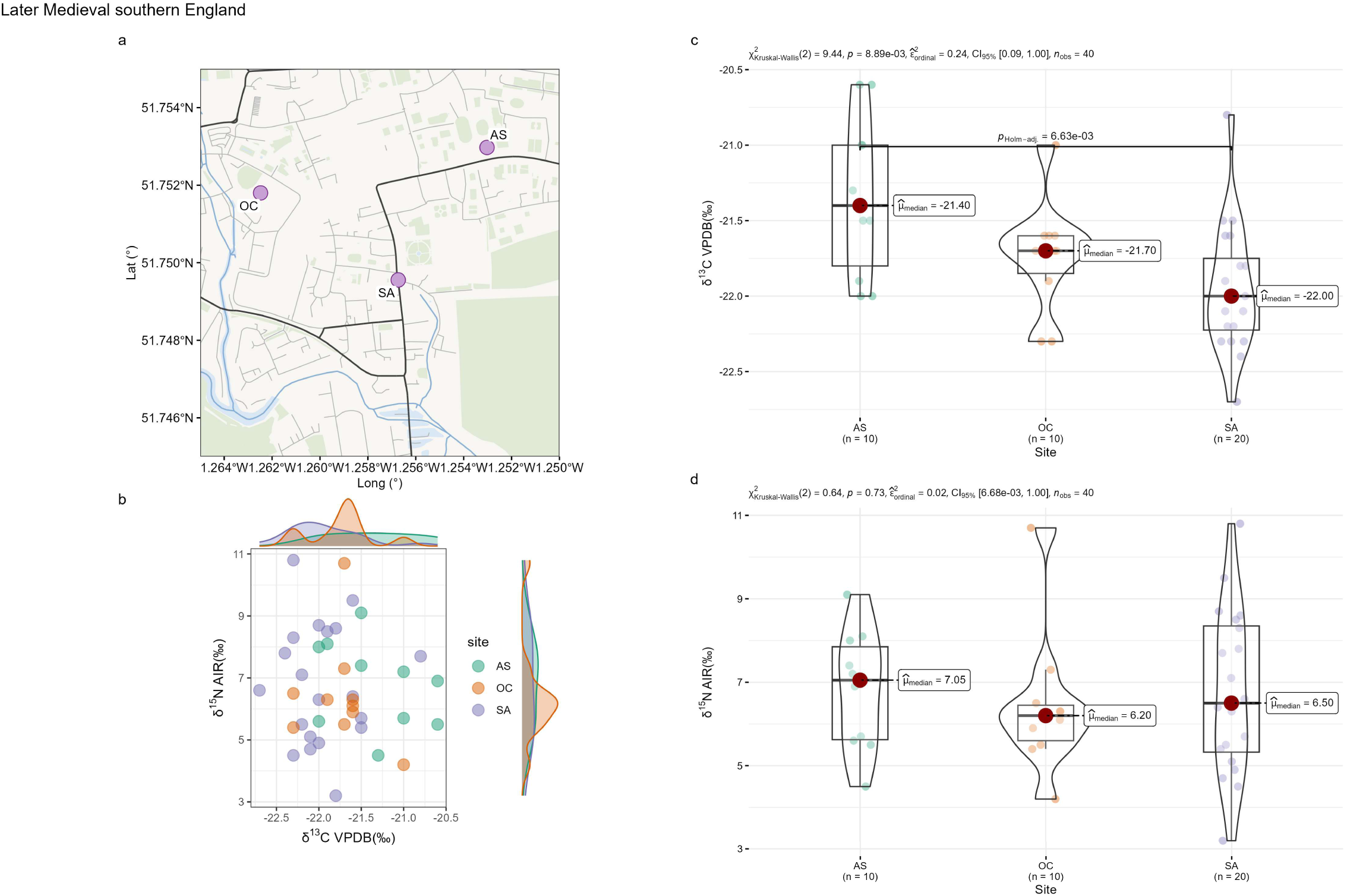
Intra-group comparison of terrestrial herbivores from Later Medieval southern England. **a,** Distribution of sites. **b,** Scatter plot. **c,** Statistical tests for δ^13^C. **d,** Statistical tests for δ^15^N. Site names: AS: All Saints; OC: Oxford Castle; SA: St Aldate’s PS.

For northern England, BL is the only outlier (Fig. 33c and d). Except for its slightly lower δ^13^C values, all other sites show broadly similar isotopic distributions. BL even yields significant statistical differences when compared with its nearest neighbor, site F (Fig. 33a), which is unexpected. As BL comprises only six samples, the observed significance is very likely spurious. Isotopic values from the remaining sites with small sample sizes fall within the main cluster (Fig. 33b).

**Fig. 33.**
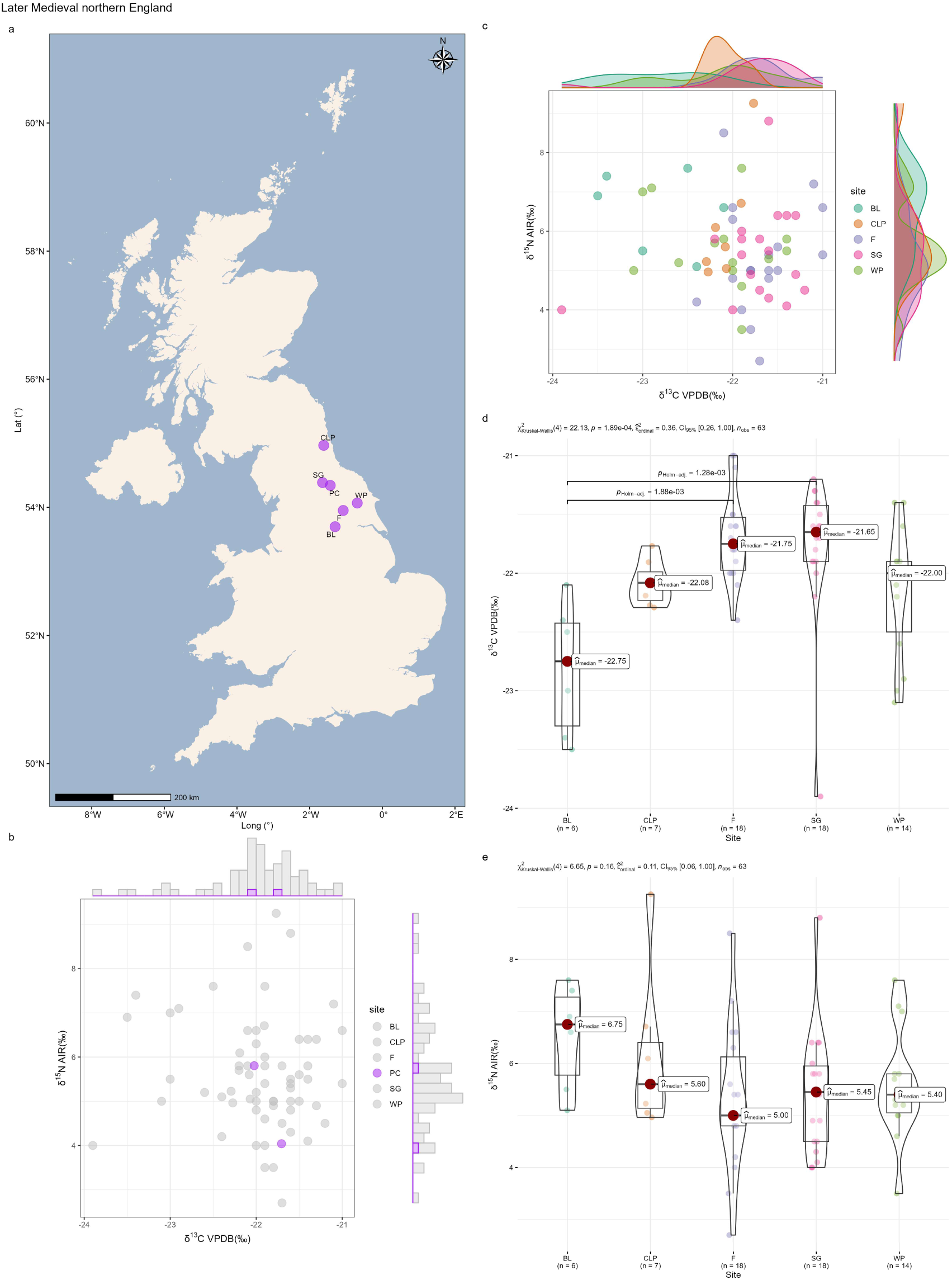
Intra-group comparison of terrestrial herbivores from Later Medieval northern England. **a,** Distribution of sites. **b,** Scatter plot (sparsely sampled sites). **c,** Scatter plot (well-sampled sites). **d,** Statistical tests for δ^13^C (well-sampled sites). **e,** Statistical tests for δ^15^N (well-sampled sites). Site names: BL: Box Lane; CLP: Clavering Place; F: Fishergate; SG: St. Giles; WP: Wharram Percy; PC: Priory Close.

The results for Scotland are more complex. Sites FF and P differ significantly from the others. The significant statistical results are driven by comparisons of sites FF and P with the other sites (Fig. 34d and e). Isotopic values from FF and P are highly similar to each other in both δ^13^C and δ^15^N, but distinct from the other five sites. Sparsely sampled sites in Scotland also include outliers (site NB) (Fig. 34b). No clear geographic pattern emerges (Fig. 34a). In Scotland, we expect FF and P to cluster together and be distinct from other sites. However, this is not the case. Apart from FF and P, the other sites with similar isotopic values also show no geographic clustering. The sparsely sampled site NB, which contains outlier samples, lies very close to other sites, indicating no geographic distinctiveness. All sites in this combined group are characterized by small sample sizes, none exceeding 20 samples. The particularly limited numbers at FF (n = 11) and P (n = 7) make sampling bias a plausible explanation for the observed differences. Therefore, isotopic values of herbivores appear largely consistent across sites within each combined group for the Later Medieval period, indicating no significant intra-group variation.

**Fig. 34.**
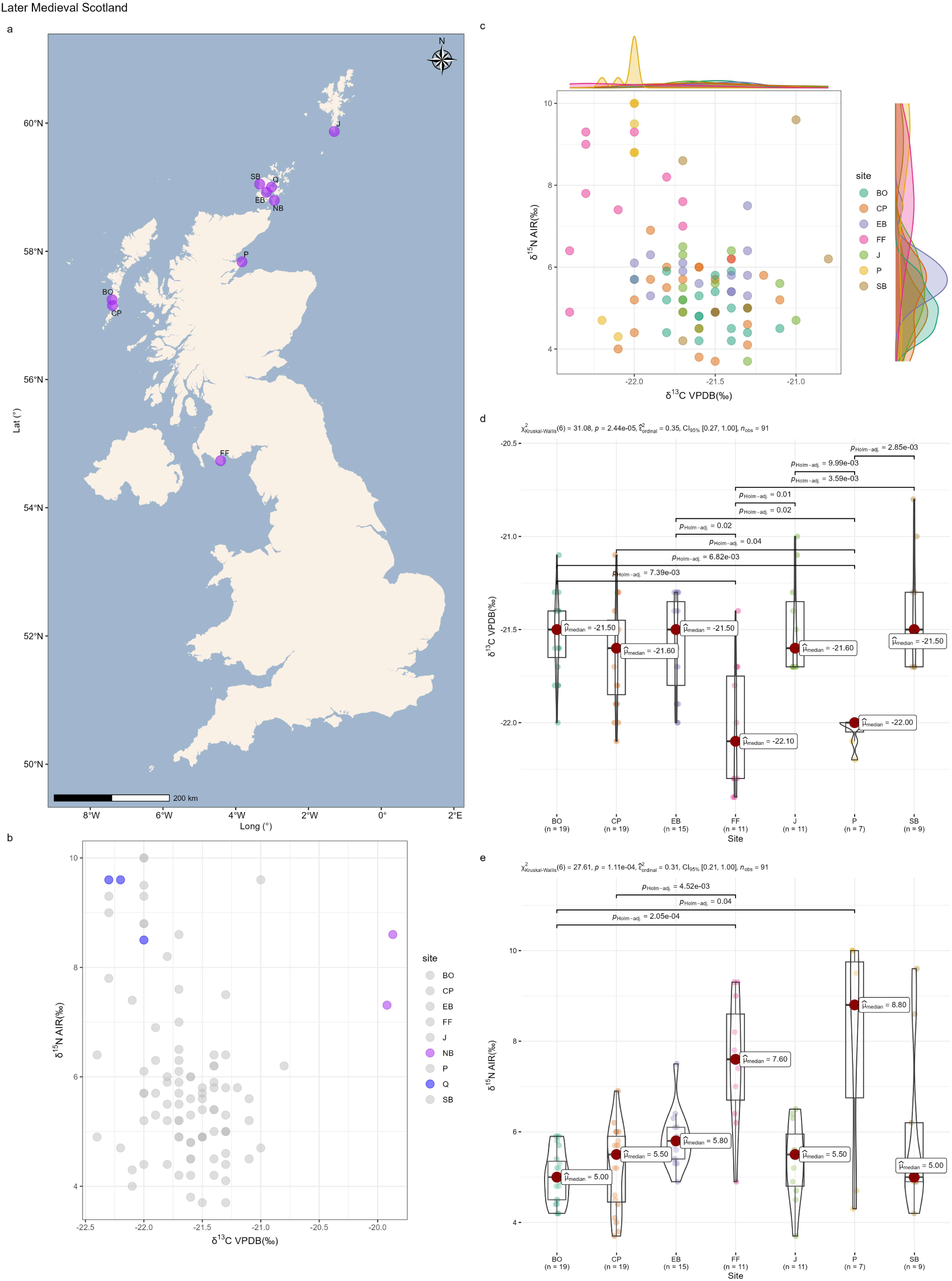
Intra-group comparison of terrestrial herbivores from Later Medieval Scotland. **a,** Distribution of sites. **b,** Scatter plot (sparsely sampled sites). **c,** Scatter plot (well-sampled sites). **d,** Statistical tests for δ^13^C (well-sampled sites). **e,** Statistical tests for δ^15^N (well-sampled sites). Site names: BO: Bornais; CP: Cille Phedair; EB: Earl’s Bu; FF: Fey Field; J: Jarlshof; P: Portmahomack; SB: Skara Brae; NB: Site of Newark Bay; Q: Quanterness.

Inter-group comparisons are shown in Fig. 35. The regions with the large sample sizes are Scotland (n = 96) and northern England (n = 65). While their isotopic values partly overlap, they remain broadly distinct. Meanwhile, samples from Wales (n = 19) and southern England (n = 40) lie within the isotopic range of northern England. The Welsh samples fall within a specific portion of the cluster formed by northern England. However, this cannot be taken as evidence that Wales differs significantly, as it is represented by only 19 samples, far fewer than that of northern England.

**Fig. 35.**
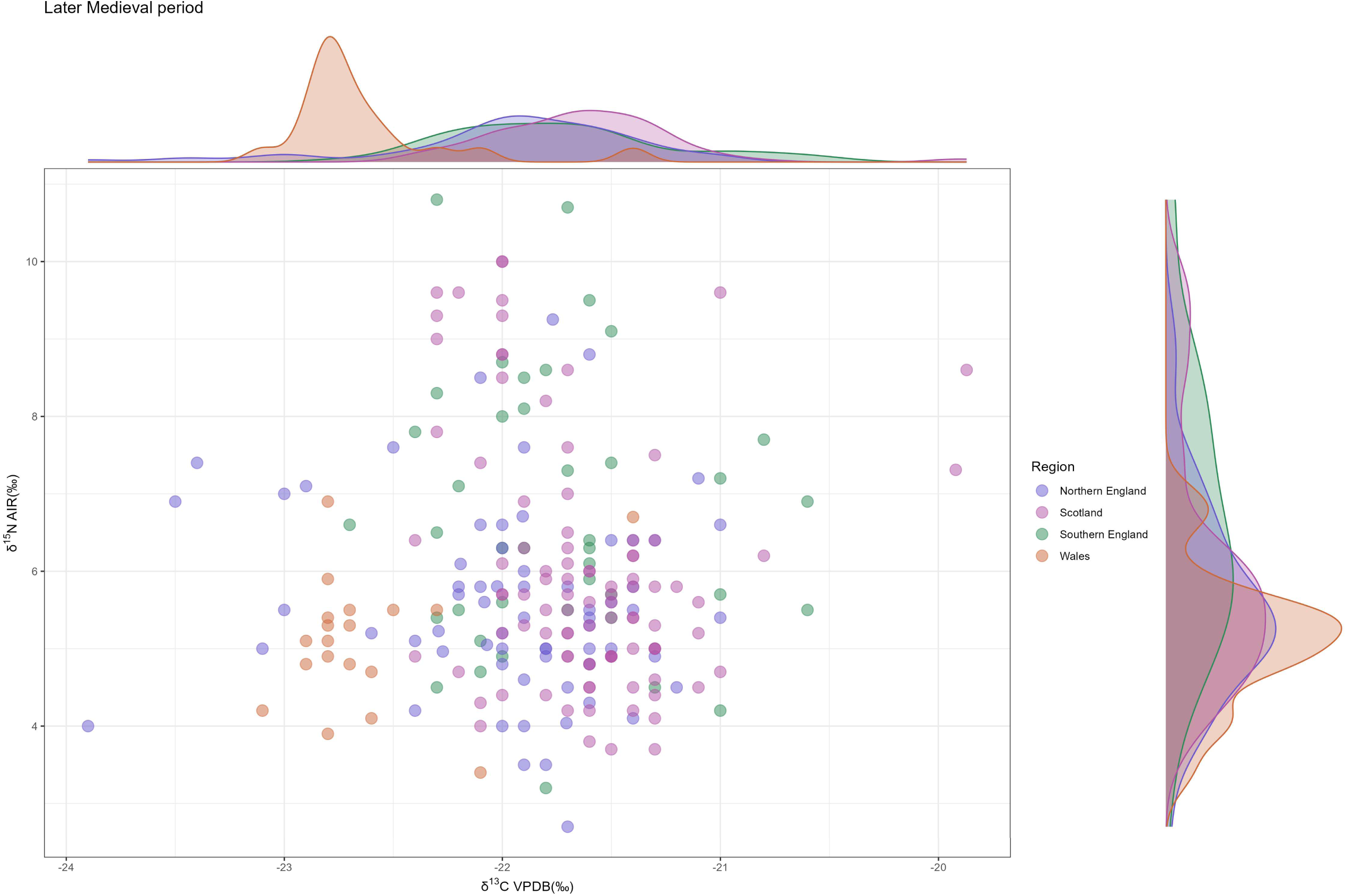
Inter-group comparison of terrestrial herbivores from the Later Medieval period (Scatter plot).

For the Post-Medieval period, the dataset is extremely limited, comprising only two sites in southern England. Both sites display similar isotopic patterns, as shown by the scatter plot and statistical tests (Fig. 36).

**Fig. 36.**
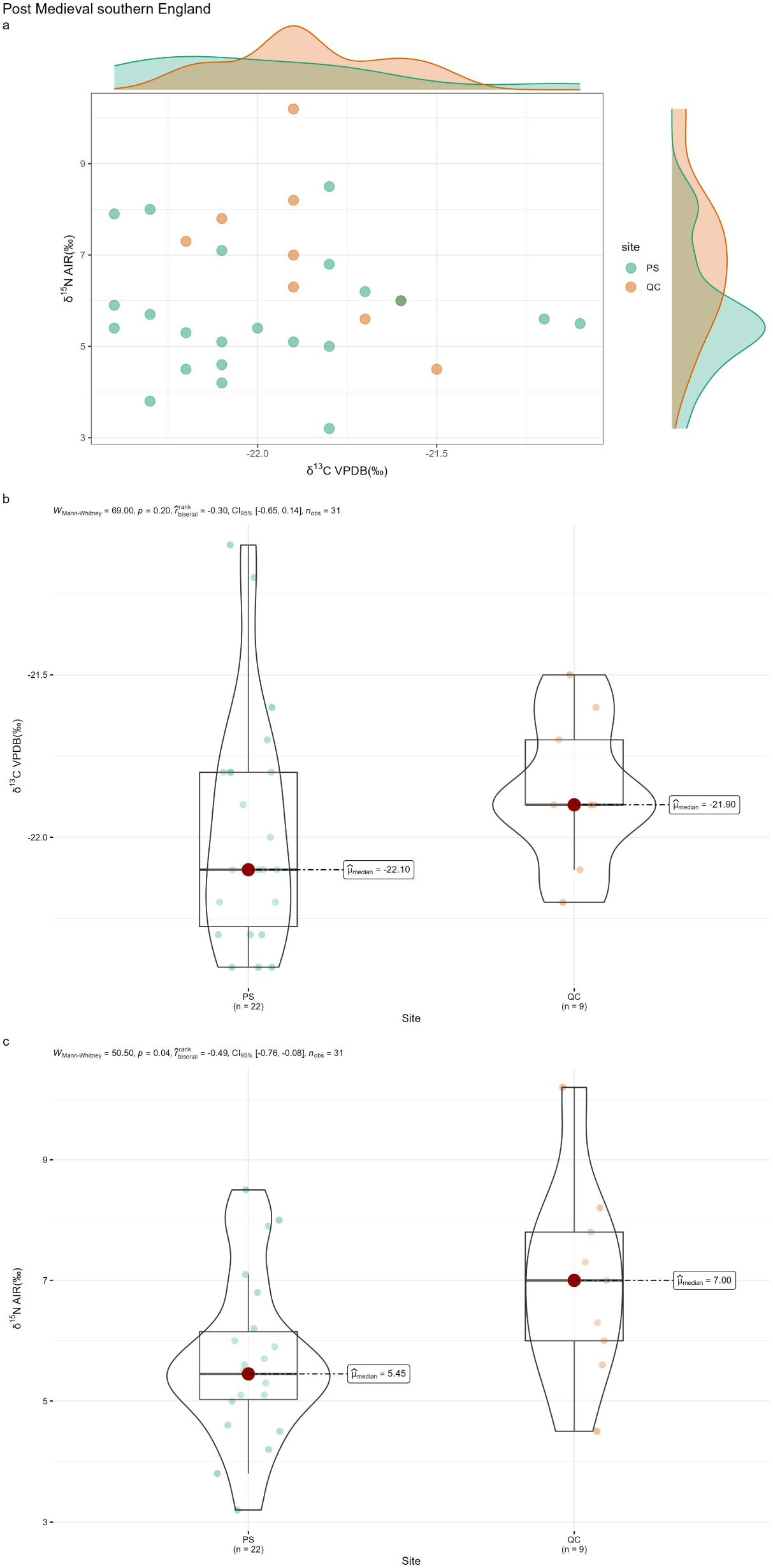
Intra-group comparison of terrestrial herbivores from Post Medieval southern England. **a,** Scatter plot. **b,** Statistical tests for δ^13^C. **c,** Statistical tests for δ^15^N. Site names: PS: Prescot Street; QC: Queen’s Chapel of the Savoy.

### Analysis of isotopic variation for terrestrial omnivores

#### Paleolithic Period to Bronze Age

We collected only minimal data on omnivores in the Paleolithic and Mesolithic periods. As a result, intra-group comparisons could not be conducted for these periods. The available data are shown in Fig. 37, but no conclusions can be drawn due to the limited sample size.

**Fig. 37.**
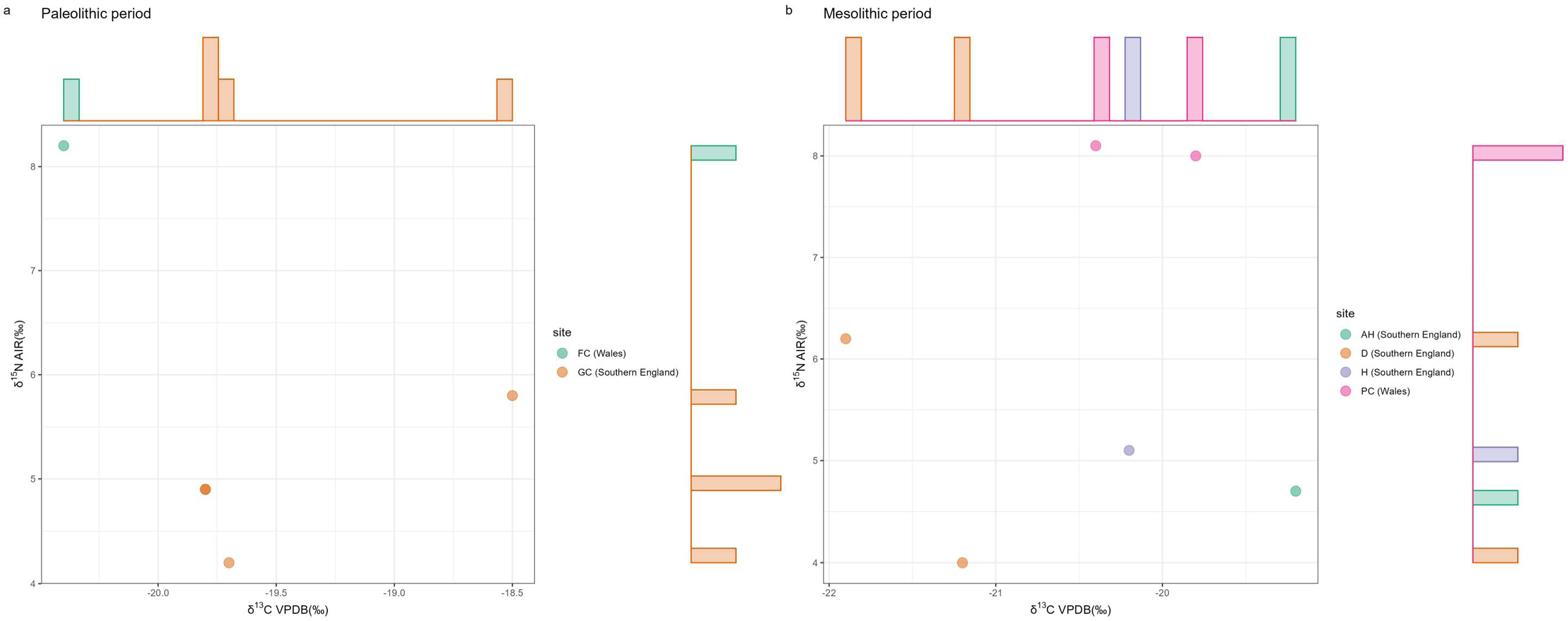
Intra-group comparison of terrestrial omnivores from the Paleolithic and Mesolithic period (Scatter plots). **a,** Scatter plot (Paleolithic period). **b,** Scatter plot (Mesolithic period). Site names: Paleolithic: FC: Foxhole Cave; GC: Gough’s Cave; Mesolithic: AH: Aveline’s Hole; D: Doggerland; H: Hazleton; PC: Potter’s Cave.

For the Neolithic period, intra-group comparisons can be made for southern England and Scotland. In southern England, the well-sampled sites AUW and HN appear to have distinct isotopic distributions (Fig. 38a), particularly in δ^15^N (Fig. 38e). AUW (n = 16) and HN (n = 13) have limited sample sizes, with δ¹⁵N values concentrated at 5–7.5‰ and 3.5–5.5‰, respectively. Based on the herbivore results, sites with larger sample sizes (>50) tend to show a more even spread across 2–8‰. Thus, the apparent difference here is more plausibly explained by sampling bias than by substantive isotopic variation. The remaining site with small sample sizes, EC, aligns with the cluster defined by AUW and HN. In Scotland, isotopic values from all six sites are intermixed (Fig. 38b). Statistical tests for the two sites with comparatively larger sample sizes yielded no significant differences (Fig. 38d and f).

**Fig. 38.**
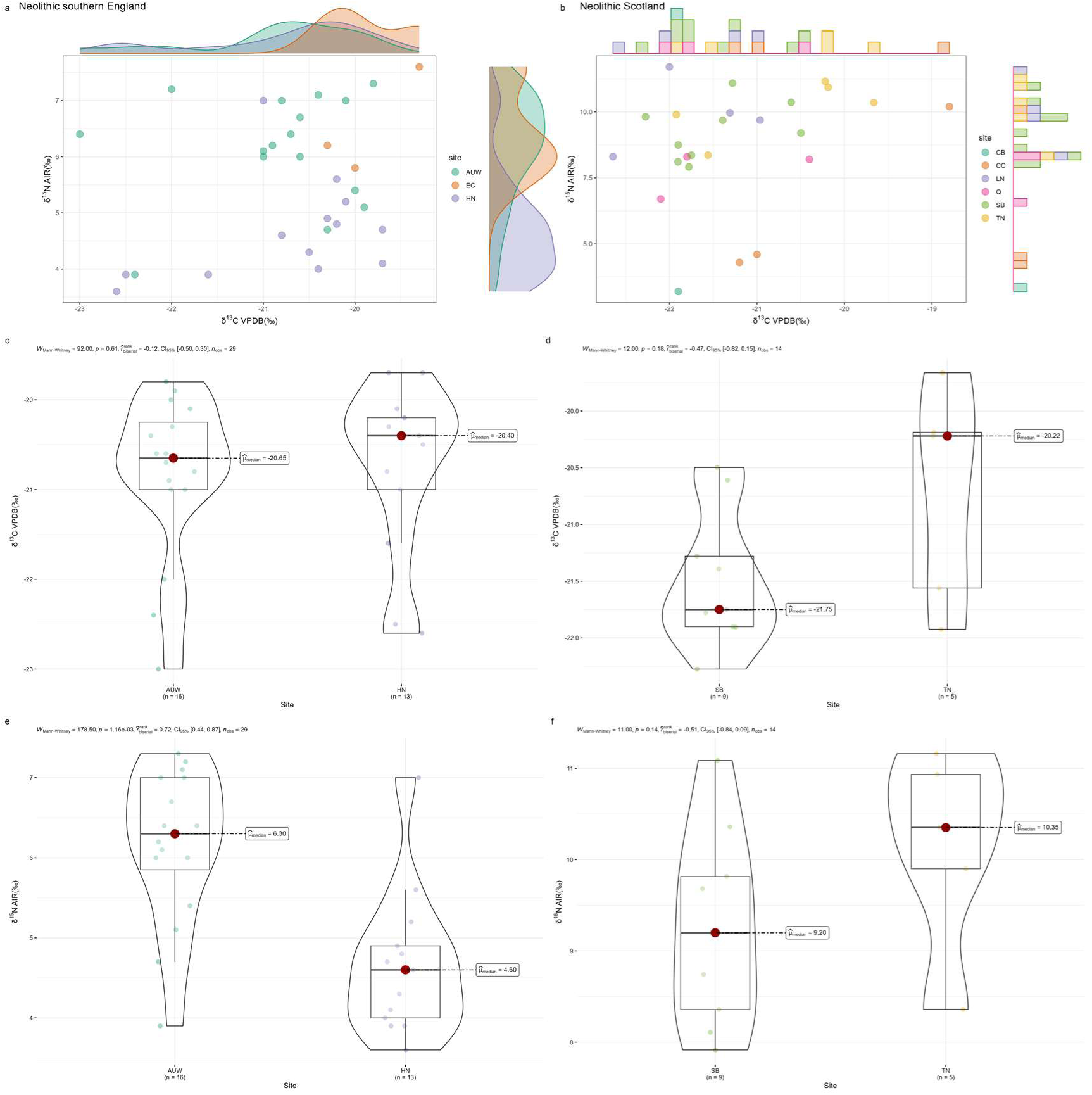
Intra-group comparison of terrestrial omnivores from the Neolithic period. **a,** Scatter plot (Neolithic southern England). **b,** Scatter plot (Neolithic Scotland). **c,** Statistical tests for δ^13^C (Neolithic southern England). **d,** Statistical tests for δ^13^C (Neolithic Scotland). **e,** Statistical tests for δ^15^N (Neolithic southern England). **f,** Statistical tests for δ^15^N (Neolithic Scotland). Site names: Neolithic southern England; AUW: Ascott-Under-Wychwood; EC: Eton College Rowing Course; HN: Hazleton North; Neolithic Scotland: CB: Carding Mill Bay; CC: Cnoc Coig; LN: Links of Noltland; Q: Quanterness; SB: Skara Brae; TN: Tofts Ness.

In the Bronze Age, intra-group comparisons can be made for the Bronze Age southern England group. Both the scatter plot and statistical tests indicate that the sites share similar isotopic distributions (Fig. 39a, c, d). In Scotland, five Bronze Age sites are represented, but four contain only a single sample (Fig. 39b), preventing meaningful conclusions. Building on the above analysis, we conclude that omnivores within the combined groups of these periods do not exhibit significant isotopic variation among sites.

**Fig. 39.**
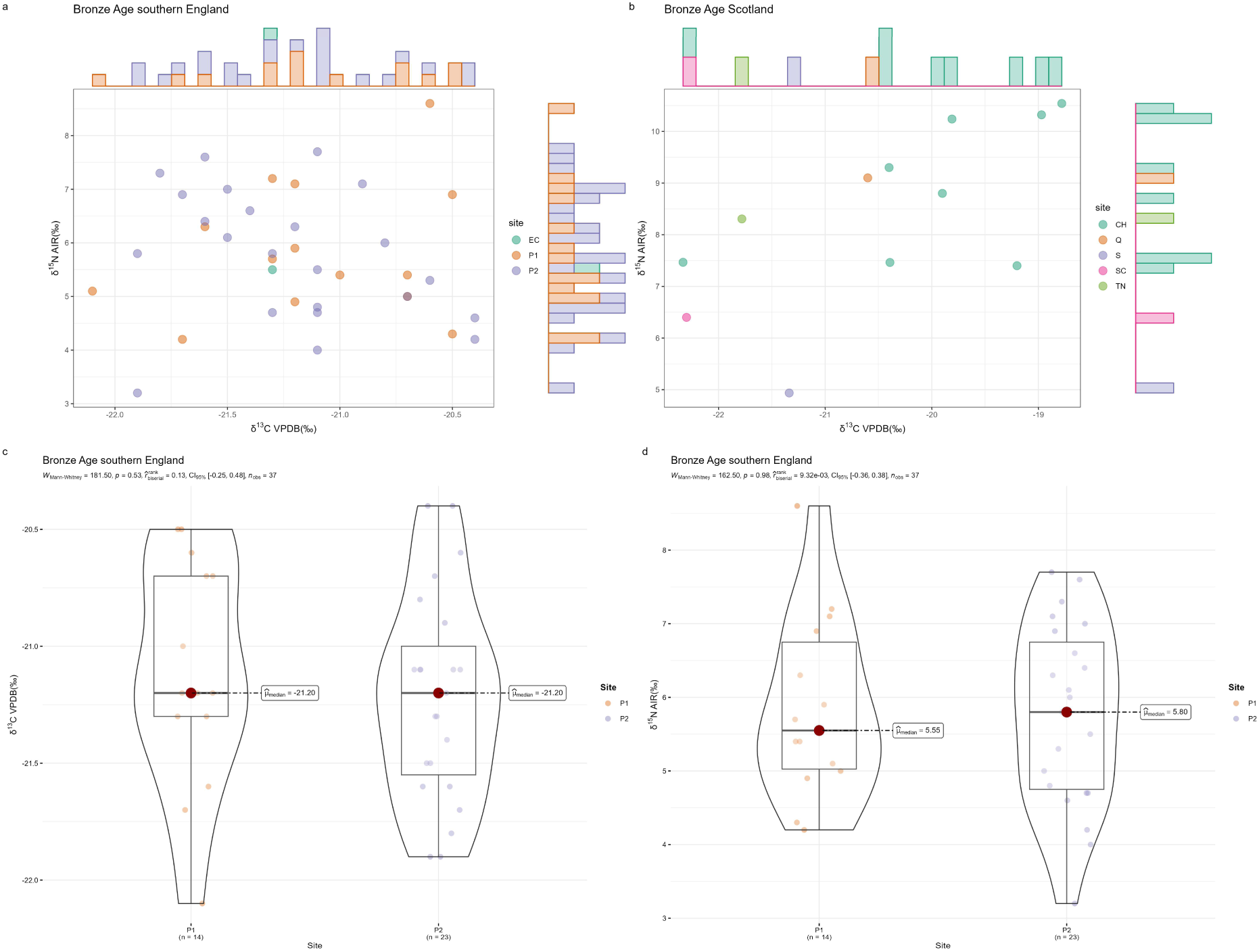
Intra-group comparison of terrestrial omnivores from the Bronze Age. **a,** Scatter plot (Bronze Age southern England). **b,** Scatter plot (Bronze Age Scotland). **c,** Statistical tests for δ^13^C (Bronze Age southern England). **d,** Statistical tests for δ^15^N (Bronze Age southern England). Site names: Bronze Age southern England: EC: Eton College Rowing Course; P1: Potterne 1; P2: Potterne 2; Bronze Age Scotland: CH: Cladh Hallan; Q: Quanterness; S: Sligenach; SC: Sculptor’s Cave; TN: Tofts Ness.

Inter-group comparisons were conducted for the Neolithic and Bronze Age (Fig. 40). Unlike the inter-group results for herbivores in most periods, Neolithic omnivores from Scotland and southern England form two clearly distinct clusters. This may reflect the effect of small sample sizes, as Neolithic omnivores are represented by far fewer samples than herbivores in any period. In the Bronze Age, where sample sizes are somewhat larger, inter-group comparisons yield patterns more consistent with herbivores: samples from southern England and Wales (represented by a single site) show nearly identical isotopic distributions, while some Scottish samples fall outside the main cluster defined by southern England and Wales.

**Fig. 40.**
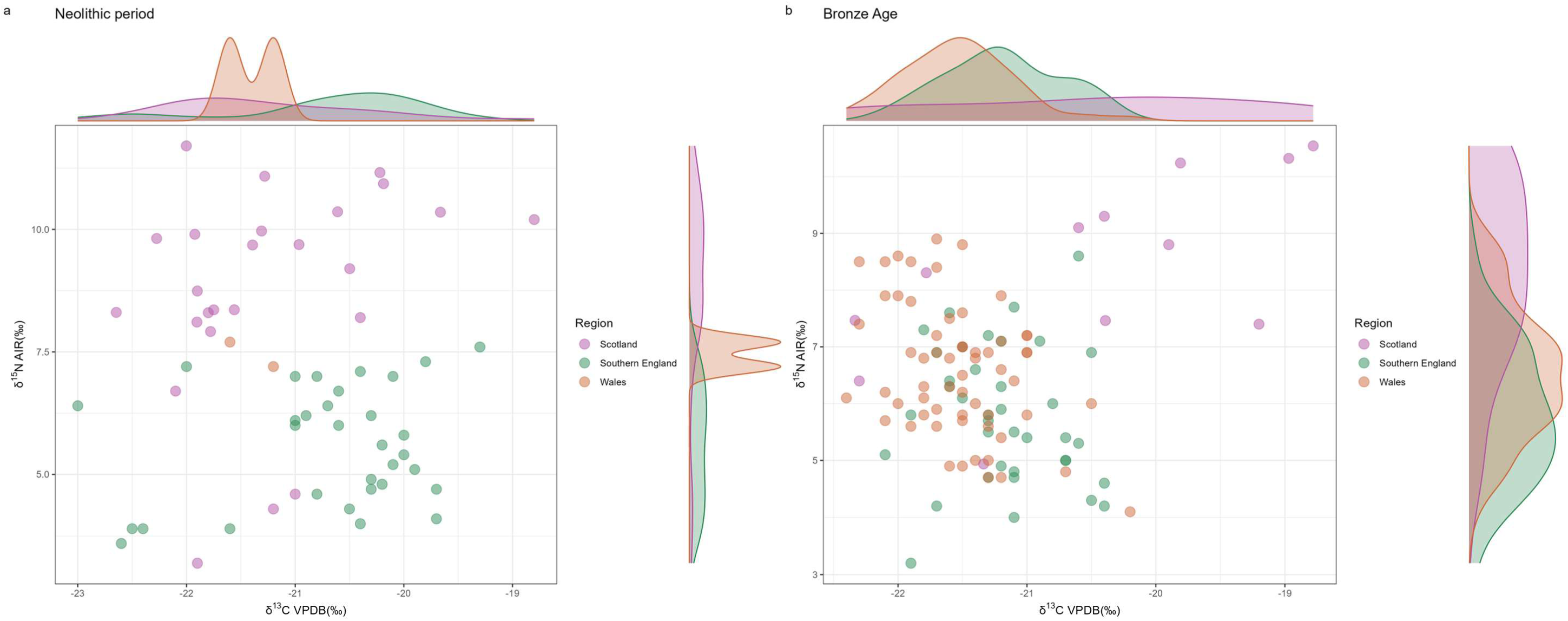
Inter-group comparison of terrestrial omnivores from two periods (Scatter plots). **a,** Neolithic period. **b,** Bronze Age.

### Iron Age

We conducted intra-group comparisons for Iron Age southern England and Scotland. For southern England, we first calculated mean isotopic values and standard deviations for each site (Fig. 41a). As with herbivores, isotopic values from the same site but different phases show highly consistent patterns (e.g., DH1–DH3, DRH1–DRH2, NC1–NC2, SF1–SF3, Y1–Y3). These phases were therefore combined into single sites for subsequent analyses (Fig. 41b and Fig. 42c). The combined datasets were then subjected to the statistical tests (Fig. 42e and f), which produced significant results for both δ^13^C and δ^15^N. Post hoc tests indicate that the main sites driving these differences are GLV (δ^13^C) and Y (δ^15^N). However, GLV includes only five samples, so the statistical difference is unlikely to have practical meaning. Although Y shows significant differences from some other sites in δ^15^N values, all of its sample values fall within the cluster defined by other sites (Fig. 42f). These results suggest that limited sample sizes may underlie the apparent significance. Sites with too few samples for the statistical tests are shown separately in Fig. 42d. All of their isotopic values fall within the main cluster. Site locations are plotted in Fig. 42a and b, which shows that Y is not geographically distinct from other sites. This further supports the interpretation that the anomaly at Y reflects sample size limitations rather than genuine geographic patterning.

**Fig. 41.**
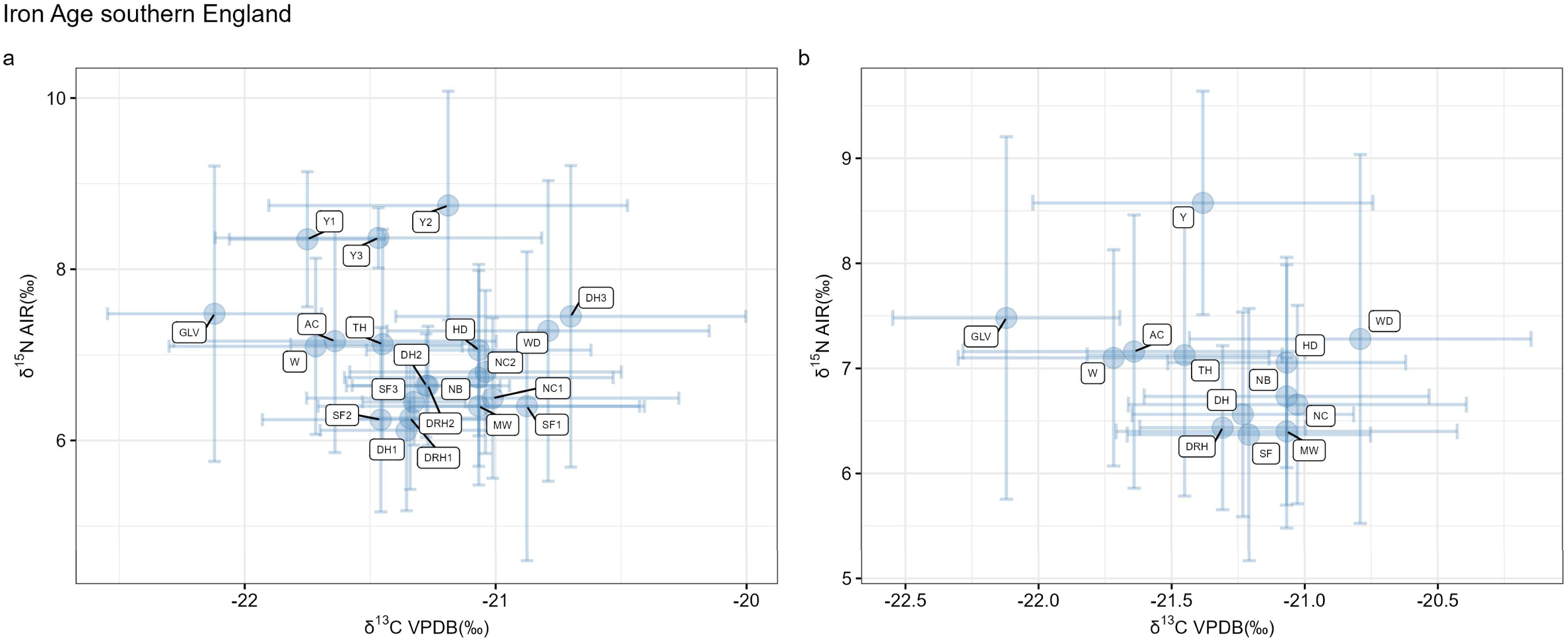
Intra-group comparison of terrestrial omnivores from Iron Age southern England (Average isotopic values with standard deviations). **a,** Average isotopic values with standard deviations prior to combining the same sites across different periods (well-sampled sites). **b,** Average isotopic values with standard deviations after combining the same sites across different periods (well-sampled sites). Site names: AC: Alfred’s Castle; DH: Danebury Hillfort (DH1: Danebury Hillfort 1; DH2: Danebury Hillfort 2; DH3: Danebury Hillfort 3); DRH: Danebury Ring Hillfort (DRH1: Danebury Ring Hillfort 1; DRH2: Danebury Ring Hillfort 2); GLV: Glastonbury Lake Village; HD: Houghton Down; MW: Micheldever Wood; NB: New Buildings; NC: Nettlebank Copse (NC1: Nettlebank Copse 1; NC2: Nettlebank Copse 2); SF: Suddern Farm (SF1: Suddern Farm 1; SF2: Suddern Farm 2; SF3: Suddern Farm 3); TH: Trethellan Head; W: Watchfield; WD: Winnall Down; Y: Yarnton (Y1: Yarnton 1; Y2: Yarnton 2; Y3: Yarnton 3).

**Fig. 42.**
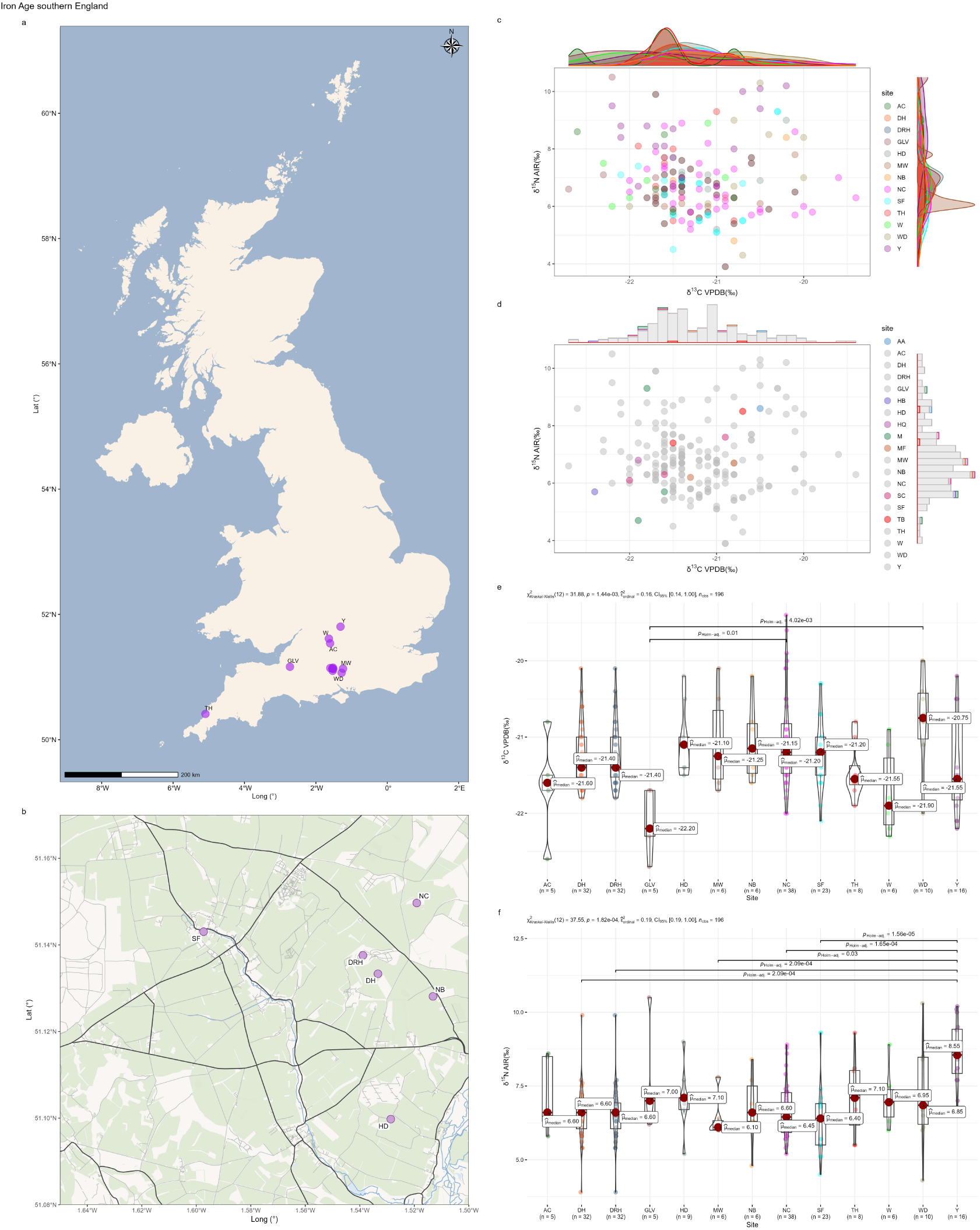
Intra-group comparison of terrestrial omnivores from Iron Age southern England. **a,** Distribution of sites. **b,** Distribution of six proximate sites. **c,** Scatter plot (well-sampled sites). **d,** Scatter plot (sparsely sampled sites). **e,** Statistical tests for δ^13^C (well-sampled sites). **f,** Statistical tests for δ^15^N (well-sampled sites). Site names: AC: Alfred’s Castle; DH: Danebury Hillfort; DRH: Danebury Ring Hillfort; GLV: Glastonbury Lake Village; HD: Houghton Down; MW: Micheldever Wood; NB: New Buildings; NC: Nettlebank Copse; SF: Suddern Farm; TH: Trethellan Head; W: Watchfield; WD: Winnall Down; Y: Yarnton; AA: Alington Avenue; HB: Harlyn Bay; HQ: Horcott Quarry; M: Marcham; MF: Manor Farm; SC: Segsbury Camp; TB: Tolpuddle Ball.

The situation for Scotland is more straightforward. The only site that differs from the others is DV, and even this difference is slight (p = 0.03) (Fig. 43e). Given the small sample sizes across the group (ranging from 6 to 11 samples per site), such statistical results are likely attributable to the small sample size. This interpretation is reinforced by the site locations, as DV is not geographically distinct from the others, especially BA and BOM1 (Fig. 43a). Isotopic values from the remaining sparsely sampled sites fall within the range of the main cluster (Fig. 43b).

**Fig. 43.**
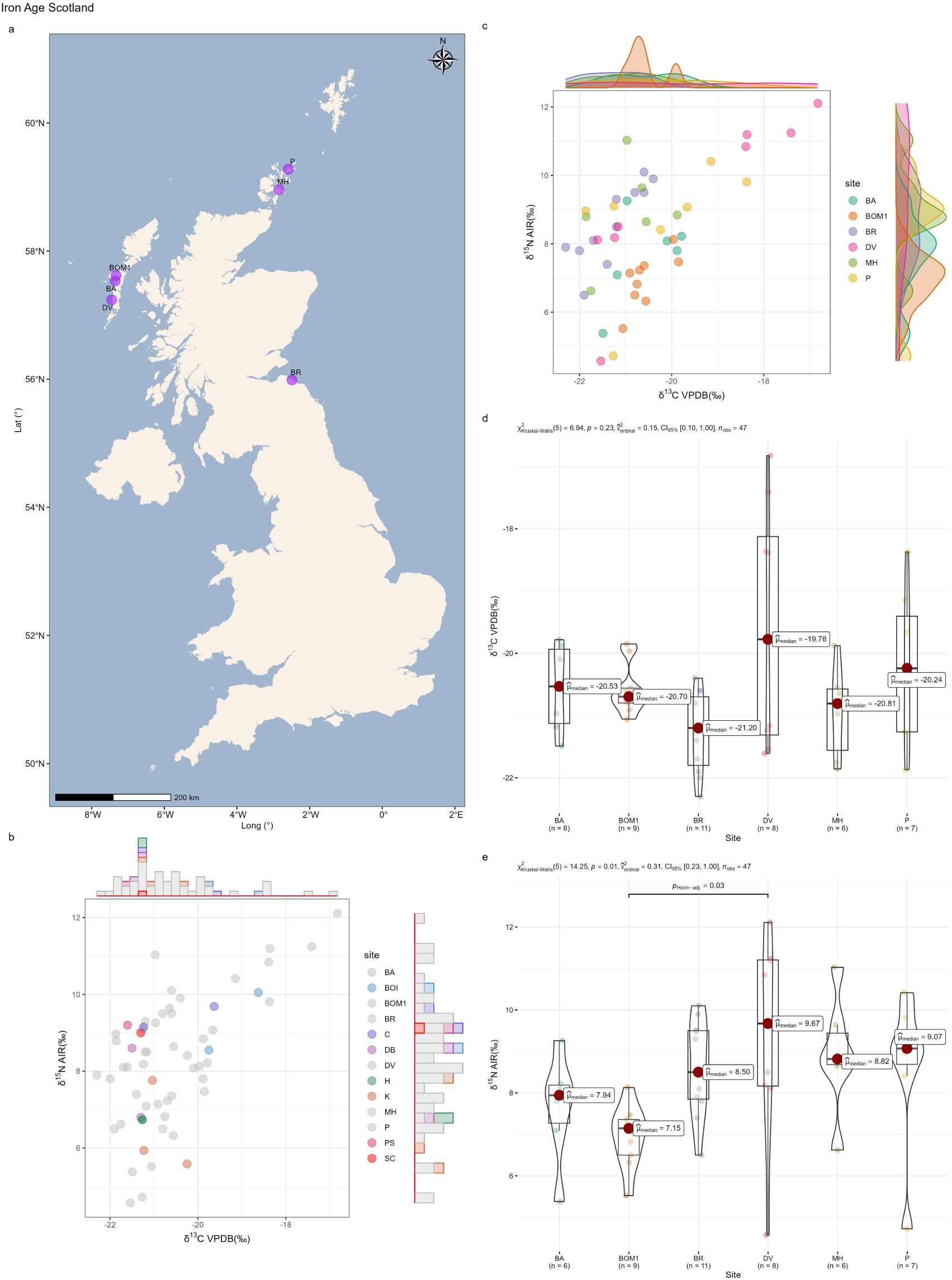
Intra-group comparison of terrestrial omnivores from Iron Age Scotland. **a,** Distribution of sites. **b,** Scatter plot (sparsely sampled sites). **c,** Scatter plot (well-sampled sites). **d,** Statistical tests for δ^13^C (well-sampled sites). **e,** Statistical tests for δ^15^N (well-sampled sites). Site names: BA: Baleshare; BOM1: BOM1 (Western Isles); BR: Broxmouth; DV: Dun Vulan; MH: Mine Howe; P: Pool; BOI: Bornish M2W; C: The Cairns; DB: Dryburn Bridge; H: Howe; K: Knowe o’ Skea; PS: Port Seton; SC: Sculptor’s Cave.

Inter-group comparisons yield results similar to those observed for herbivores: southern and northern England cluster together, while most Scottish samples also fall within this cluster, with a few lying outside (Fig. 44).

**Fig. 44.**
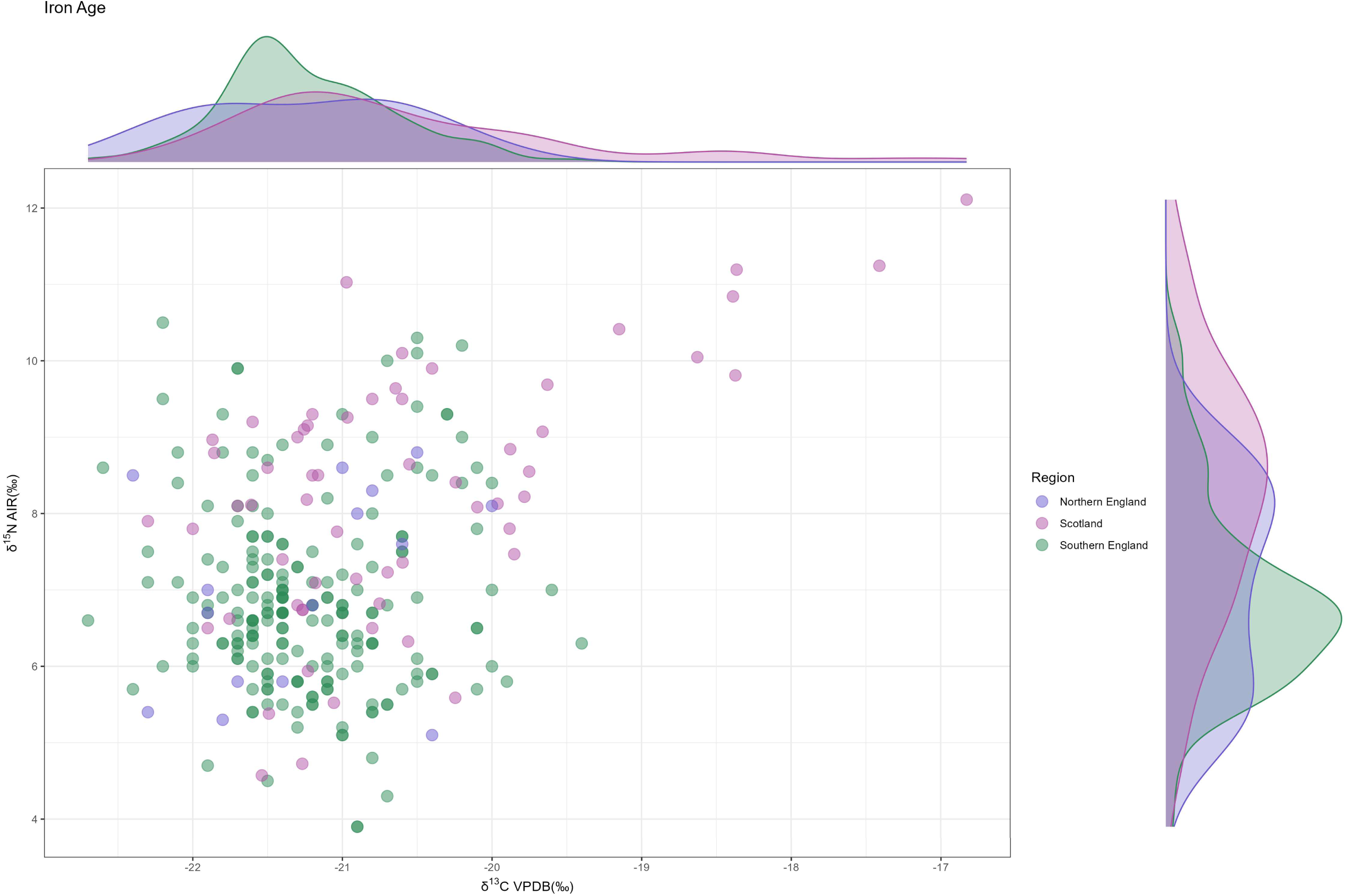
Inter-group comparison of terrestrial omnivores from the Iron Age (Scatter plot).

### Roman period/Roman Iron Age and Early Medieval Period

For the Roman period, intra-group comparisons were conducted for southern and northern England. In southern England, all sites show broadly similar isotopic values, with the exception of SG in δ^15^N (Fig. 45g). Sparsely sampled sites are plotted in Fig. 45e, and all of their values fall within the cluster defined by the well-sampled sites. Examination of site locations (Fig. 45c) shows that SG is geographically very close to L (almost the same site). Geographically, this pattern reflects variation within a single site. Thus, the observed statistical significance likely results from sampling bias. Thus, isotopic values among sites in Roman southern England can be regarded as similar. The sample sizes of L and SG are 12 and 13, respectively. This further illustrates that when the number of samples per site falls below 20, as is the case for most sites, spurious statistical significance can easily arise due to sampling bias. If L and SG are regarded as a single site, their δ^15^N values would span evenly across a range of 4–10‰. Similar to the pattern seen in herbivores, this indicates that when the sample size of omnivores at a given site is sufficiently large, their isotopic values will cover a wide and uniform range. Consequently, isotopic differences among sites are expected to be minimal.

**Fig. 45.**
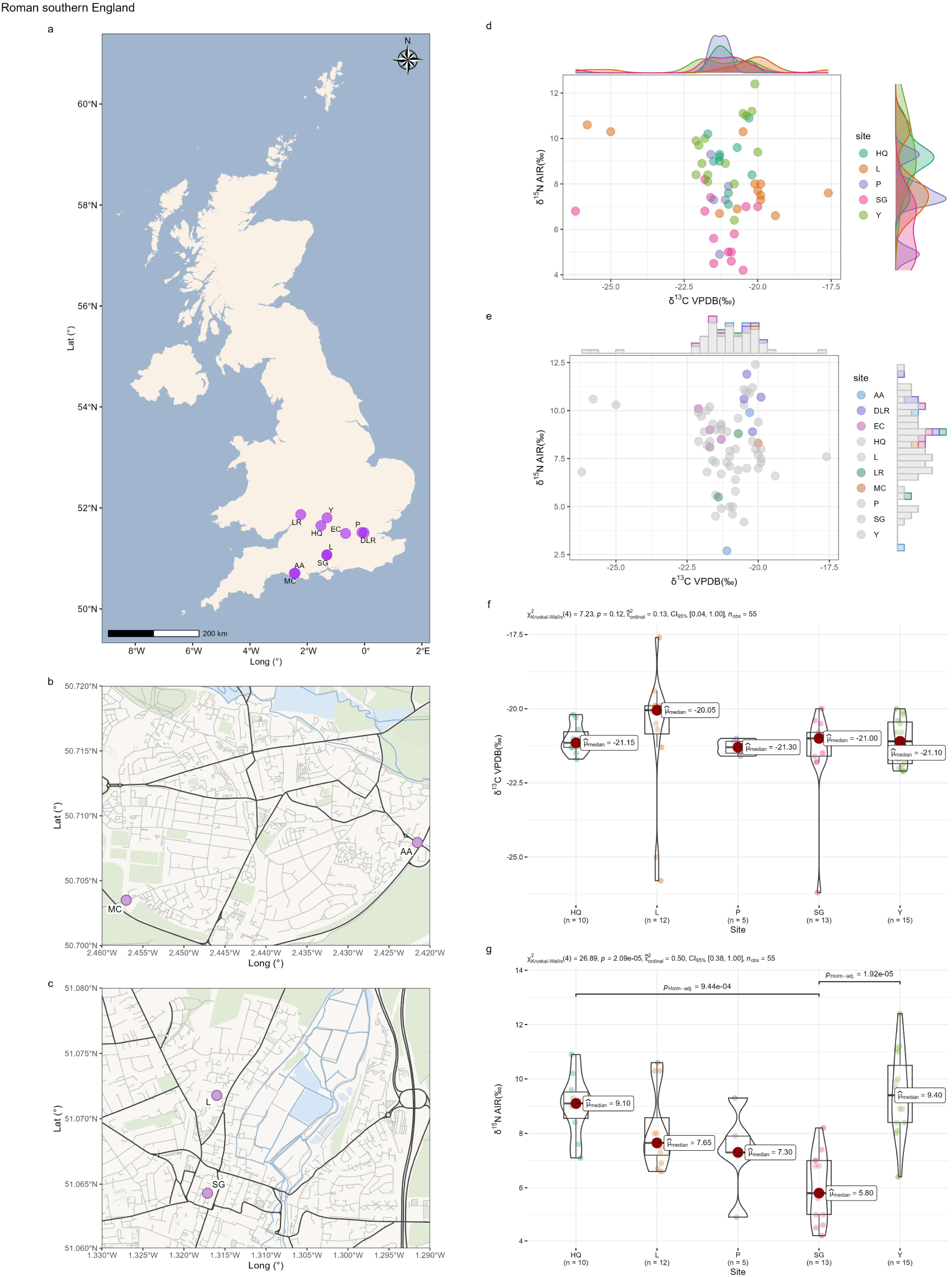
Intra-group comparison of terrestrial omnivores from Roman southern England. **a,** Distribution of sites. **b,** Distribution of two proximate sites (MC and AA). **c,** Distribution of two proximate sites (L and SG). **d,** Scatter plot (well-sampled sites). **e,** Scatter plot (sparsely sampled sites). **f,** Statistical tests for δ^13^C (well-sampled sites). **g,** Statistical tests for δ^15^N (well-sampled sites). Site names: HQ: Horcott Quarry; L: Lankhills; P: 1 Poultry; SG: Staple Gardens; Y: Yarnton; AA: Alington Avenue; DLR: Docklands Light Railway; EC: Eton College Rowing Course; LR: 120-122 London Road; MC: Maiden Castle Road.

Roman northern England includes only three sites. The isotopic distributions of sites with large and small sample sizes are shown in Fig. 46. Sites with large sample sizes display similar isotopic distributions (Fig. 46a, c, d). The isotopic values of the sparsely sampled site fall within the isotopic range defined by the well-sampled ones (Fig. 46b).

**Fig. 46.**
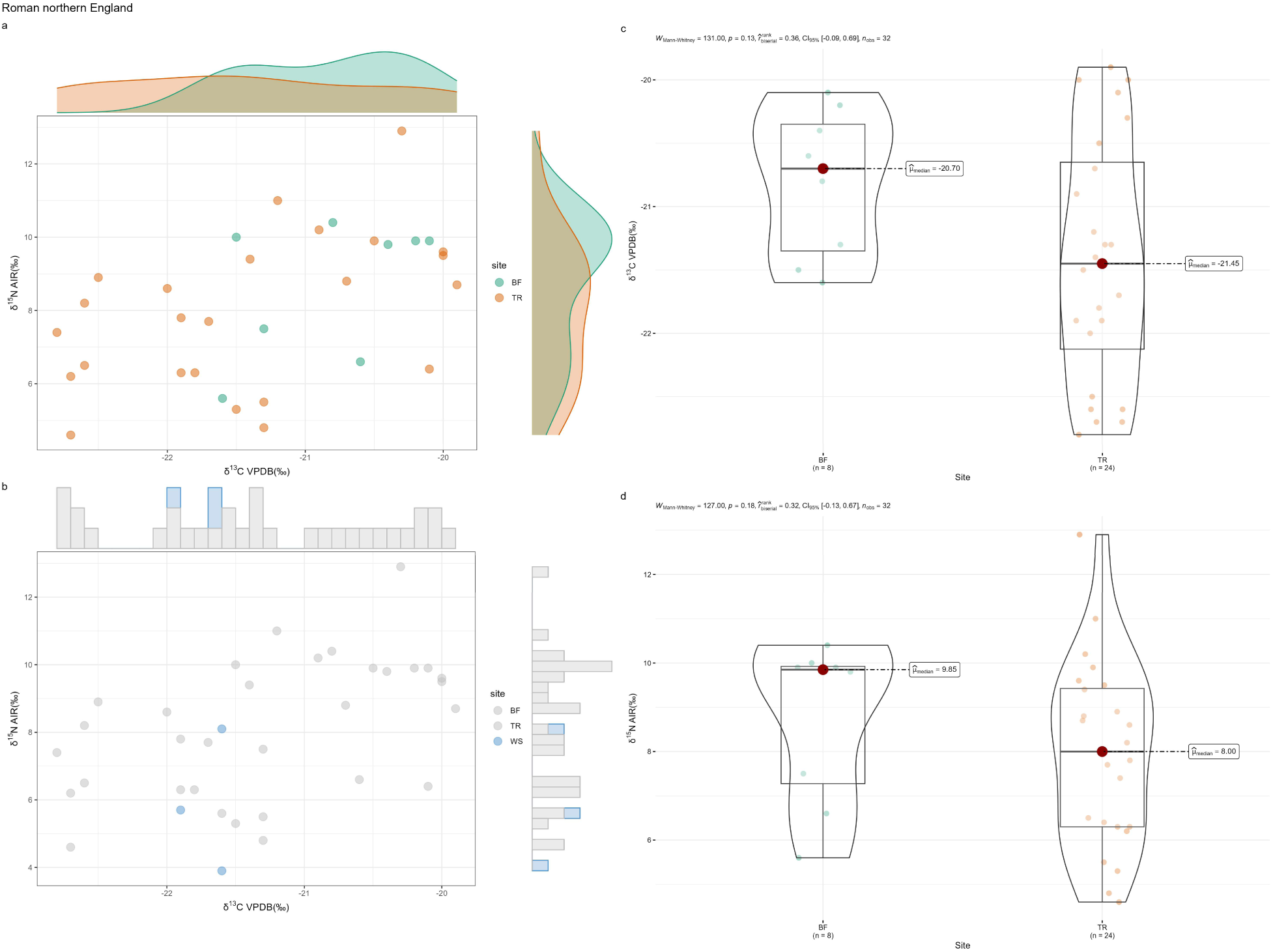
Intra-group comparison of terrestrial omnivores from the Roman northern England. **a,** Scatter plot (well-sampled sites). **b,** Scatter plot (sparsely sampled sites). **c,** Statistical tests for δ^13^C (well-sampled sites). **d,** Statistical tests for δ^15^N (well-sampled sites). Site names: BF: Bainesse Farm; TR: Tanner Row; WS: Wetwang Slack.

Inter-group comparisons for the Roman period show that southern and northern England exhibit nearly identical isotopic distributions (Fig. 47). The few samples from Wales and Scotland also fall within the clusters defined by England.

**Fig. 47.**
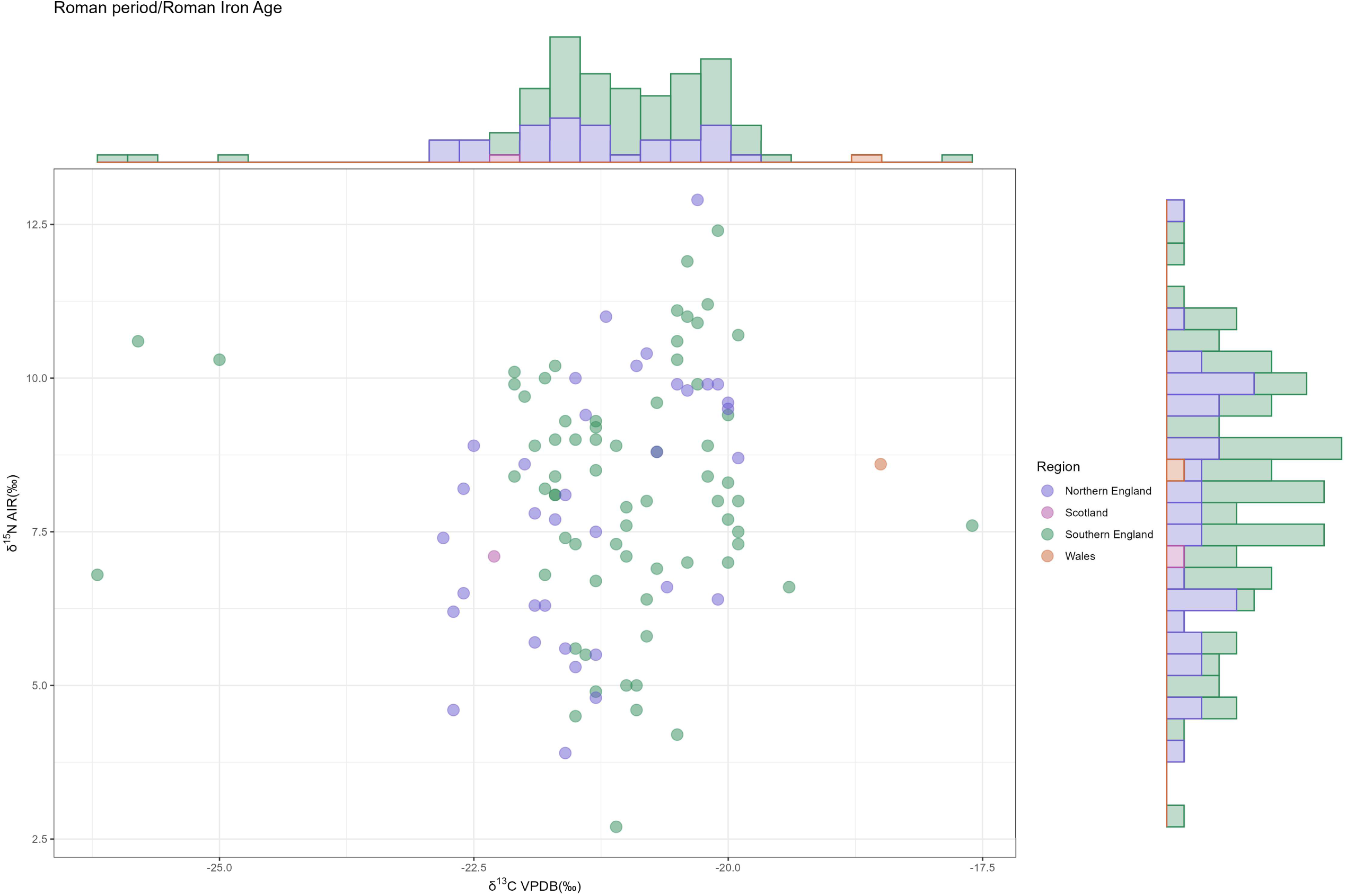
Inter-group comparison of terrestrial omnivores from the Roman period/Roman Iron Age (Scatter plot).

For the Early Medieval period, intra-group comparisons were conducted for northern England, Scotland, and southern England. In northern England, isotopic values show no significant differences across sites, as indicated by both the scatter plot and statistical tests (Fig. 48). For Scotland and southern England, statistical tests could not be performed because of the highly uneven sample sizes, so only scatter plots were produced. These results indicate that, within each region, isotopic values from sites with smaller sample sizes generally fall within the clusters defined by the larger sites (Fig. 49), indicating no significant isotopic differences among sites in the same region.

**Fig. 48.**
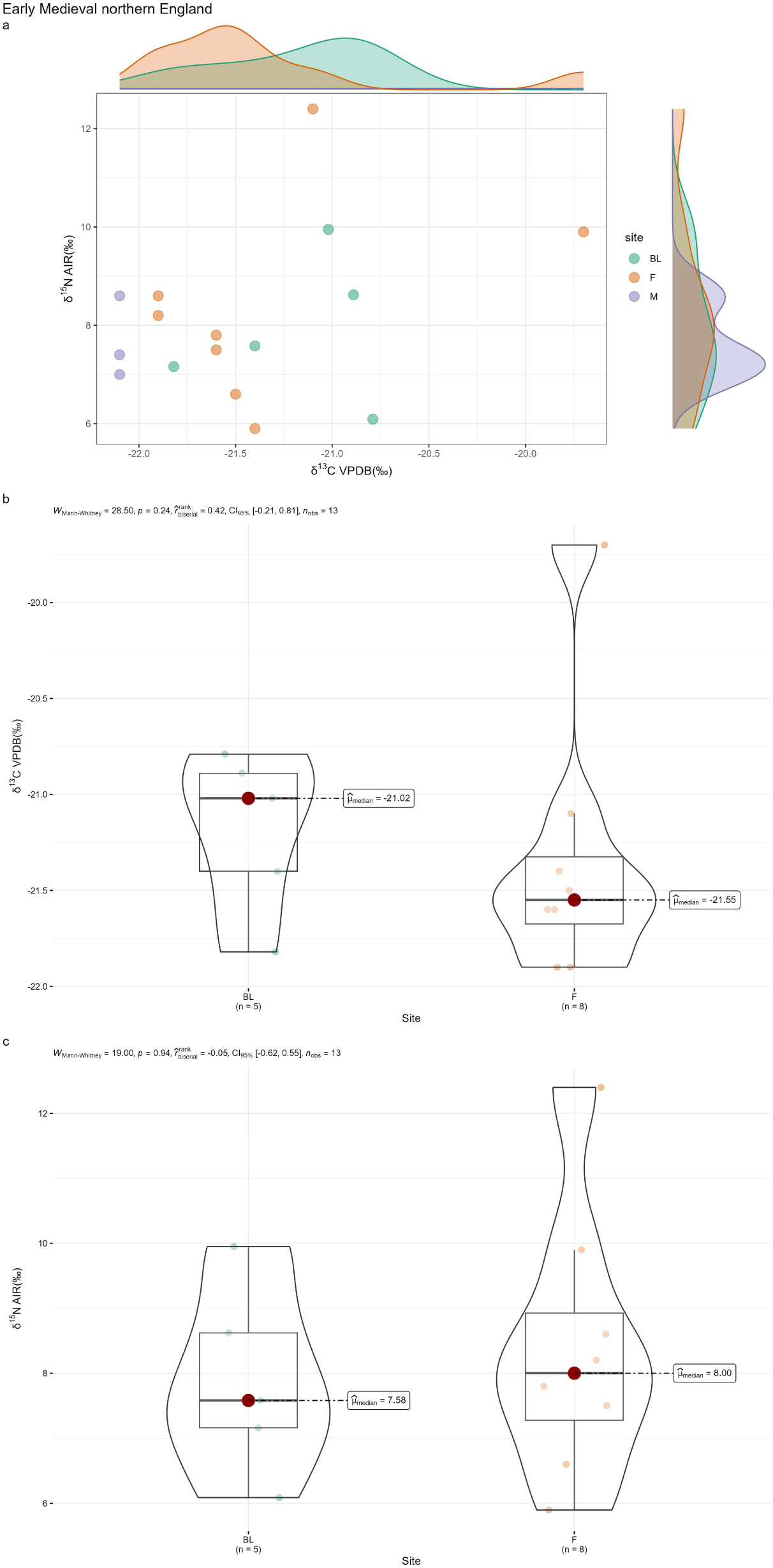
Intra-group comparison of terrestrial omnivores from Early Medieval northern England. **a,** Scatter plot. **b,** Statistical tests for δ^13^C. **c,** Statistical tests for δ^15^N. Site names: BL: Blackgate; F: Fishergate; M: Masham.

**Fig. 49.**
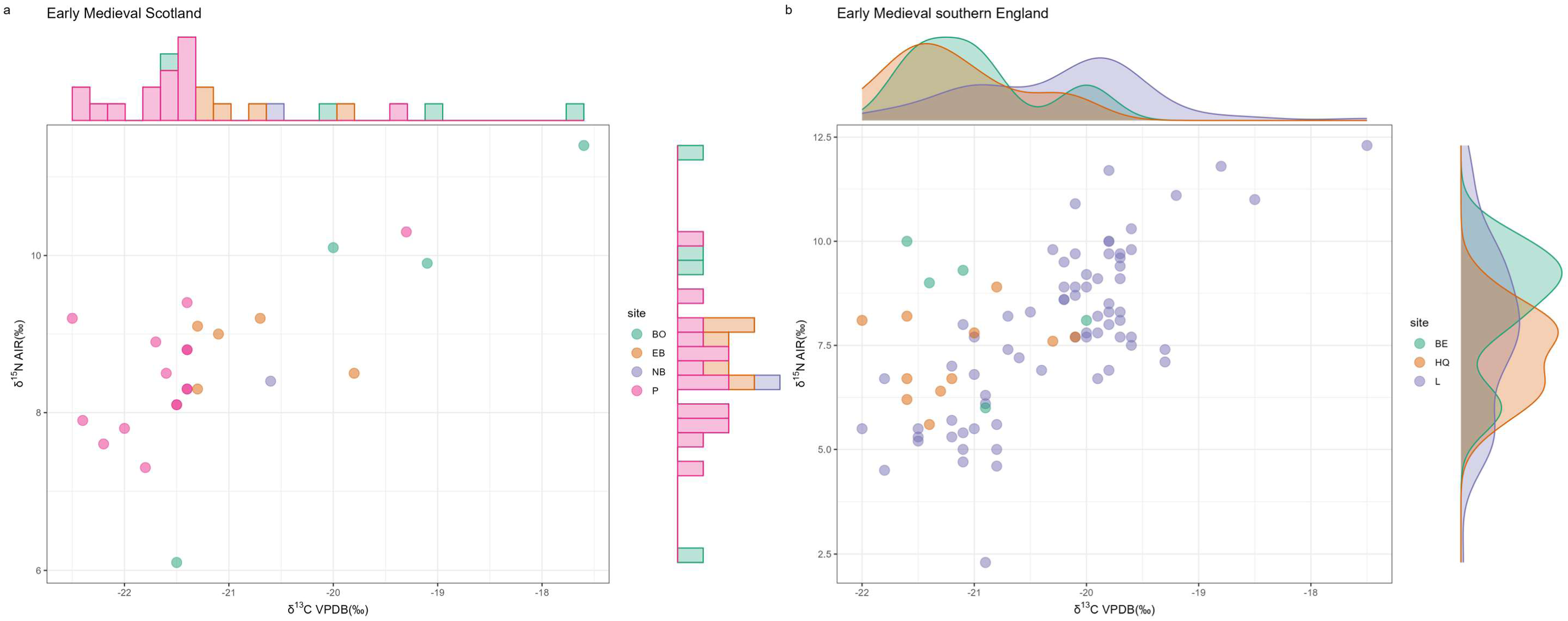
Intra-group comparison of terrestrial omnivores from Early Medieval Scotland and southern England (Scatter plots). **a,** Early Medieval Scotland. **b,** Early Medieval southern England. Site names: Scotland: BO: Bornais; EB: Earl’s Bu; NB: Site of Newark Bay; P: Portmahomack; Southern England: BE: Beaconsfield; HQ: Horcott Quarry; L: Lyminge.

Inter-group comparisons further demonstrate that isotopic values from nearly all samples in Scotland, Wales, central England, and northern England (all with smaller sample sizes than southern England) fall within the cluster defined by southern England (Fig. 50). This suggests that isotopic values do not differ substantially among combined groups in the Early Medieval period.

**Fig. 50.**
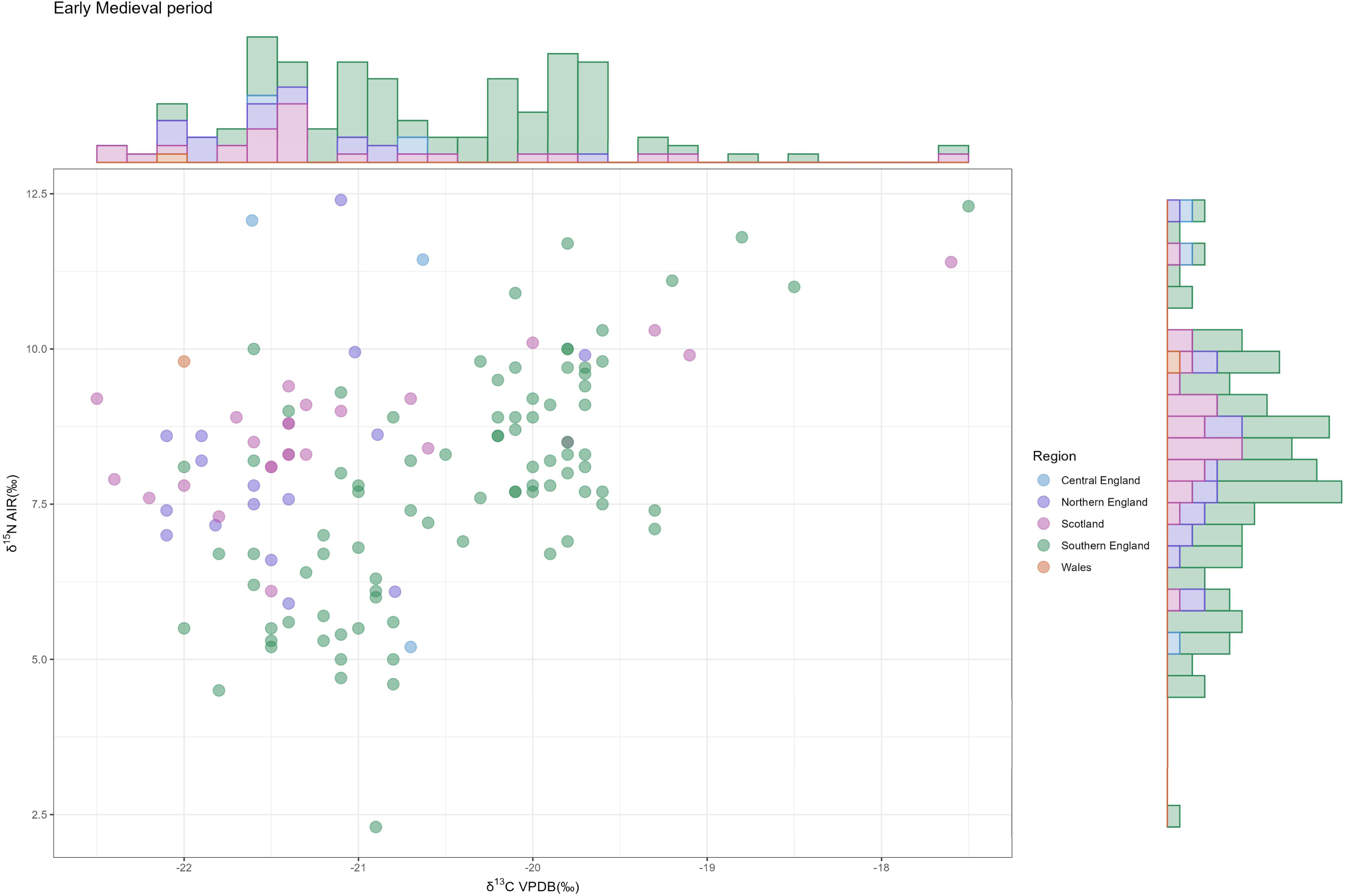
Inter-group comparison of terrestrial omnivores from the Early Medieval period (Scatter plot).

### Later Medieval and Post Medieval Periods

We conducted three intra-group comparisons for the Later Medieval period: southern England, northern England, and Scotland. In southern England, one post hoc comparison (AS vs. SA) of δ^15^N shows significant results (Fig. 51d). However, given the sample sizes and the close geographic proximity of these two sites (Fig. 51a), this statistical result likely lacks practical significance. SA, OC, and AS are all located in the city center of Oxford and can therefore be regarded as a single site. The isotopic values of this single site span evenly across a range of 4–10‰, similar to the pattern observed for sites L and SG in Roman southern England.

**Fig. 51.**
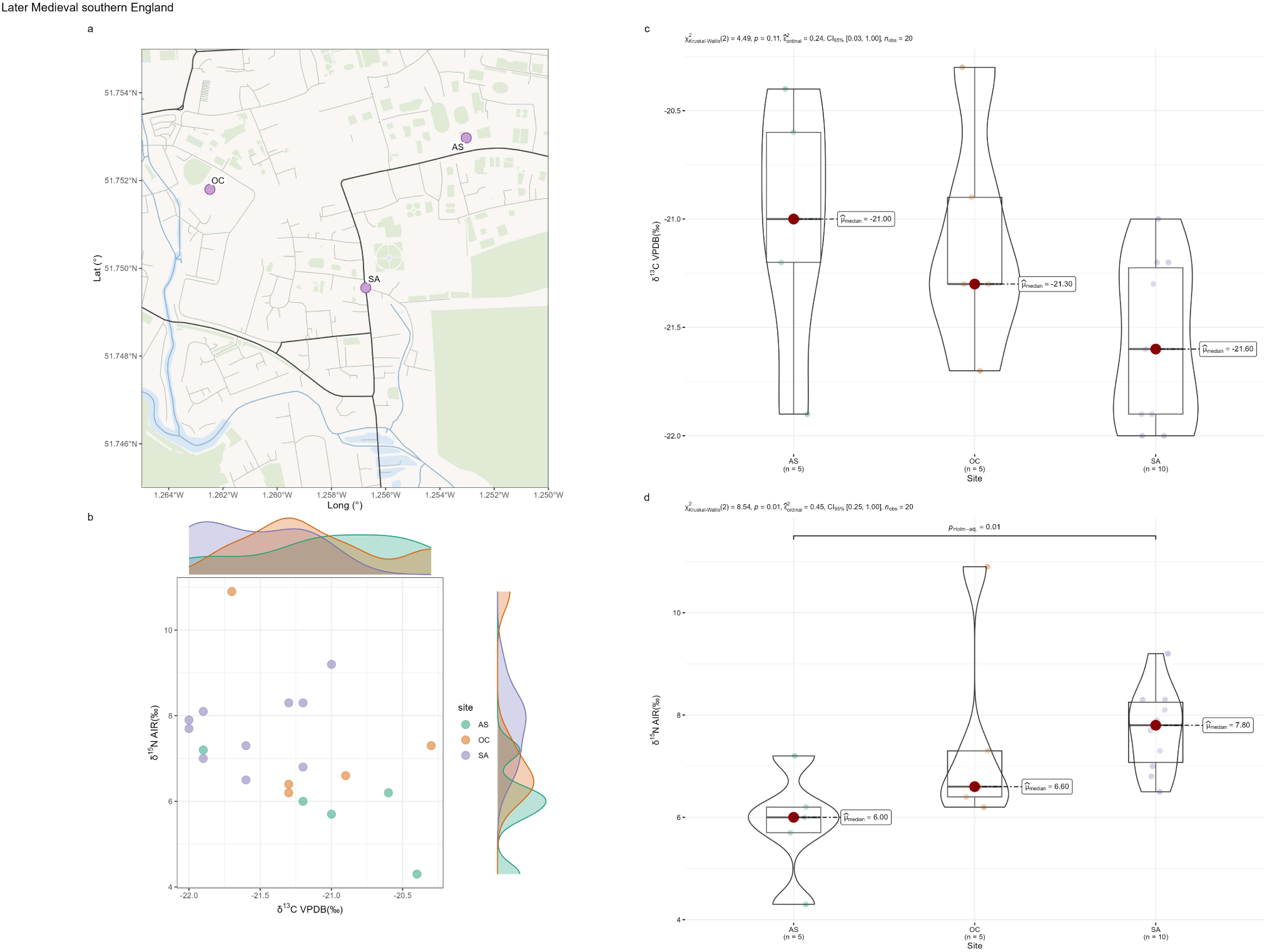
Intra-group comparison of terrestrial omnivores from Later Medieval southern England. **a,** Distribution of sites. **b,** Scatter plot. **c,** Statistical tests for δ^13^C. **d,** Statistical tests for δ^15^N. Site names: AS: All Saints; OC: Oxford Castle; SA: St Aldate’s LAPS.

In northern England, δ^13^C values from WP differ significantly from those of other sites (Fig. 52e). In addition, the post hoc test of δ^15^N between C and F yields a low p-value (Fig. 52f). Yet, C and F are geographically almost indistinguishable (Fig. 52b), and WP is also located close to the sites with which it shows significant post hoc differences (TR and C) (Fig. 52a). These findings suggest that the apparent differences are artifacts of limited sample sizes rather than meaningful isotopic variation. These variations can all be regarded as reasonable variation within a single site. If TR, S, C, and F are treated as one site, their δ^15^N values would cover a continuous range of 4.5–12‰. This once again suggests that with an adequate sample size, the isotopic coverage of one site will become wide. This may result in only minor differences between sites. Consistently, isotopic values from sparsely sampled sites in northern England fall within the cluster defined by the well-sampled sites (Fig. 52d).

**Fig. 52.**
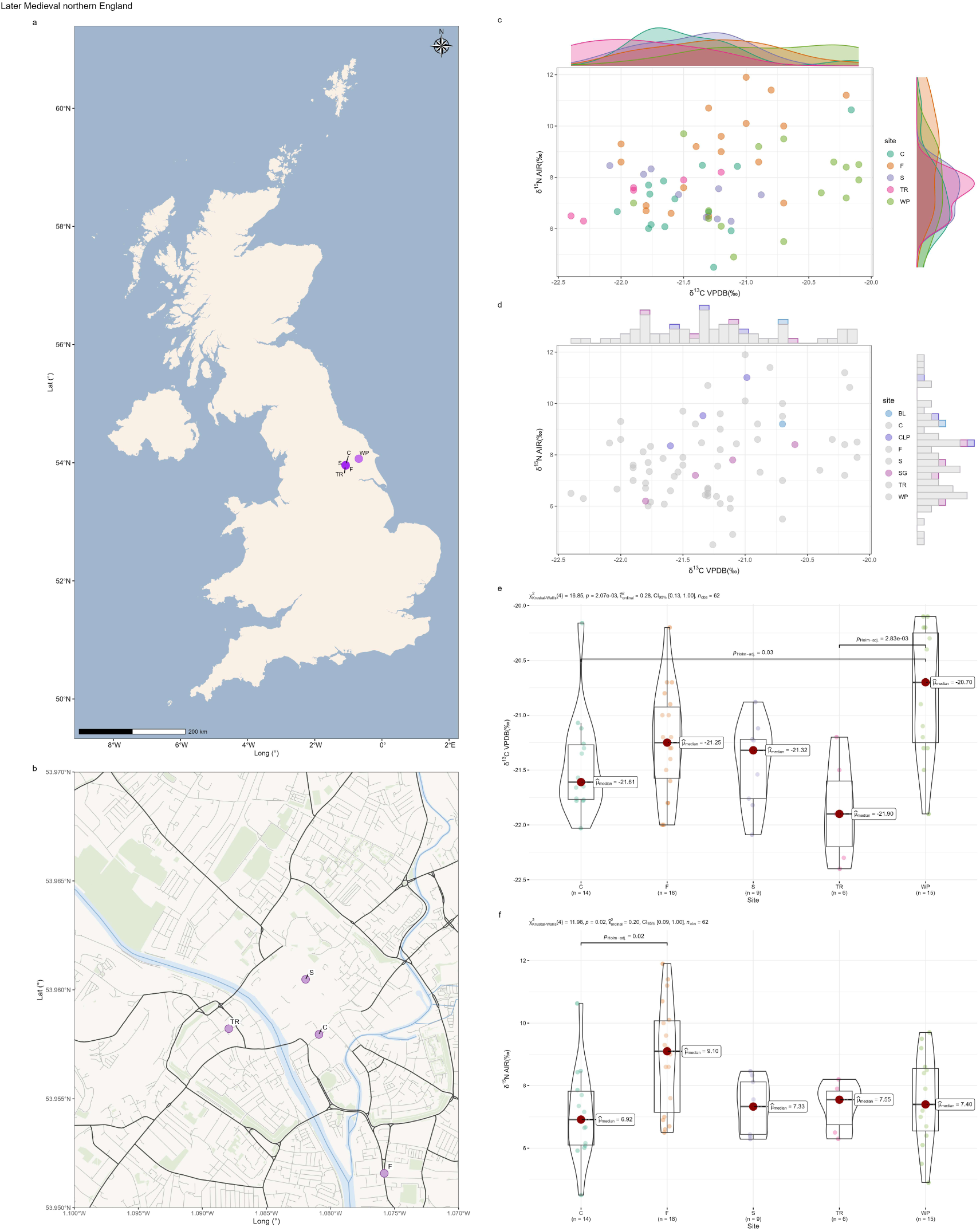
Intra-group comparison of terrestrial omnivores from Later Medieval northern England. **a,** Distribution of sites. **b,** Distribution of four proximate sites. **c,** Scatter plot (well-sampled sites). **d,** Scatter plot (sparsely sampled sites). **e,** Statistical tests for δ^13^C (well-sampled sites). **f,** Statistical tests for δ^15^N (well-sampled sites). Site names: C: Coppergate; F: Fishergate; S: Swinegate; TR: Tanner Row; WP: Wharram Percy; BL: Box Lane; CLP: Clavering Place; SG: St. Giles.

The situation in Scotland is broadly similar to that in northern England. One site (P) differs significantly from the others in δ^15^N (Fig. 53e). However, the sites here have even smaller sample sizes than those in northern England, with the largest site (CP) represented by only eight samples. Thus, the statistical results may have limited practical meaning. The isotopic values of the remaining sparsely sampled site fall entirely within the main cluster defined by the well-sampled sites. (Fig. 53b). We therefore conclude that the sites belonging to the same combined group in the Later Medieval period exhibit broadly similar isotopic values for omnivores.

**Fig. 53.**
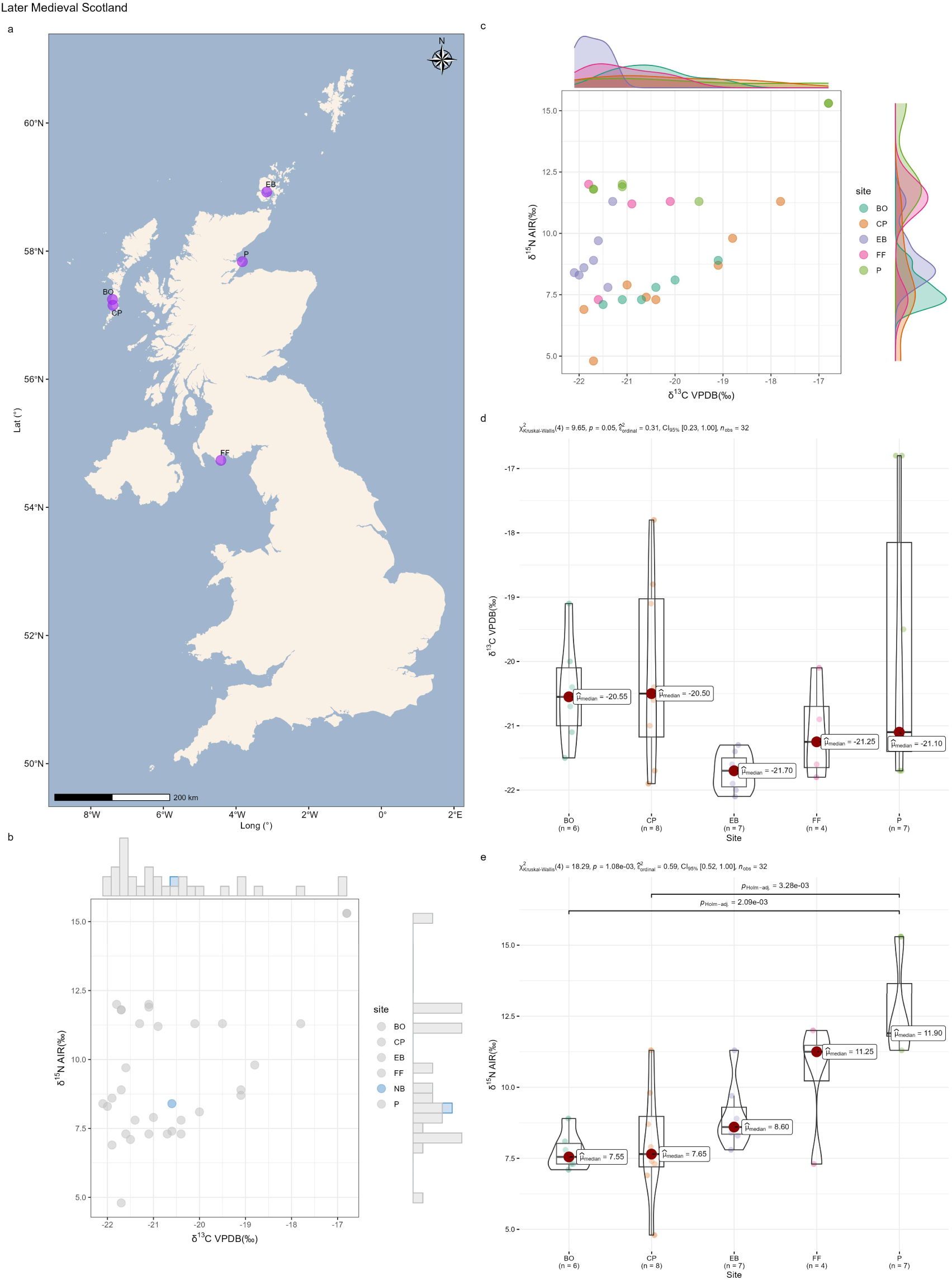
Intra-group comparison of terrestrial omnivores from Later Medieval Scotland. **a,** Distribution of sites. **b,** Scatter plot (sparsely sampled sites). **c,** Scatter plot (well-sampled sites). **d,** Statistical tests for δ^13^C (well-sampled sites). **e,** Statistical tests for δ^15^N (well-sampled sites). Site names: BO: Bornais; CP: Cille Phedair; EB: Earl’s Bu; FF: Fey Field; P: Portmahomack; NB: Site of Newark Bay.

Inter-group comparisons show patterns that are somewhat inconsistent with those observed in earlier periods (Fig. 54). The isotopic values of samples from southern England, northern England, and Scotland largely overlap, although a few Scottish samples fall outside the main cluster. The Welsh samples display a slightly distinct isotopic distribution, suggesting a modest degree of regional differentiation, which differs from the pattern observed in earlier periods. Data for the Post Medieval period is extremely limited, and although isotopic values are plotted in Fig. 55, the sample size is too small to permit meaningful conclusions.

**Fig. 54.**
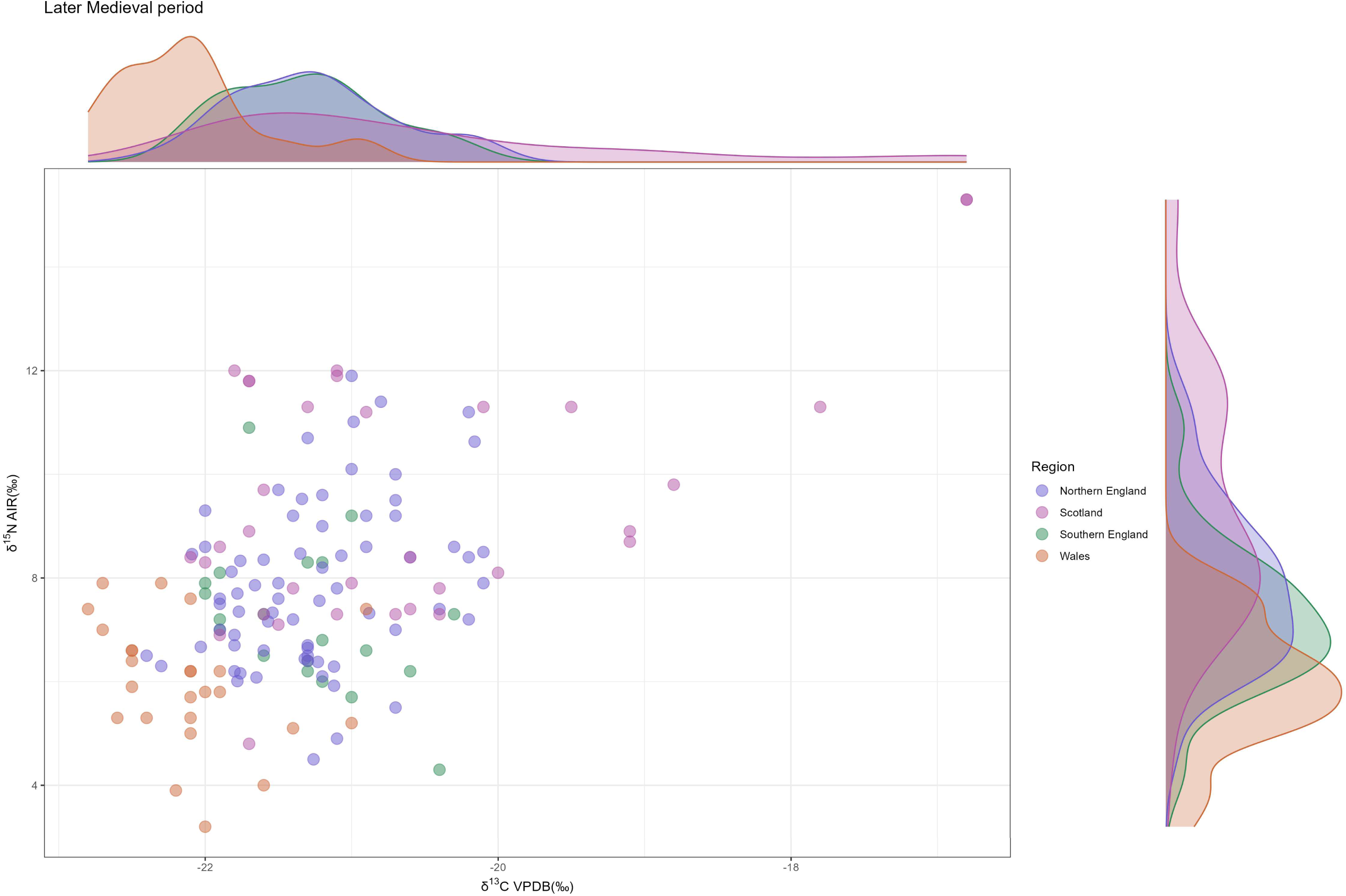
Inter-group comparison of terrestrial omnivores from the Later Medieval period (Scatter plot).

**Fig. 55.**
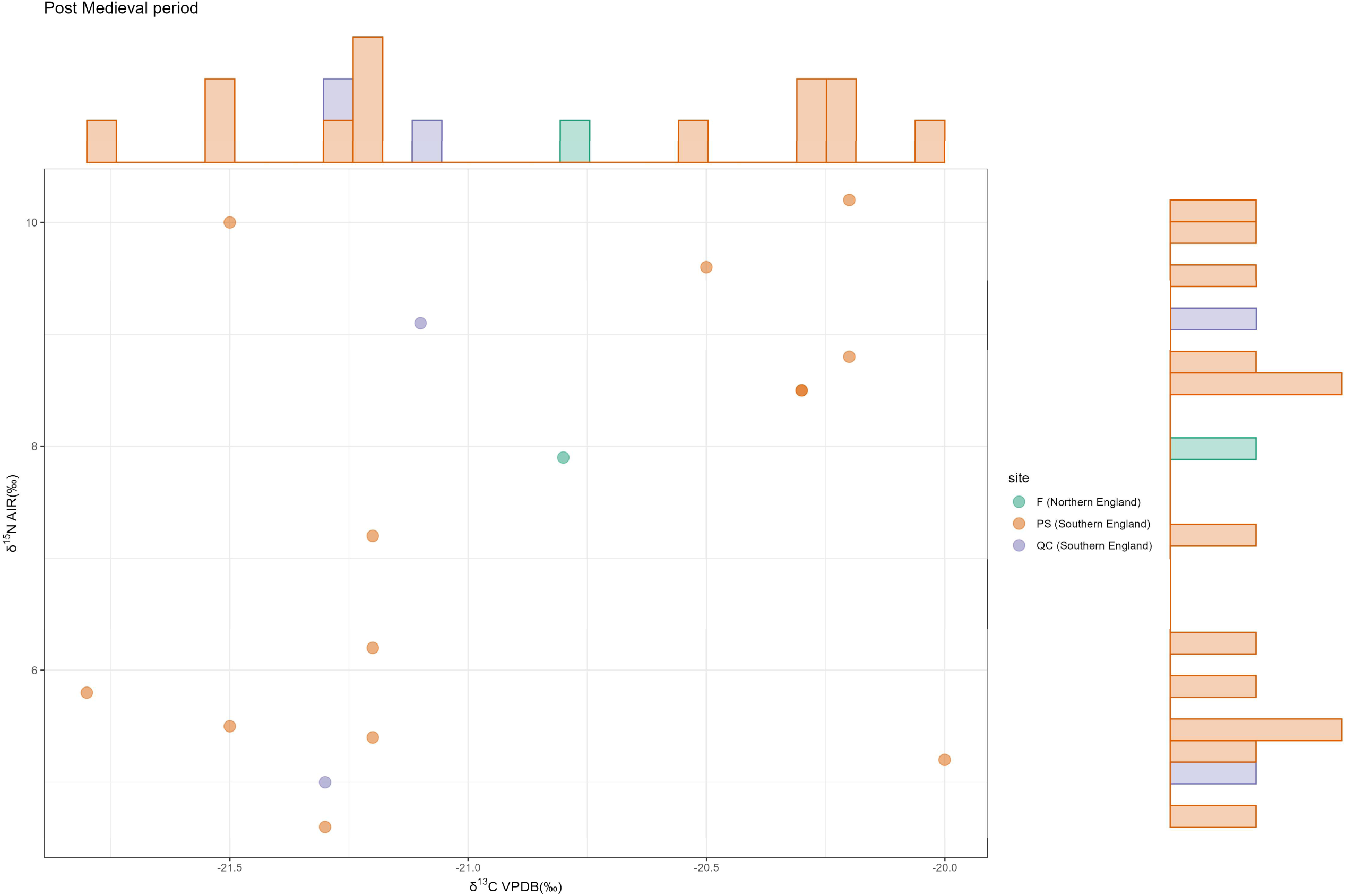
Isotopic comparison of terrestrial omnivores from the Post Medieval period (Scatter plot).

### Analysis of isotopic variation for terrestrial plants, marine fish, and freshwater fish

#### Plant data

The overall isotopic data for plants, organized by region–period combined groups, are shown in Fig. 56. One of the most striking patterns is that the isotopic distribution of the region-period combined group with the largest sample size, Neolithic Scotland (n = 206), forms an isotopic envelope that almost entirely encompasses the distributions of all other combined groups (n = 194). When Neolithic Scotland is excluded, it is evident that some groups are clearly separated—for example, Neolithic Wales, Early Medieval central England, and Iron Age southern England. By contrast, some groups show substantial overlap, such as Early Medieval versus Later Medieval central England, and Iron Age versus Roman southern England. This pattern may suggest that plant isotopic values vary more strongly among regions than among periods. However, these patterns should be interpreted cautiously, as they may partly reflect small sample sizes, which can underestimate the full range of isotopic variability within each group. With a sufficiently large sample size, as illustrated by Neolithic Scotland, a single combined group can encompass much of the isotopic variation observed across all other combined groups. To examine this pattern more closely, we conducted more fine-grained analyses by comparing isotopic data across periods while controlling for region, and across regions while controlling for period.

**Fig. 56.**
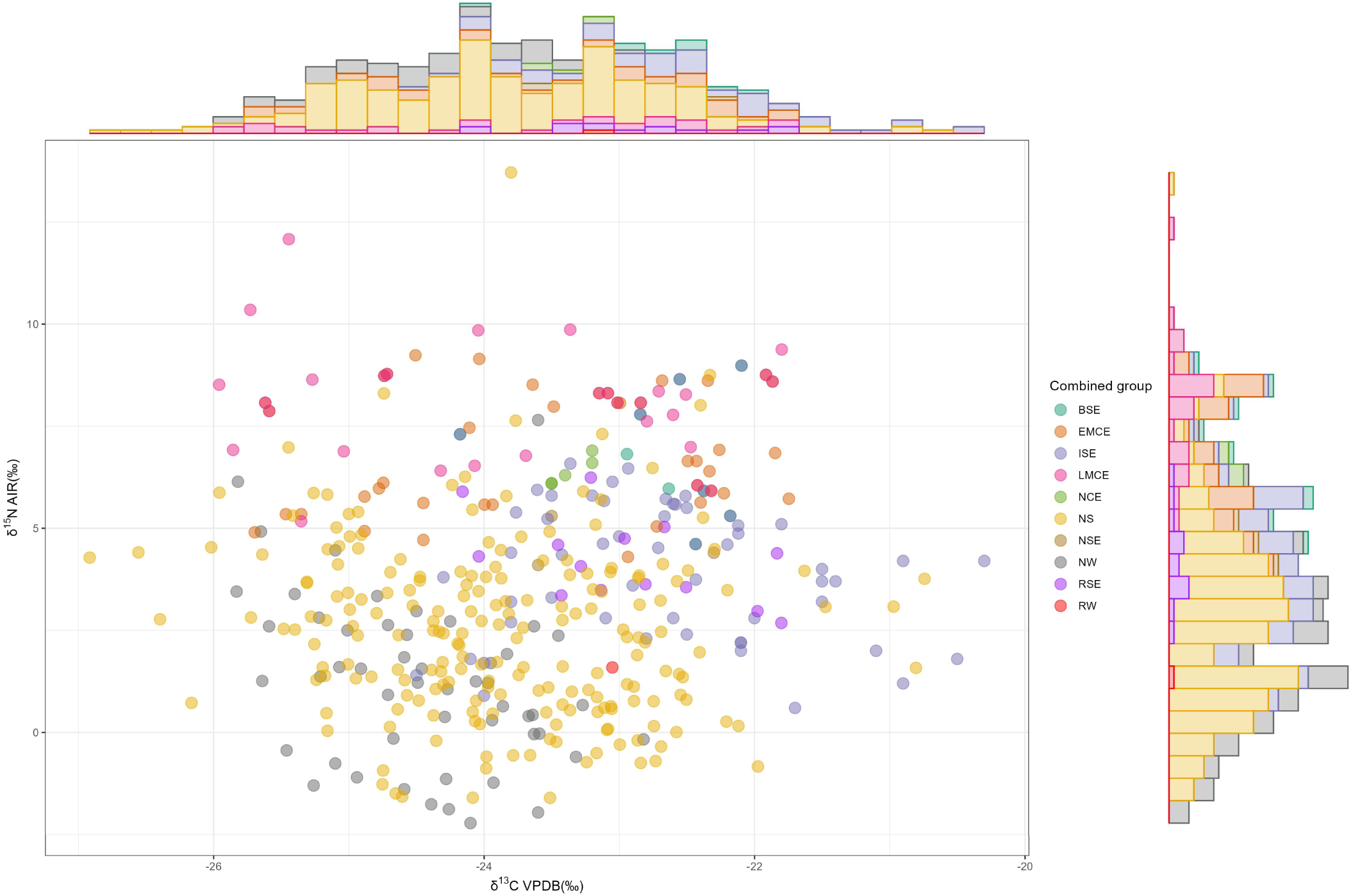
Isotopic comparison of plant data from different region-period combined groups (Scatter plot). Group names: BSE: Bronze Age southern England; EMCE: Early Medieval central England; ISE: Iron Age southern England; LMCE: Later Medieval central England; NCE: Neolithic central England; NS: Neolithic Scotland; NSE: Neolithic southern England; NW: Neolithic Wales; RSE: Roman southern England; RW: Roman Wales.

1) Comparison of isotopic data across periods while controlling for region

Firstly, we compared isotopic values of region–period groups within the same region across different periods. Such analyses were possible for southern England (Neolithic, Bronze, Iron, Roman) and central England (Neolithic, Early Medieval, Later Medieval) (Fig. 57 and Fig. 58). The scatter plots show that groups from different periods are largely intermixed, a pattern confirmed by the statistical tests. For δ^13^C, both regions yield very high p-values. For δ^15^N, a small number of post hoc tests show low p-values (southern England: Bronze Age vs. Iron Age, Bronze Age vs. Roman period) (Fig. 57c). However, the group showing significant differences (BSE) includes only nine samples, all of which fall within the range of the ISE (59 samples). We therefore regard these low p-values as artifacts of sample size. Overall, our data indicate that isotopic values from different periods within the same region do not differ significantly, although δ^15^N may show somewhat greater variability than δ^13^C.

**Fig. 57.**
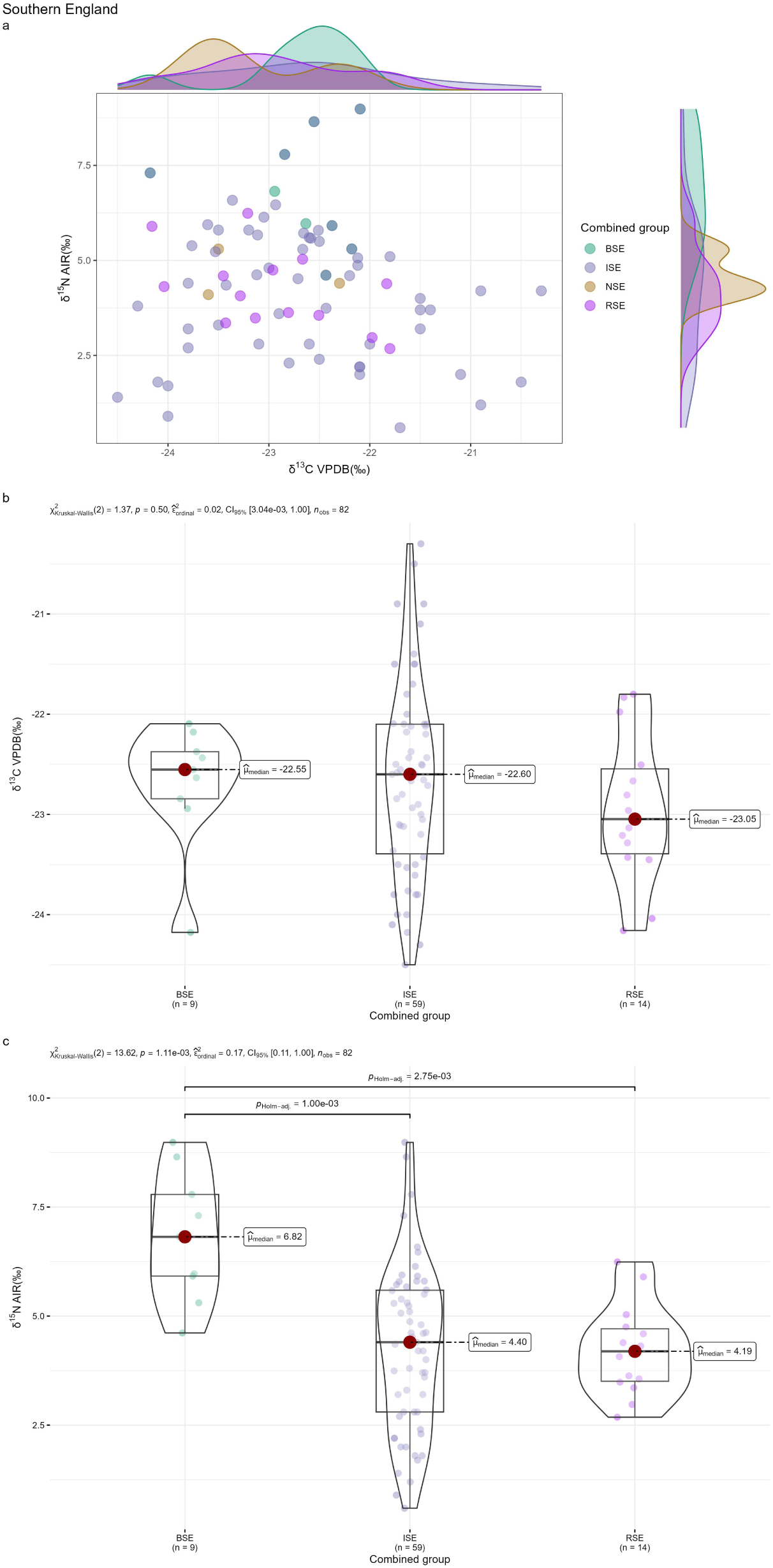
Isotopic comparison of plant data from the different periods but the same region: Southern England. **a,** Scatter plot; **b,** Statistical test for δ^13^C; **c,** Statistical test for δ^15^N. Group names: BSE: Bronze Age southern England; ISE: Iron Age southern England; NSE: Neolithic southern England; RSE: Roman southern England.

**Fig. 58.**
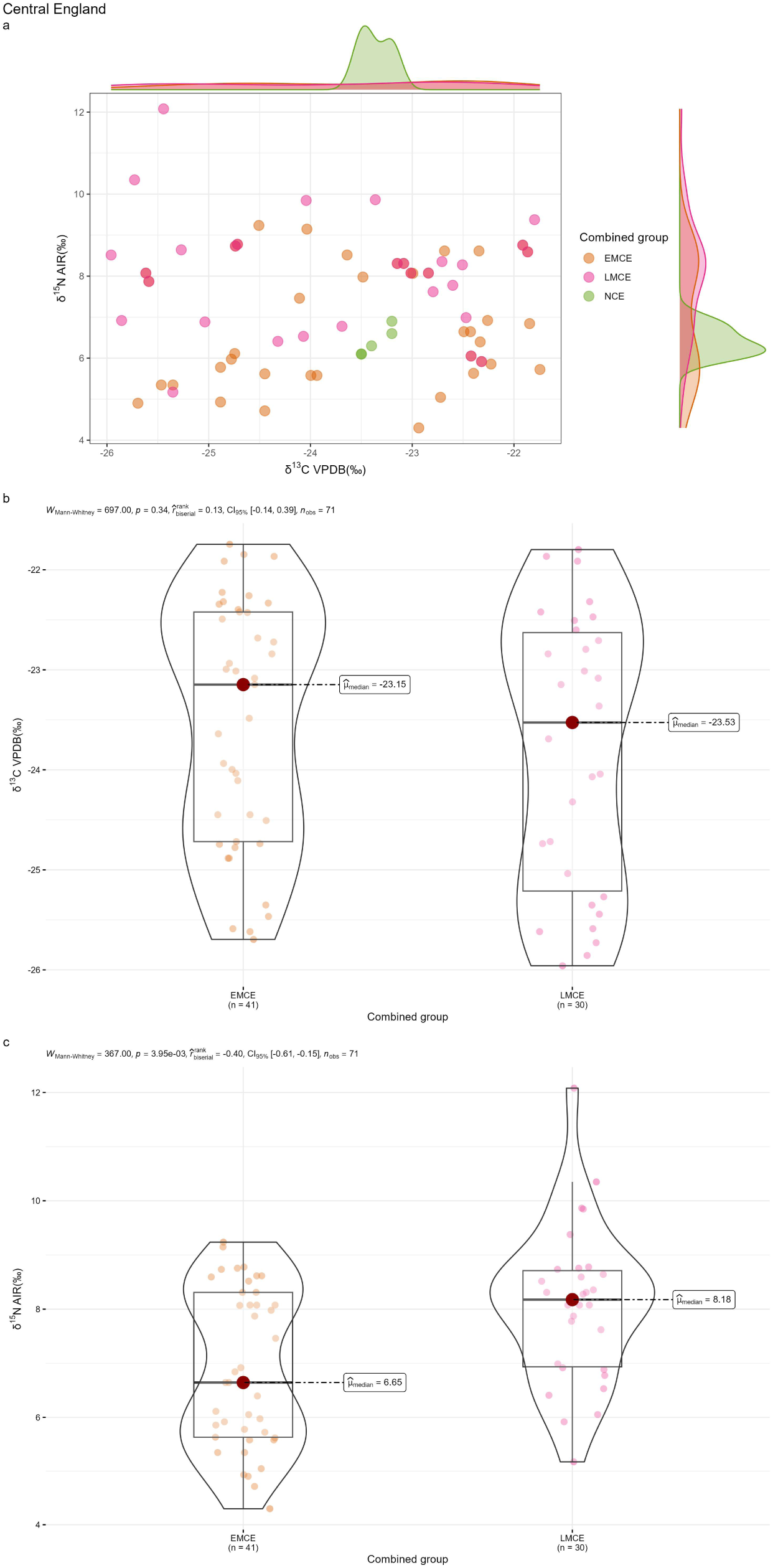
Isotopic comparison of plant data from the different periods but the same region: Central England. **a,** Scatter plot; **b,** Statistical test for δ^13^C; **c,** Statistical test for δ^15^N. Group names: EMCE: Early Medieval central England; LMCE: Later Medieval central England; NCE: Neolithic central England.

2) Comparison of isotopic data across regions while controlling for period

We then compared region–period groups from the same period across different regions. This analysis was possible for the Neolithic (central England, Scotland, southern England, Wales) and the Modern period (Scotland, southern England). The Neolithic comparison is relatively straightforward: the broad isotopic distribution of Neolithic Scotland completely encompasses the distributions of the other Neolithic groups (Fig. 59a). The modern plant data provide a different picture, with southern England and Scotland showing significantly different isotopic distributions (Fig. 59b). A similar pattern is observed in the scatter plot of modern soils from Scotland and southern England (Fig. 59c). For the Neolithic data, all groups except Scotland have small sample sizes, and this sample-size imbalance prevents us from drawing stronger conclusions. In contrast, the modern dataset has large sample sizes for both Scotland and southern England, making the conclusion drawn from their comparison more reliable. Their comparison shows clear regional differences in isotopic distributions. Overall, plant isotopic values demonstrate strong inter-regional variability but remain relatively stable through time within each region.

**Fig. 59.**
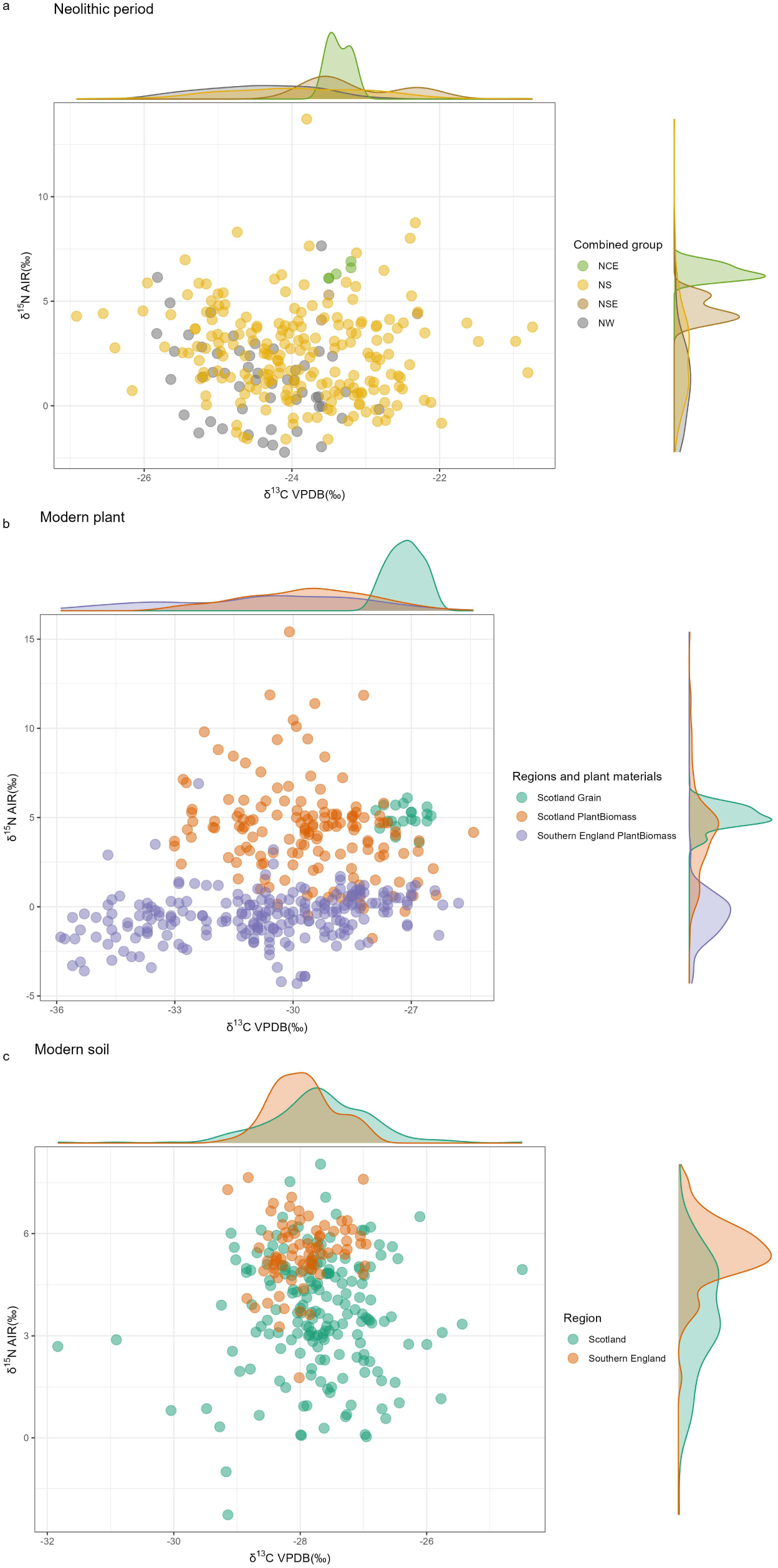
Isotopic comparison of plant and soil data from the different regions but the same period (Scatter plots). **a,** Neolithic period. **b,** Modern plant. **c,** Modern soil. Group names of **a**: NCE: Neolithic central England; NS: Neolithic Scotland; NSE: Neolithic southern England; NW: Neolithic Wales.

### Marine fish data

The overall pattern of marine fish isotopic values is shown in Fig. 60. The most striking feature is that all values from other groups fall within the cluster defined by the two groups with the largest sample sizes (LMNE and LMSE). Moreover, the isotopic values of these two large groups overlap extensively. This suggests that the isotopic composition of marine fish does not vary significantly across regions or periods. To further test this conclusion, we applied the same approach used for plants, controlling for region and period separately in the comparisons.

**Fig. 60.**
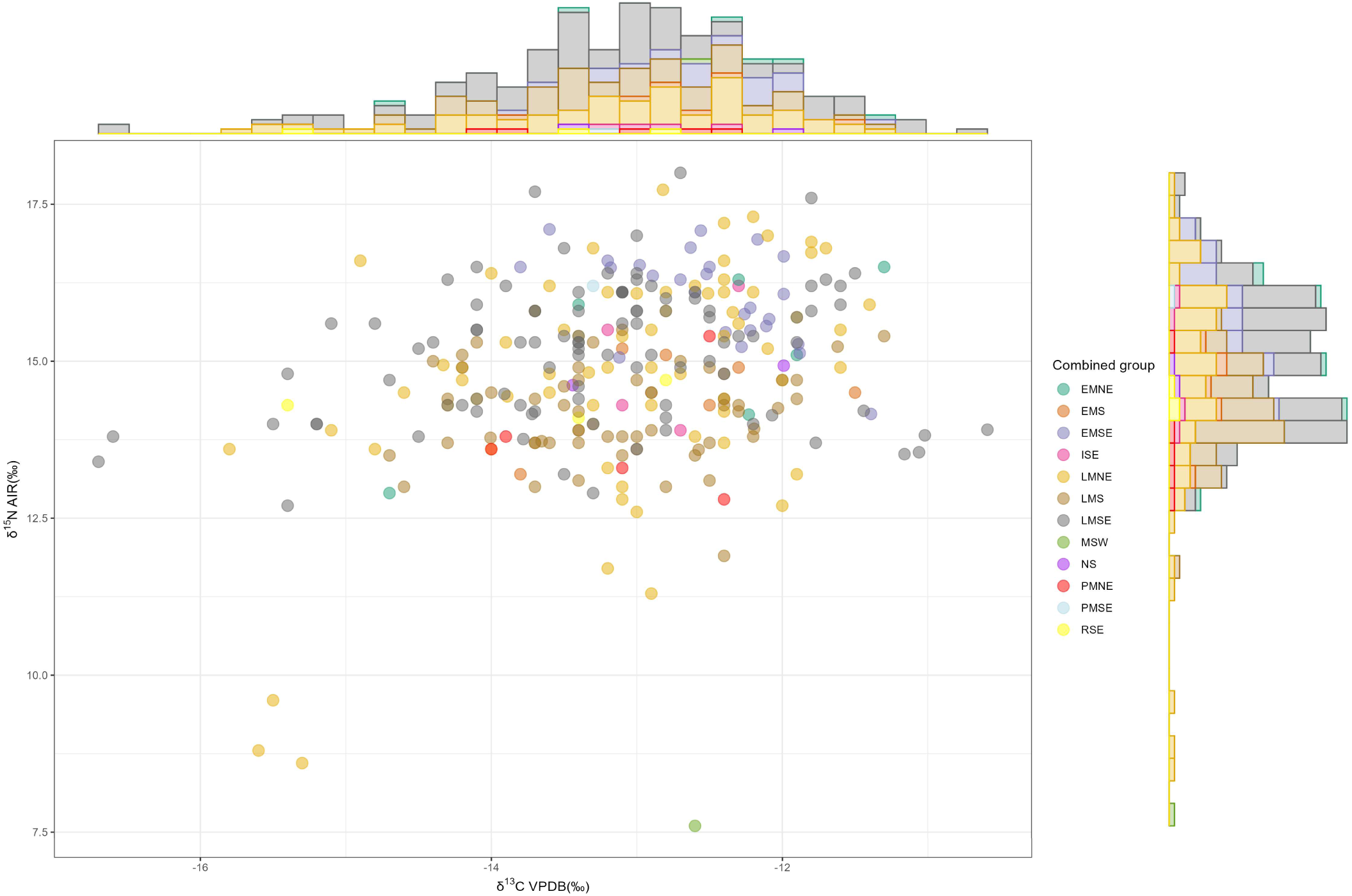
Isotopic comparison of marine fish data from different region-period combined groups (Scatter plot). Group names: EMNE: Early Medieval northern England; EMS: Early Medieval Scotland; EMSE: Early Medieval southern England; ISE: Iron age southern England; LMNE: Later Medieval northern England; LMS: Later Medieval Scotland; LMSE: Later Medieval southern England; MSW: Mesolithic Wales; NS: Neolithic Scotland; PMNE: Post Medieval northern England; PMSE: Post Medieval southern England; RSE: Roman southern England.

1) Comparison of isotopic data across periods while controlling for region

According to our marine fish dataset, period-based comparisons can be made for southern England (Iron Age, Roman, Early Medieval, Later Medieval), Scotland (Early Medieval, Later Medieval), and northern England (Early Medieval, Later Medieval, Post-Medieval) (Fig. 61). In all three scatter plots, isotopic data from different periods show no clear separation, with groups with smaller sample sizes falling within the clusters defined by the larger ones. We therefore conclude that the isotopic values of marine resources do not vary significantly over time.

**Fig. 61.**
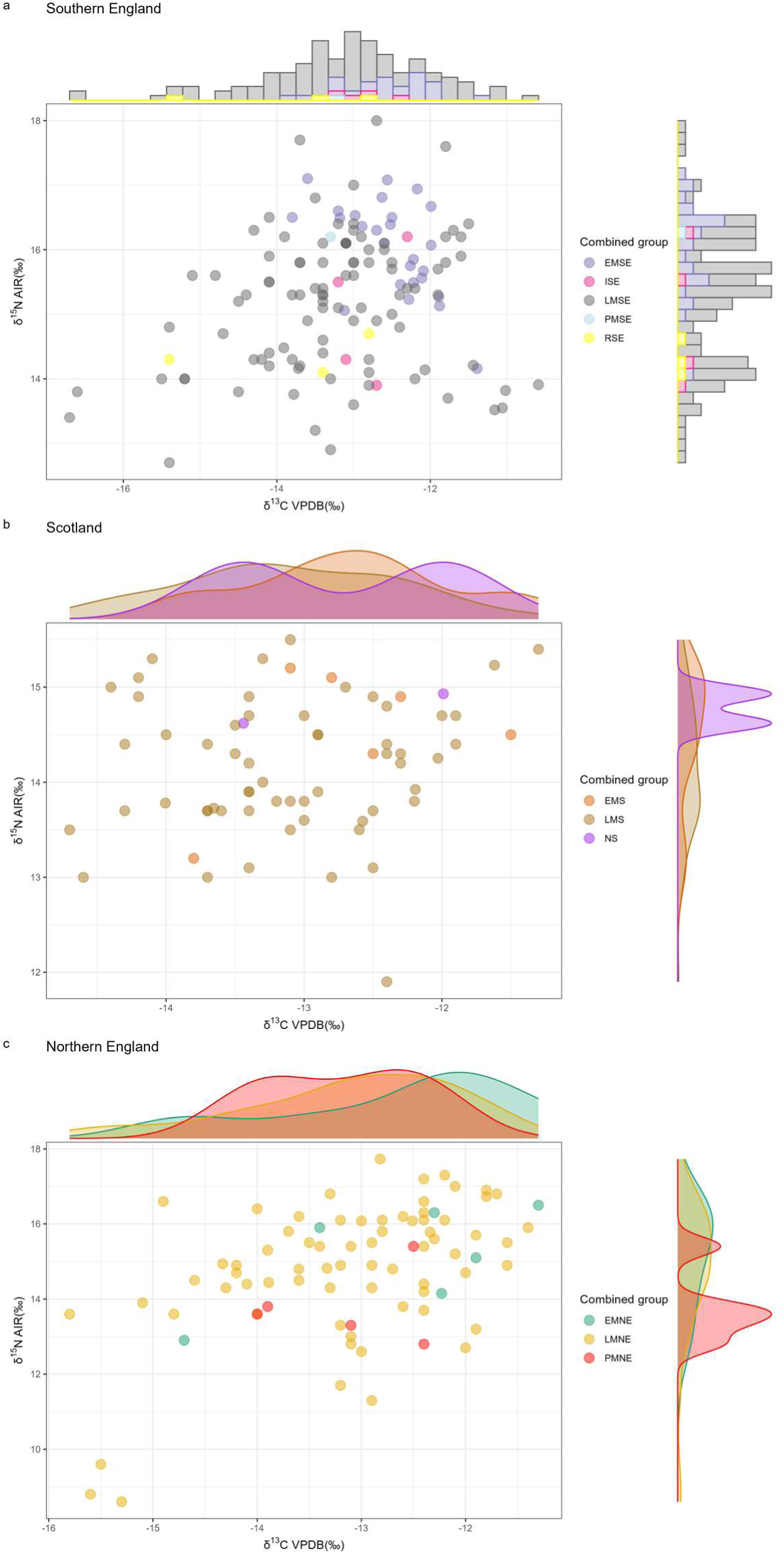
Isotopic comparison of marine fish data from the different periods but the same region (Scatter plots). **a,** Southern England. **b,** Scotland. **c,** Northern England. Group names: EMNE: Early Medieval northern England; EMS: Early Medieval Scotland; EMSE: Early Medieval southern England; ISE: Iron age southern England; LMNE: Later Medieval northern England; LMS: Later Medieval Scotland; LMSE: Later Medieval southern England; NS: Neolithic Scotland; PMNE: Post Medieval northern England; PMSE: Post Medieval southern England; RSE: Roman southern England.

2) Comparison of isotopic data across regions while controlling for period Comparisons of different regions within the same period were possible for two periods: Early Medieval (northern England, Scotland, southern England) and Later Medieval (northern England, Scotland, southern England) (Fig. 62 and Fig. 63). For the Early Medieval period, the scatter plot shows that data from groups with smaller sample sizes (EMNE and EMS) fall within the cluster defined by the group with the larger sample size (EMSE). The significant post hoc tests are likely driven by differences in sample size (EMS: 6 samples; EMSE: 26 samples) (Fig. 62c). A similar pattern is observed for the Later Medieval data (Fig. 63). We therefore conclude that the isotopic values of marine resources do not vary geographically.

**Fig. 62.**
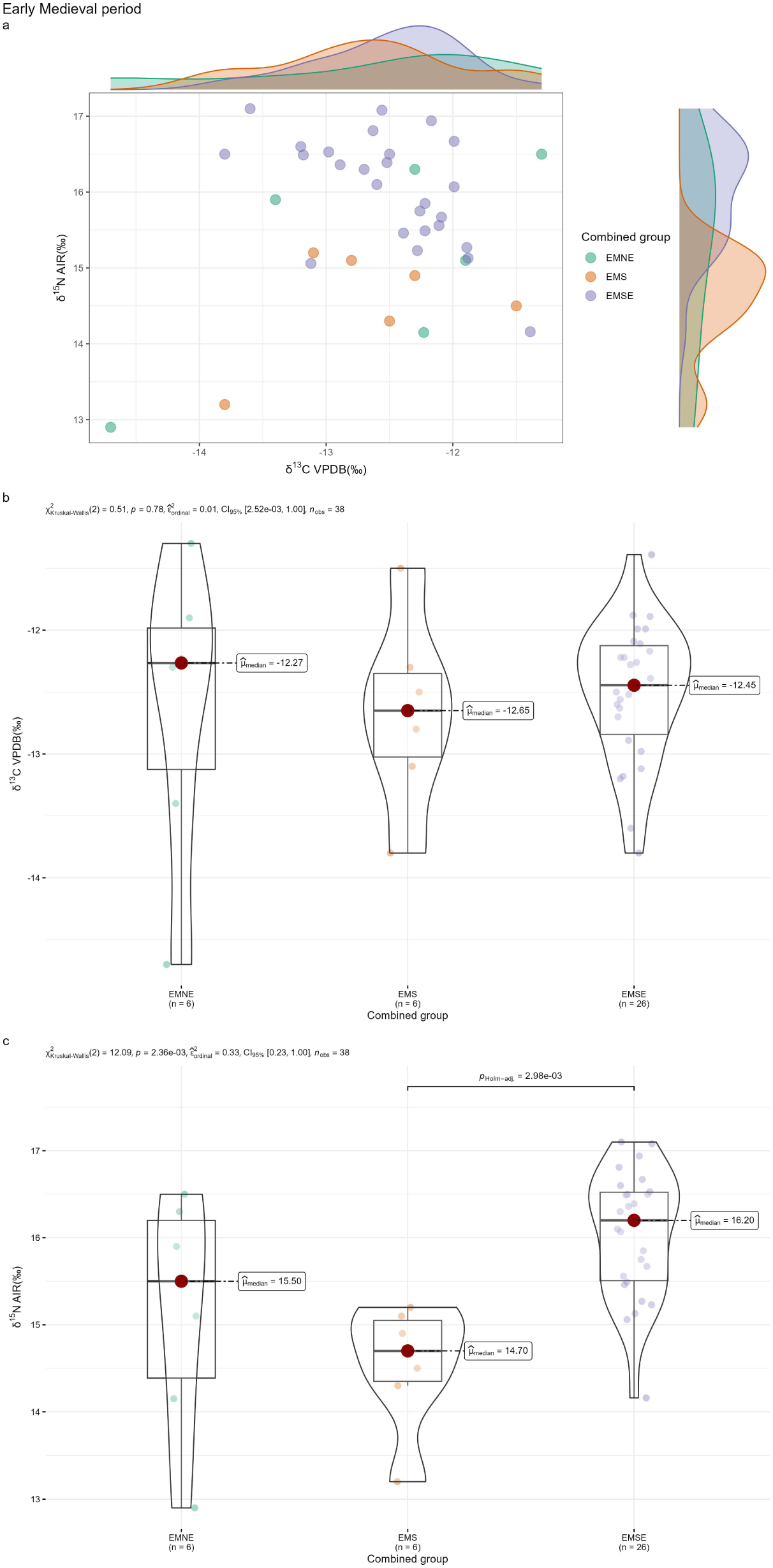
Isotopic comparison of marine fish data from the different regions but the same period: Early Medieval period. **a,** Scatter plot; **b,** Statistical test for δ^13^C; **c,** Statistical test for δ^15^N. Group names: EMNE: Early Medieval northern England; EMS: Early Medieval Scotland; EMSE: Early Medieval southern England.

**Fig. 63.**
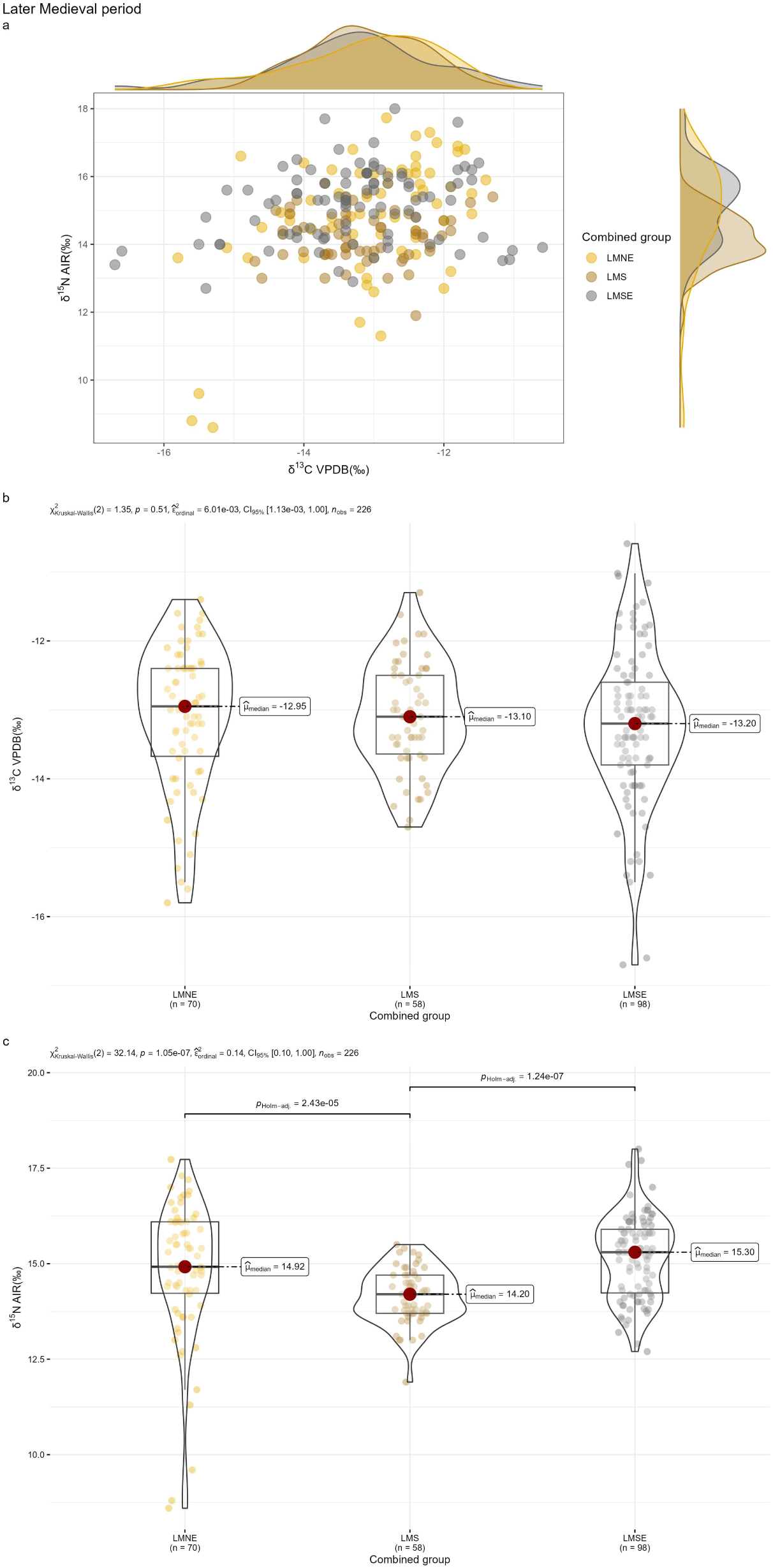
Isotopic comparison of marine fish data from the different regions but the same period: Later Medieval period. **a,** Scatter plot; **b,** Statistical test for δ^13^C; **c,** Statistical test for δ^15^N. Group names: LMNE: Later Medieval northern England; LMS: Later Medieval Scotland; LMSE: Later Medieval southern England.

### Freshwater fish data

The overall pattern of freshwater fish isotopic values is shown in Fig. 64. The scatter plot indicates that data from different region–period groups cluster distinctly, particularly LMNE, ISE, EMS, and LMSE. However, the isotopic ranges of the sparsely sampled groups are largely encompassed by that of the group with the largest sample size (LMNE). Even within a single site, freshwater fish samples can span a considerable isotopic range. This suggests that freshwater fish resources may not differ substantially in isotopic composition across regions or periods, although the limited number of samples makes this conclusion uncertain.

**Fig. 64.**
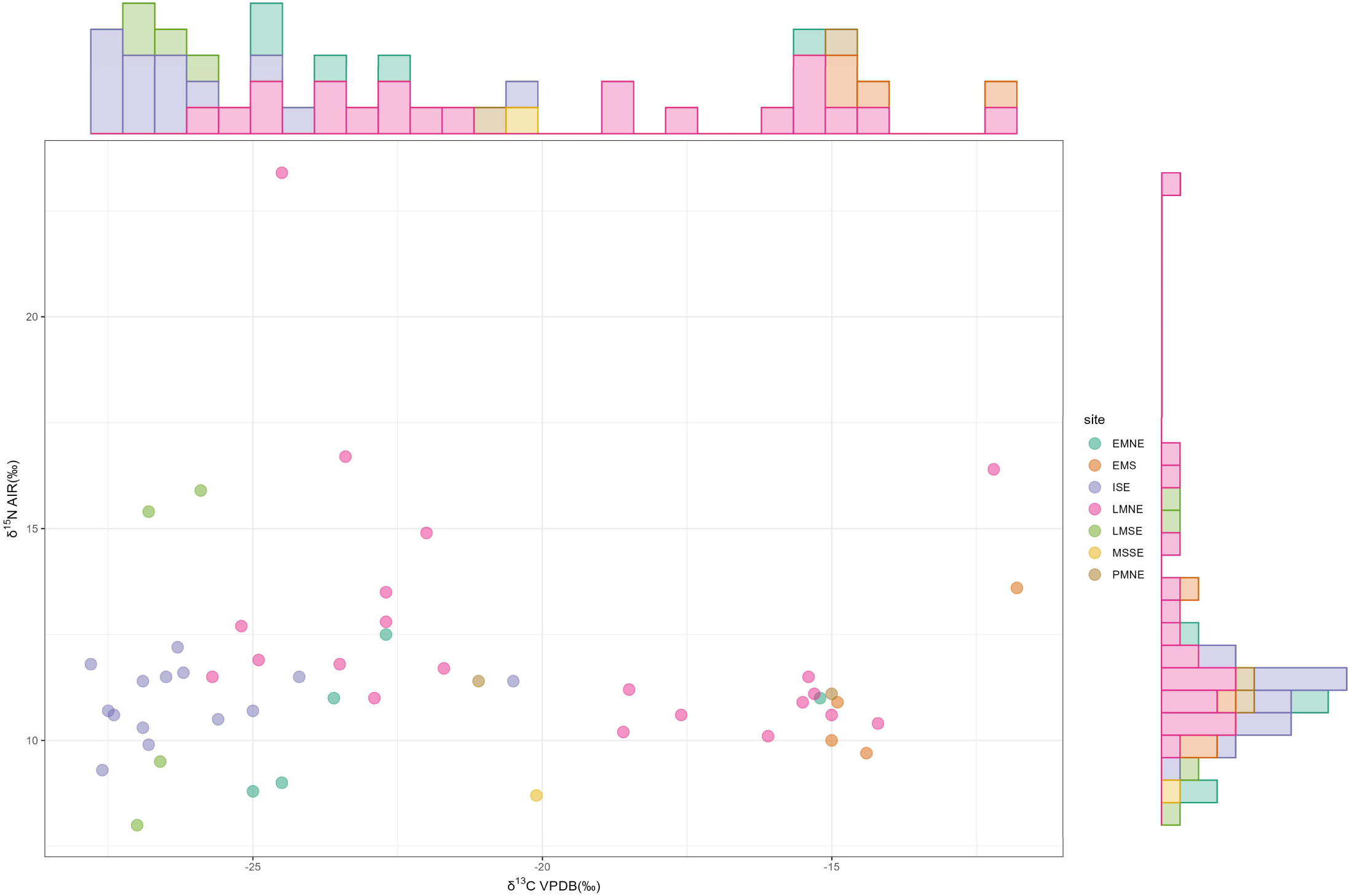
Isotopic comparison of freshwater fish data (Scatter plot). Group names: EMNE: Early Medieval northern England; EMS: Early Medieval Scotland; ISE: Iron Age southern England; LMNE: Later Medieval northern England; LMSE: Later Medieval southern England; MSSE: Mesolithic southern England; PMNE: Post Medieval northern England.

## Discussion

The preceding analyses indicate that the isotopic values of terrestrial herbivores and omnivores do not differ significantly among sites within each combined group. Although the statistical tests for some combined groups yield significant results, these are likely attributable to insufficient and uneven sample sizes. Most sites within each combined group contain fewer than 20 samples, and some even fewer than 10. Such small sample sizes represent a very limited subset of the food resources that would have been available at a site and are therefore highly susceptible to sampling bias[96]. Moreover, the sample sizes vary substantially among sites, showing considerable inequality in representation. We then provide a detailed examination of how small and uneven sample sizes affect statistically significant results from three perspectives: the geographical distribution of sites, the magnitude of isotopic differences among sites, and the isotopic scatterplot patterns contrasting sites with small and large sample sizes.

Firstly, in combined groups where statistically significant results were detected, the pairwise significant differences among sites revealed by post hoc tests seldom follow any clear geographical pattern. Soils and plants from geographically proximate sites generally exhibit comparable isotopic signatures because they are exposed to similar climatic, environmental, and ecological factors. This geographical isotopic pattern is transferred through food webs and consequently reflected in animals[97,98]. However, contrary to this expectation, many of our results show that isotopic values differ significantly between geographically proximate sites, whereas more distant sites exhibit similar isotopic signatures. It is particularly puzzling that isotopic differences are sometimes observed between sites in very close geographical proximity (e.g., Fig. 9), which are close enough to be regarded as a single location. In every instance of this kind, the sites involved have small sample sizes, typically fewer than 20. The most plausible explanation is that these isotopic differences arise from sampling bias rather than genuine inter-site variation. They likely reflect variability within sites rather than meaningful differences between them.

The second situation concerns isotopic differences that are statistically significant but too small to be of practical significance. For instance, in Iron Age southern England, pairwise comparisons among 16 sites reveal several significant contrasts, yet the largest δ^13^C difference between sites is only 0.97‰ (Fig. 17). Such a small difference suggests that the statistically significant results have little substantive meaning. After all, such a discrepancy can arise even for the same sample when analyzed in different laboratories, owing to variations in sample preparation procedures, analytical instruments, and data calibration[99].

The third situation arises when pairwise comparisons yield statistically significant results, but the sample sizes among sites are extremely uneven. Two main situations can be identified. The first occurs when a statistically significant result is driven primarily by a special site represented by fewer than 10 samples, a sample size at which sampling bias is highly likely[96]. However, the isotopic values of this special site still fall entirely within the range defined by the other sites in the scatterplot (e.g., Fig. 33). This clearly indicates that the apparent significance is a consequence of sampling bias rather than genuine inter-site variation. The second situation is more complex. The sample sizes vary widely among sites, and a large number of pairwise post hoc comparisons yield significant results. When the isotopic values are visualized in scatterplots, it becomes clear that data points from sites with smaller sample sizes all lie within the range defined by those with larger sample sizes. A particularly representative example is the SF site in Iron Age southern England, which contains 217 herbivore samples (Fig. 16). The herbivore isotopic values of nearly all other southern England sites fall entirely within the range defined by this site. Other similar herbivore examples include the Y site in Roman southern England (Fig. 24) and the HQ and L sites in Early Medieval southern England (Fig. 30). For omnivores, when two geographically proximate sites (L and SG) in Roman southern England are combined, a similar pattern is also observed (Fig. 45).

This phenomenon raises the question of whether, if the sample size from a single site were sufficiently large (e.g., 500 samples, though practically impossible), its isotopic distribution could approximate the overall isotopic signature of that region (e.g., England, Scotland, Wales). Our analysis partly supports this view. Apart from the sites SF, Y, HQ, and L mentioned above, the Iron Age inter-group comparisons for herbivores further support this view. All samples from northern England came from a single site with a relatively large sample size, yet its isotopic distribution was nearly identical to that of southern England, which comprised samples from 28 different sites (Fig. 22). Thus, our data analysis tends to support the view that isotopic differences among sites within each combined group are not significant. The apparent significant differences are most likely the result of sampling bias caused by small sample sizes at individual sites. This finding strongly suggests that traditional baseline reconstructions relying on samples from one or a few sites may be subject to substantial sampling bias. At present, it remains unclear what constitutes a sufficiently representative sample size. Earlier simulation studies suggested that large discrepancies are likely when fewer than eight samples are used to estimate population means, whereas sampling redundancy tends to occur when more than 40 samples are used[96]. Our results, however, indicate that this estimate may be overly optimistic. In our analysis, only sites represented by more than 50 samples were found to display a sufficiently comprehensive range of isotopic values (e.g., Fig. 16, Fig. 24, and Fig. 30). Unfortunately, the vast majority of sites contain fewer than 20 samples.

Therefore, based on the above analyses, while not without limitations, it is appropriate to use the combined group as the criterion for constructing the isotopic baselines of terrestrial herbivores and omnivores. By adopting this approach, we not only overcome the lack of accessible food-resource data for some sites (e.g., Warrington and Towton[7]) but also partially alleviate the estimation bias that may result from small sample sizes at individual sites. Inter-group comparisons reveal no significant differences among southern, central, and northern England. Across all periods, inter-group comparisons of both herbivores and omnivores yield consistent results. When the sample sizes are comparable, the isotopic values of samples from southern England, central England, and northern England are largely intermixed. When sample sizes differ substantially, the groups with smaller sample sizes tend to fall within the isotopic range of those with larger sample sizes. For this reason, we integrated data from southern England, central England, and northern England within the same period into a single group. In addition, because of the small sample size for the post-Medieval period, we combined Later Medieval and post-Medieval samples. The temporal gap between these periods is not large, and our analysis shows that isotopic values from the same site but slightly different phases do not differ substantially (Fig. 3, Fig. 5, Fig. 9, Fig. 12, Fig. 16, Fig. 39, and Fig. 41). Such a combination is therefore reasonable. The final combined groups are shown in Table 1. For terrestrial herbivores and omnivores, accessible food resources are newly defined at the level of region–period combined groups. Accordingly, terrestrial herbivores and omnivores from a given combined group can be used as accessible food resources for human individuals assigned to the same group. The sample sizes, mean δ^13^C and δ^15^N values, and standard deviations for each combined group are also presented in Table 1. As shown in Table 1, even after applying these groupings, some groups still have very small sample sizes. For terrestrial herbivores, these include Mesolithic England; Paleolithic, Mesolithic, Neolithic, and Roman Wales; and Roman Iron Age Scotland. For terrestrial omnivores, these include Paleolithic and Mesolithic England; Paleolithic, Mesolithic, Neolithic, Roman, and Early Medieval Wales; and Roman Iron Age Scotland. These groups should therefore be interpreted with caution when used as accessible food resources.

**Table 1.**
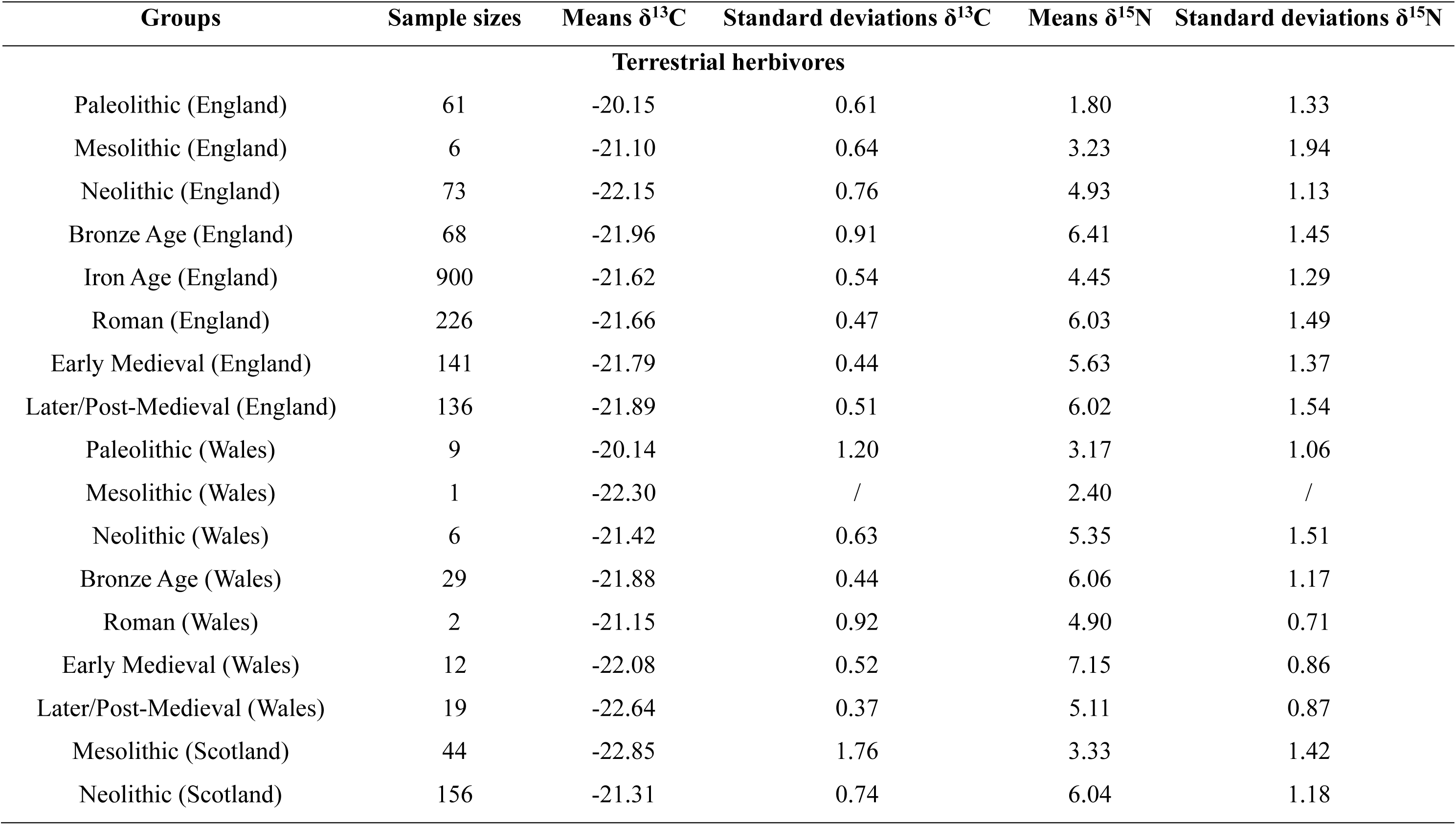
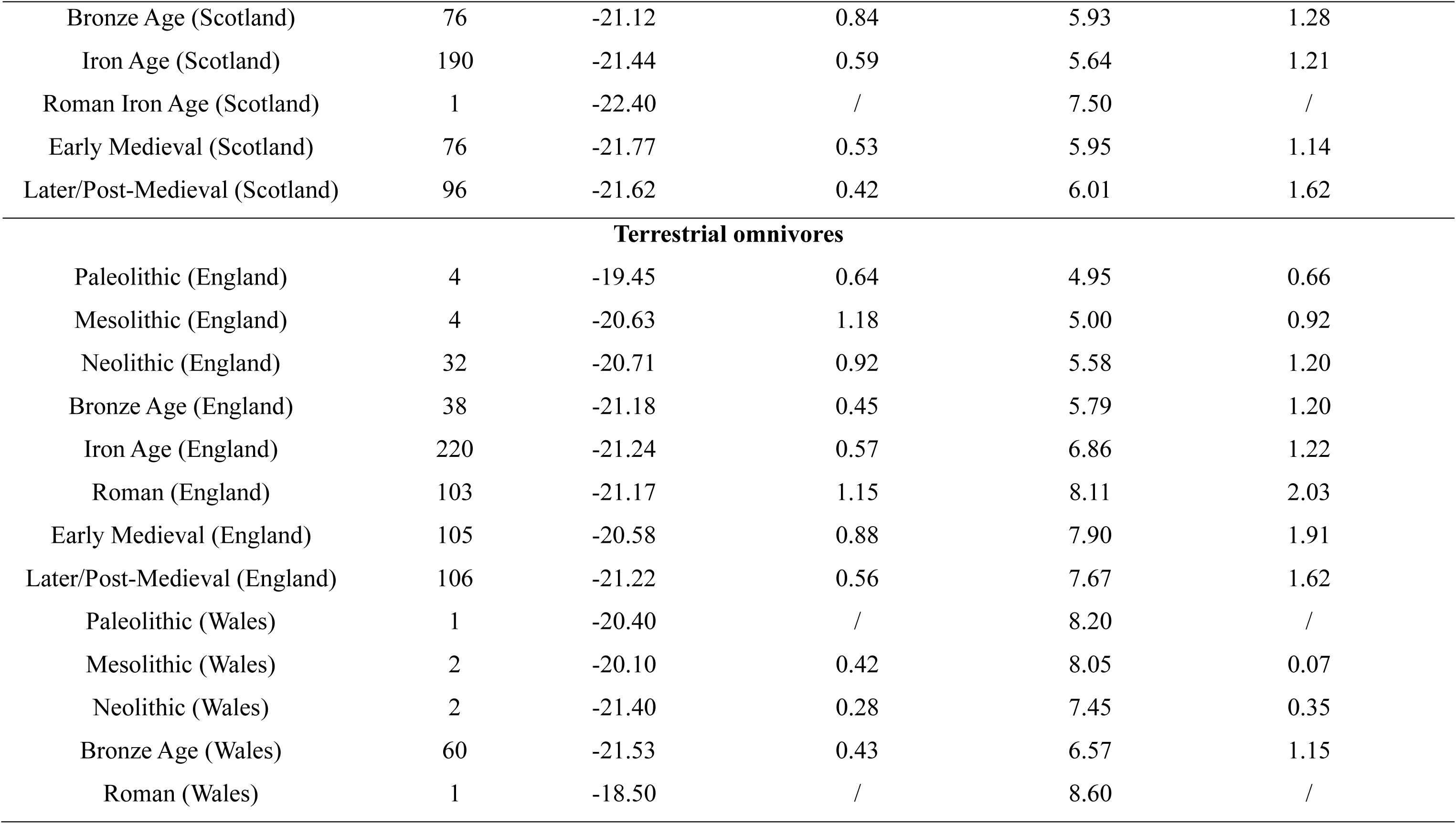
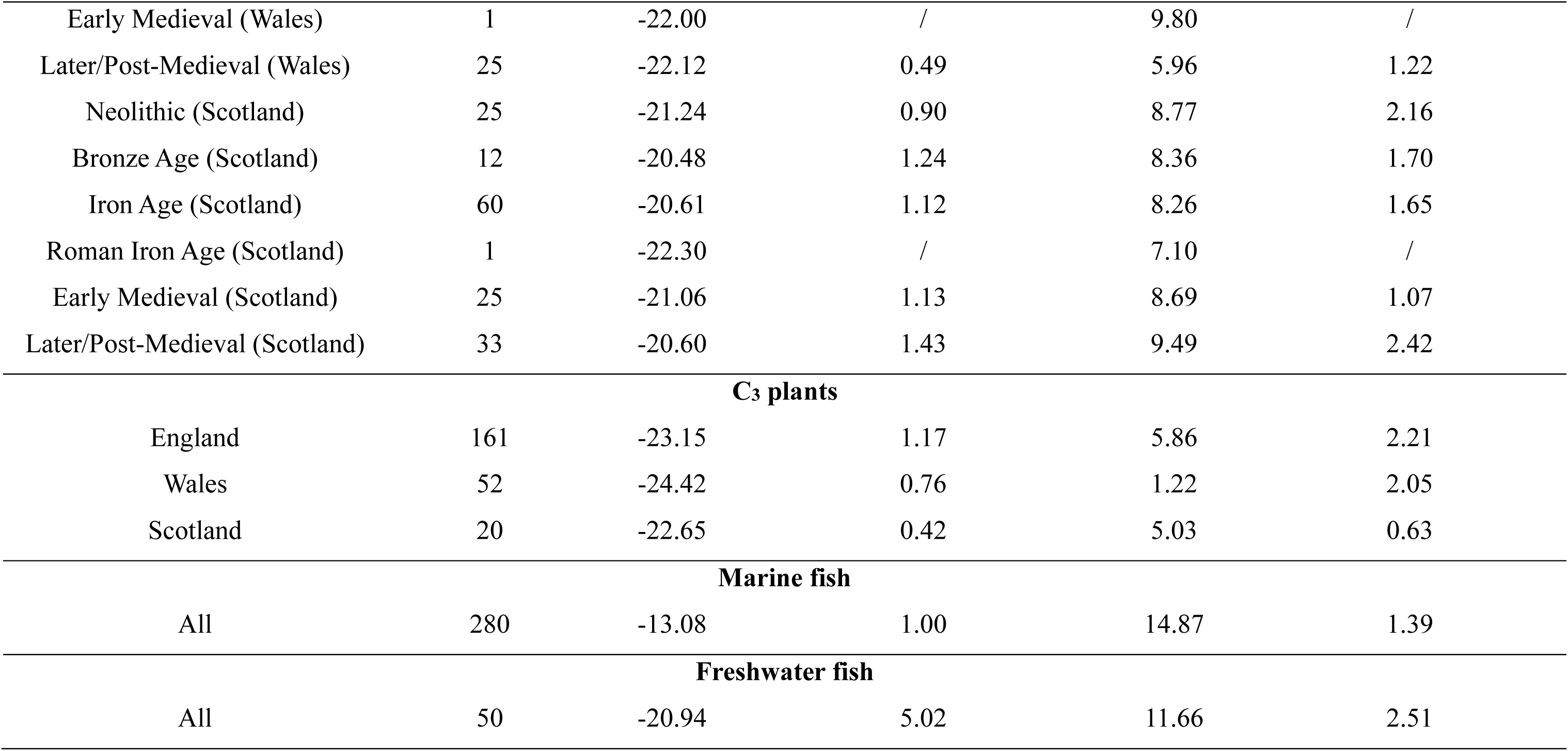
Summary statistics for recommended accessible food-resource isotope baselines in Britain.

The analyses for the remaining three food categories are considerably simpler due to their very limited sample sizes. For plants, our comparisons indicate that isotopic values vary significantly among regions, even within the same period (Fig. 59b and c). When holding region constant and varying time, isotopic variation is not pronounced, either showing no statistically significant differences or, where significant differences are detected, they are likely attributable to the sample size limitation (Fig. 57 and Fig. 58). Based on this analysis, we classified the plant groups according to region only (i.e., England, Wales, Scotland), irrespective of period (Table 1). C_3_ plants from a given region can be used as accessible food resources for human individuals from the same region. The atemporal classification is well supported by scientific evidence, particularly with regard to δ^13^C. Ice-core records indicate that the δ^13^C values of C_3_ plants remained remarkably stable from 10,000 years BP until the onset of the industrial era (∼1900 AD)[100]. After 1900, δ^13^C values increased sharply. However, this period falls outside the scope of ancient diet reconstruction. The case of δ^15^N is more complex. The δ^15^N values of plants systematically decrease with increasing mean annual precipitation (MAP) and decreasing mean annual temperature (MAT)[101,102]. These factors undoubtedly vary geographically, yet there is no reason to believe that they have remained unchanged over time. In addition, plant δ^15^N values are affected by various factors, including the sea-spray effect and anthropogenic fertilization. Thus, our region-specific baseline constructions are unlikely to accurately represent the δ^15^N values of the C_3_ plants that were actually available to ancient communities. While both our plant data and previous herbivore/omnivore isotopic comparisons suggest that temporal variation within a site has little impact on isotopic values, caution is still warranted.

The analysis of marine fish is more straightforward, as no clear isotopic differences are observed between groups, whether controlling for time while varying region or controlling for region while varying time (excluding the effects of sample size). Samples from the combined group with a large sample size (Later Medieval southern England) form an isotopic cluster that almost encompasses all other combined groups (Fig. 60). This may be attributed to the fact that marine environments change far more slowly than terrestrial environments[103], resulting in less difference over time. The fact that the marine environment forms one extremely large, interconnected system also means that the δ^13^C and δ^15^N values of marine fish typically show little geographical variation[104–106]. For example, the overall isotopic variability of particulate organic matter (POM) for the Southern California Bight was relatively low, with mean (±SD) δ^13^C and δ^15^N values of −22.7 ± 2.0‰ and 8.0 ± 1.5‰, respectively[107]. Based on these results, marine fish are classified as a single group to be used consistently as a source input for all ancient human samples (Table 1).

The results for freshwater fish are more nuanced. While the isotopic values from the largest combined group almost cover the distributions of all other groups, the total sample size is still too limited to draw firm conclusions (Fig. 64). For qualitative isotopic analysis, freshwater fish values may be of limited importance: it is sufficient to note that human δ^13^C values are not markedly elevated while δ^15^N values are distinctly higher, and to consider either the site’s proximity to rivers or the presence of fish bones as evidence for freshwater fish consumption. For quantitative dietary modeling, however, there appears to be only one feasible approach: to pool all available freshwater fish samples and use them collectively as input for modeling the diet of different human populations. Many freshwater fish migrate over considerable distances within a year[108], resulting in wide isotopic ranges even within a single site (Fig. 64). Although using a single freshwater fish database for all individuals is problematic, it remains the only viable option given the limited data and the intrinsic isotopic variability of freshwater fish (Table 1).

### Conclusion

Through our analysis, we establish a detailed classification scheme for defining accessible food resources across five food-resource categories. For terrestrial herbivores and omnivores, accessible food resources are best defined at the level of region–period combined groups. Plant data vary significantly across regions but show no marked temporal changes. It is therefore more appropriate to organize the plant data by England, Wales, and Scotland, and to apply these data to the dietary reconstruction of humans from the corresponding region. Marine fish do not appear to exhibit strong temporal or spatial isotopic variation. Accordingly, all marine fish data can be pooled and applied collectively to dietary reconstructions of all ancient human individuals. Freshwater fish follow a similar pattern to marine fish, but the limited sample size makes this conclusion less robust. The corresponding isotope baselines (δ^13^C and δ^15^N), including mean values and standard deviations, for each accessible food-resource group across the five food-resource categories are shown in Table 1. This approach not only addresses the challenge that some human individuals lack accessible food-resource data, but also helps alleviate estimation bias that may arise from small sample sizes when food resources from a single site or a few sites are used as accessible food resources.

This classification should be viewed as an attempt to maximize the use of the limited isotopic data currently available. Future studies will yield more robust conclusions as additional isotopic data on food resources become available. For terrestrial herbivores, future food-resource isotope sampling should focus on Mesolithic England; Paleolithic, Mesolithic, Neolithic, and Roman Wales; and Roman Iron Age Scotland. For terrestrial omnivores, future sampling should prioritize Paleolithic and Mesolithic England; Paleolithic, Mesolithic, Neolithic, Roman, and Early Medieval Wales; and Roman Iron Age Scotland. These region–period combined groups currently have very small sample sizes, making their use to construct isotope baselines relatively less robust. Future increases in sample size for each group would therefore substantially improve the reliability of the corresponding isotope baselines. The overall sample sizes for C_3_ plants, marine fish, and freshwater fish are very small, making the definition of accessible food resources for these categories relatively coarse and the resulting isotope baselines less robust. Thus, future sampling of C_3_ plants, marine fish, and freshwater fish should be substantially expanded to cover broader temporal and spatial ranges. This would enable more detailed isotopic analyses of these currently underrepresented food-resource categories, allowing more refined definitions of accessible food resources for each category.

## Data Availability

All data used in this study are provided in Supplementary Data 1.

## Acknowledgements

We gratefully acknowledge the financial support provided by the Durham University–China Scholarship Council PhD Scholarship. We are very grateful to Dr. Max Price for his helpful comments and suggestions on this manuscript.

## Author contributions

Z.C. conceptualized the study, collected and analyzed the data, and wrote the original manuscript. A.M. and E.F.-D. reviewed the manuscript and provided valuable comments and critical feedback.

